# Ubiquitylation by Rab40b/Cul5 regulates Rap2 localization and activity during cell migration

**DOI:** 10.1101/2021.12.06.471477

**Authors:** Emily D. Duncan, Ke-Jun Han, Margaret A. Trout, Rytis Prekeris

## Abstract

Cell migration is a complex process that involves coordinated changes in membrane transport, actin cytoskeleton dynamics, and extracellular matrix remodeling. Ras-like small monomeric GTPases, such as Rap2, play a key role in regulating actin cytoskeleton dynamics and cell adhesions. However, how Rap2 function, localization, and activation are regulated during cell migration is not fully understood. We previously identified the small GTPase Rab40b as a regulator of breast cancer cell migration. Rab40b contains a Suppressor of Cytokine Signaling (SOCS) box, which facilitates binding to Cullin5, a known E3 Ubiquitin Ligase component responsible for protein ubiquitylation. In this study, we show that the Rab40b/Cullin5 complex ubiquitylates Rap2. Importantly, we demonstrate that ubiquitylation regulates Rap2 activation, as well as recycling of Rap2 from the endolysosomal compartment to the lamellipodia of migrating breast cancer cells. Based on these data, we propose that Rab40b/Cullin5 ubiquitylates and regulates Rap2-dependent actin dynamics at the leading-edge, a process that is required for breast cancer cell migration and invasion.

**SUMMARY:** The Rab40b/Cul5 complex is an emerging pro-migratory molecular machine. Duncan et al. identify the small GTPase Rap2 as a substrate of the Rab40b/Cul5 complex. They provide evidence that Rab40b/Cul5 ubiquitylates Rap2 to regulate its localization and activity during breast cancer cell migration, ultimately proposing a model by which Rap2 is targeted to the leading-edge plasma membrane to regulate actin dynamics during cell migration.

## INTRODUCTION

Cell migration is essential for many normal biological processes including development, wound healing, and the immune response (Franz, Jones, and Ridley 2002; Vicente-Manzanares 2005). On the other hand, it is also critical for the progression of cancer metastasis, where dissemination of cancer cells from the primary tumor to a distant site is the leading cause of mortality in patients (Bravo-Cordero, Hodgson, and Condeelis 2012). It is well accepted that cell migration requires coordinated changes in membrane trafficking, the actin cytoskeleton, adhesion dynamics, and the targeted secretion of matrix metalloproteinases (MMPs) (Ridley et al. 2003; Pollard and Borisy 2003; Parsons, Horwitz, and Schwartz 2010; Murphy and Courtneidge 2011; Abitha Jacob and Prekeris 2015; Warner, Wilson, and Caswell 2019). However, the molecular machinery that governs these processes is not fully understood and is a main focus of current studies in the field.

Small monomeric GTPases are tightly regulated molecular switches, cycling between an “active” GTP state and an “inactive” GDP state in order to facilitate proper vesicular transport and protein recruitment to distinct subcellular locations (Reiner and Lundquist 2018). Members of the Ras GTPase superfamily have been shown to play essential roles in cell migration and invasion by coordinating the dynamics of many events described above (Wennerberg, Rossman, and Der 2005; Sadok and Marshall 2014; Lawson and Ridley 2018; Gimple and Wang 2019). Though the Rho family is arguably the most studied in the context of cell migration, it is increasingly clear that other members of the Ras superfamily are equally important for this process. One example is the Rap subfamily (containing Rap1a, 1b and Rap2a, 2b, 2c), which are the most closely related to Ras itself (more than 50% overall identity) compared to the Rho, Rab, Ran, and Arf families (Wennerberg, Rossman, and Der 2005). Rap1, originally described as a suppressor of Ras-mediated transformation, is now widely accepted as a key regulator of integrin-mediated cell adhesion and actin reorganization (Kahana and Gottschling 1999; Bivona et al. 2004; Bos 2005; Jeon et al. 2007; Kooistra, Dubé, and Bos 2007; Boettner and Van Aelst 2009). Recent reports have shown that Rap1 can activate MAP kinase signaling cascades similar to Ras, and has clear implications in the pathogenesis of cancer (Stork 2003). Additionally, work in *Dictyostelium discoideum*, *Drosophila*, and other systems has provided important insight into the regulation of Rap1 (Rebstein et al. 1997; Asha et al. 1999; Jeon et al. 2007; C. Liu et al. 2010; Chang et al. 2018).

Though Rap1 and Rap2 share ∼60% identity, much less is known about the function of Rap2 GTPases. What we do know is that Rap1 and Rap2 do have, to some extent, overlapping functions as well as several shared effector proteins (Nancy et al. 1999).

One of the earliest reported functions of Rap2 was the promotion of B cell migration via integrin-mediated adhesions (McLeod et al. 2002, 2004). This, combined with another initial finding suggesting Rap2 is critical for *Xenopus* gastrulation and development, provided early evidence that the Rap2 family may play a crucial role during cell migration across diverse biological systems (Choi and Han 2005). Over the last decade, the field has slowly emerged, where several studies have linked Rap2 to cell polarity, actin cytoskeleton regulation, cancer cell invasion, and most recently, coordination of mechanosensing and Hippo signaling (Taira et al. 2004; Gloerich et al. 2012; Bruurs and Bos 2014; J. Di et al. 2015; Meng et al. 2018). However, we still do not fully understand the main function of Rap2 during cell migration, nor the regulation of its subcellular localization or activation. Determining whether modulation of Rap2 trafficking, localization, and activation is necessary to drive cell migration is the main focus of this study.

We previously identified another small monomeric GTPase Rab40b as an important regulator of 3D ECM remodeling, specifically during breast cancer cell migration (Jacob et al. 2013; Jacob et al. 2016). The Rab subfamily of proteins comprise the largest class of small monomeric GTPases, functioning as master regulators of intracellular membrane traffic (Stenmark 2009). While all Rabs share a conserved globular G-domain (housing the Switch I and Switch II regions), Rab40 GTPases have a unique extended C-terminal domain that contains the conserved SOCS (Suppressor of Cytokine Signaling) box (Coppola et al. 2019). This ∼40 amino acid motif facilitates binding to Cullin5 (Cul5) (Kile et al. 2002; Kamura et al. 2004; Lee et al. 2007; Dart et al. 2015; Yatsu et al. 2015; Linklater et al. 2021; Duncan et al. 2021). Together, this Rab40/Cul5 module, along with RING-box protein Rbx2 and adaptor proteins Elongin B and Elongin C, make up the larger Cullin-RING Ligase (CRL5) complex, which canonically regulates protein ubiquitylation (Kile et al. 2002; Petroski and Deshaies 2005; Linossi and Nicholson 2012; Okumura et al. 2016; Linklater et al. 2021).

Here, we demonstrate that Rap2 is a substrate of the Rab40b/Cul5 complex and that ubiquitylation of selective Rap2 residues plays a major role in regulating its function. Specifically, we find that Rab40b/Cul5-dependent ubiquitylation regulates targeting of Rap2 to the leading-edge plasma membrane of migrating cells. We demonstrate that inhibition of Rap2 ubiquitylation blocks Rap2 endosome-to-plasma membrane recycling, leading to rapid lysosomal degradation, and termination of Rap2 signaling. We also show that Rap2 ubiquitylation is required for its activation. Ultimately, we offer evidence for why proper localization and activation of Rap2 is critical for regulating actin dynamics at the leading-edge and promoting cell migration and how this process is regulated by Rab40b/Cul5. Based on our combined data, we propose a model in which Rab40b is a dual-functioning Rab GTPase, given its co-regulation of vesicular MMP trafficking as well as Rap2 spatiotemporal dynamics, and such co-regulation plays a key role in driving cell migration.

## RESULTS

### Rap2 is required for breast cancer cell migration and invasion

The Rap subfamily of small GTPases (Rap1a, b, and Rap2a, b, c) belong to the Ras protein superfamily and have been implicated in a variety of biological processes including signal transduction, cell migration, and cell adhesion (Itoh et al. 2007; McLeod et al. 2004, 2002). Since their discovery, the Rap1 GTPases have elicited the most interest, specifically for their role in regulating integrin-mediated cell adhesion (Bos et al. 2003; Bos 2005; Boettner and Van Aelst 2009). Though suspected to be involved in the pathogenesis of cancer, the exact function of the Rap2 family remains elusive and existing reports are conflicting. To gain a clearer understanding of Rap2’s role in our breast cancer model system, we made a triple CRISPR/Cas9 knockout of all three Rap2 isoforms in MDA-MB-231 cells (denoted as Rap2 KO), which to our knowledge is the first triple KO developed in this system. We generated two knockout lines (Rap2 KO1 and Rap2 KO2) to diminish the possibility of off-target effects. Total loss of Rap2 protein was confirmed via western blot using a pan-Rap antibody (Figure 1A) and genotyping (see methods).

**Figure 1.**
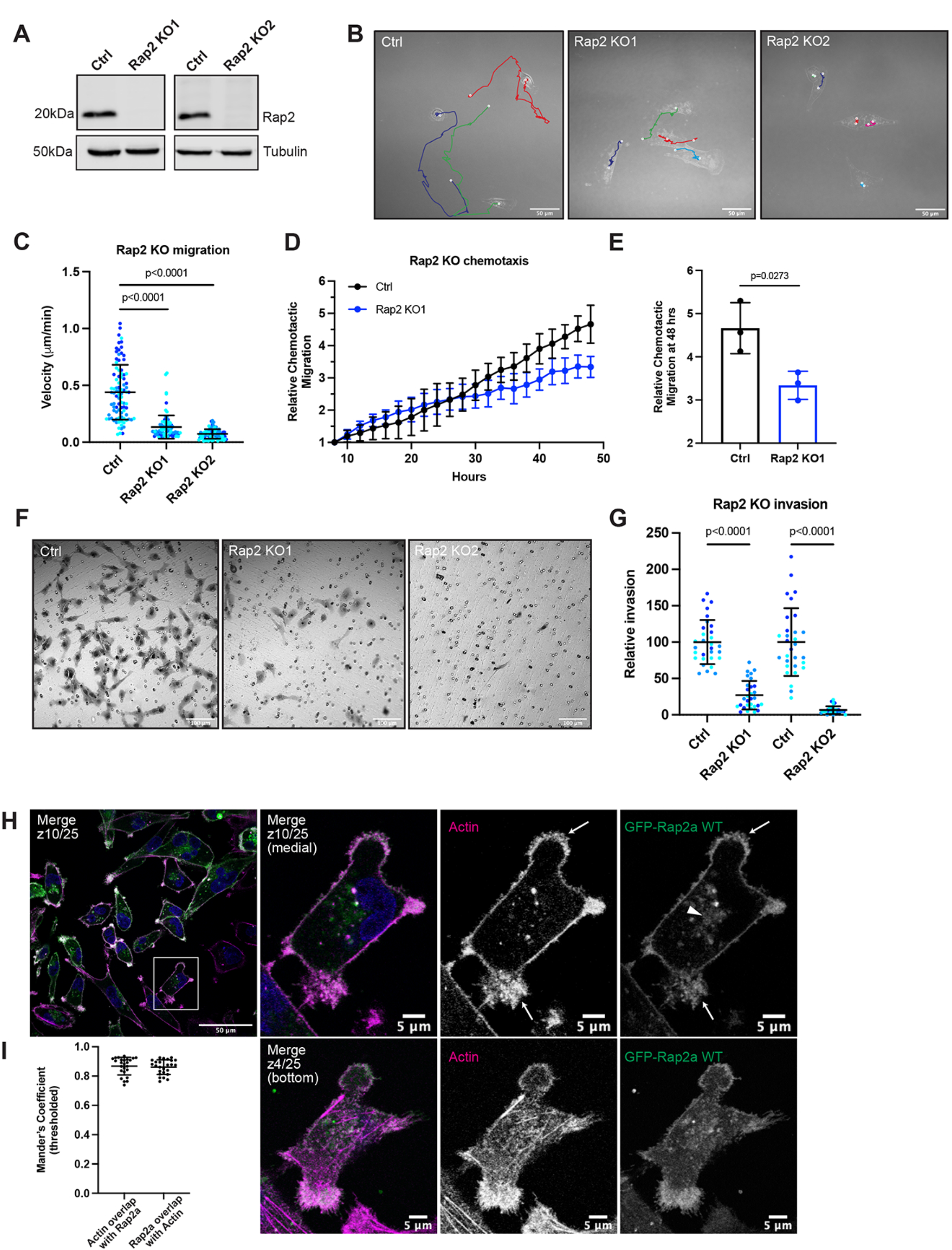
Rap2 is necessary for MDA-MB-231 migration and invasion. (A) Loss of Rap2 confirmed via western blot. Two different triple CRISPR knockouts were made in MDA-MB-231s (see methods). All “Ctrl” cells shown in this figure are dox-inducible Cas9 MDA-MB-231s that were used to generate CRISPR lines. 50µg of lysate was loaded for each sample. (B) Time-lapse 2D migration. Ctrl and Rap2 KO cells were plated on collagen-coated glass dishes and imaged every 10 mins for 16hrs using a brightfield 40x objective. Representative still images show the last frame of the time-lapse experiment, where the cell body indicates the location of the cell at the last time-point. Cells were manually tracked in Fiji using the “Manual Tracking” plugin. Colors denote individual cell tracks over 16hrs. White dots represent the start and finish of each cell track. Scale bars = 50µm. (C) 2D migration velocity quantification. From the Fiji manual tracking, velocity data (µm/min) was extracted for each individual cell. Three biological replicates were performed for each cell line. For each experiment, 30 cells (color coded) were randomly chosen for velocity tracking (∼3 cells from ∼10 fields of view). Each cell was treated as its own data point (n=90). One-way ANOVA with Tukey’s multiple comparisons test. Mean ± SD. Ctrl vs Rap2 KO1 p<0.0001. Ctrl vs Rap2 KO2 p<0.0001. (D) Chemotactic cell migration Ctrl vs Rap2 KO1 cells. Briefly, cells were resuspended in serum-starved media and plated in an IncuCyte ClearView 96-Well Plate with chemoattractant (full media) in the bottom chamber. Cells were imaged every 2hrs for 48hrs using an IncuCyte S3 instrument. Raw data (the sum area of all migrated cells normalized to the area at time zero) was extracted and averaged for each technical replicate. Data was then adjusted/normalized to the 8hr timepoint (see methods). The graph shows “Relative Chemotactic Migration” over the 48hr time course. Mean ± SD. (E) Chemotactic migration quantification. Three biological replicates were performed, with six technical replicates in each experiment. Statistical analysis (unpaired t-test) was performed on the Relative Chemotactic Migration at 48hrs. Mean ± SD at 48hrs. Ctrl vs Rap2 KO1 p=0.0273. (F) Boyden chamber invasion assay. Ctrl and Rap2 KO cells were plated in a modified Boyden chamber coated with Matrigel. Cells were allowed 20hrs to invade through the Matrigel coated pores before fixation and crystal violet staining. Inserts were imaged using a brightfield 20x air objective. Representative images are shown. Hollow circles indicate the 8µm pores. Scale bars = 100µm. (G) Invasion assay quantification. Three biological replicates were performed, with technical duplicates in each experiment. 5 fields of view were imaged for each Matrigel insert (resulting in 10 fields of view per experiment per condition, color coded). Raw number of cells invaded per field of view were normalized to Ctrl. Rap2 KO1 and Rap2 KO2 cells were analyzed at different times, hence two Ctrl samples. Each field of view was treated as its own data point (n=30). One-way ANOVA with Tukey’s multiple comparisons test. Mean ± SD. Ctrl vs Rap2 KO1 p<0.0001. Ctrl vs Rap2 KO2 p<0.0001. (H) GFP-Rap2a WT localization. MDA-MB-231 cells stably expressing GFP-Rap2a were fixed and stained with Phalloidin (magenta) and DAPI (blue). Z-slices are indicated on merged images. In most images, medial z-slices are shown. Here, we also provide a z-slice at the bottom of the cell, to highlight overlap between Actin and GFP-Rap2a. Arrows point to example sites of Actin and GFP-Rap2a co-localization. The arrowhead indicates the GFP-Rap2a intracellular organelle population. Scale bars = 50µm, 5µm. (I) Co-localization analysis of Actin and GFP-Rap2a. Thresholded Mander’s coefficients were calculated using the Fiji Coloc 2 plugin. One biological replicate was performed, with 5 fields of view and 5 cells in each field (n=25). The fraction of Actin overlapping with Rap2a (tM1, mean=0.8678) and the fraction of Rap2a overlapping with Actin (tM2, mean=0.8612) are shown. Mean ± SD.

We first asked whether loss of Rap2 leads to any global 2D migration defects, using live-cell imaging. Cells were plated on collagen-coated glass dishes, allowed to adhere overnight, and imaged every 10 minutes for 16 hours using a brightfield objective (Videos 1-3). Compared to control cells, Rap2 KO cells exhibit decreased individual cell migration as indicated by significantly lower velocity (Figure 1B & C). We next asked whether Rap2 KO cells have defects in chemotactic cell migration. Control and Rap2 KO cells were plated in an IncuCyte ClearView 96-well Chemotaxis Plate with serum-starved media in the top chamber and full MDA-MB-231 media (chemoattractant) in the bottom chamber. Cells were allowed to migrate toward the chemoattractant for 48 hours, with images taken every 2 hours. As expected, based on the individual cell migration analysis, Rap2 KO cells show decreased chemotactic migration compared to control cells (Figure 1D & E), although, we cannot rule out whether this chemotactic migration defect is due to lack of signal sensing or the inherent inability of these cells to move. Interestingly, while the Rap2 KO individual cell velocity defect in Figure 1C is quite dramatic, we only see ∼50% inhibition of chemotactic migration at the 48-hour timepoint. This may point at differences in Rap2’s function during individual vs collective cell migration. Nonetheless, these results in sum demonstrate that Rap2 modulates 2D breast cancer cell migration.

We next tested the invasive capability of MDA-MB-231 Rap2 KO cells. To that end, control and Rap2 KO cells were plated in a modified Boyden Chamber pre-coated with Matrigel™ and given 20 hours to invade through the Matrigel™ coated pores before fixation and imaging. Rap2 KO cells show a striking decrease in their ability to invade through Matrigel™ compared to control cells (Figure 1F & G). Taken together, our collective data demonstrate that Rap2 is necessary for both 2D and 3D migration and invasion in MDA-MB-231s. These data provide strong evidence for the Rap2 subfamily being pro-migratory in breast cancer cells. Additionally, these striking phenotypes convey a clear importance for studying the function and regulation of Rap2 during cell migration.

### Rap2 cycles between the lamellipodia plasma membrane and endosomes during breast cancer cell migration

To start dissecting how Rap2 may regulate breast cancer cell migration and invasion, we first examined subcellular localization of Rap2 in MDA-MB-231s, as this has not yet been fully characterized. In humans, the Rap2 family encompasses three closely related paralogs—Rap2a, Rap2b, and Rap2c, which have over 90% sequence identity. All of the evidence so far suggests that all three Rap2 isoforms have similar functions, thus, for most of our studies we have focused on the Rap2a isoform. Our attempts to visualize endogenous Rap2 in cells with the pan-Rap antibody were unsuccessful, so instead we generated an MDA-MB-231 cell line stably expressing GFP-Rap2a (Supplementary Figure 1C) and analyzed its subcellular distribution using immunofluorescence microscopy. As shown in Figure 1H, GFP-Rap2a localizes predominantly to the plasma membrane, where it is enriched at lamellipodia ruffles.

Importantly, plasma membrane bound Rap2a is clearly co-localized with the actin cytoskeleton (Figure 1I), consistent with the putative role of Rap2 in regulating actin cytoskeleton dynamics.

In addition to the plasma membrane population, some of GFP-Rap2a is also observed within intracellular organelles (Figure 1H). Since some of the Ras subfamily members, namely HRas and NRas have been proposed to signal from the Golgi (Hancock 2003), we wondered about the identity of these Rap2a-containing intracellular organelles. To this end, we co-stained GFP-Rap2a cells with known endocytic pathway markers including GM130 (Golgi), EEA1 (early endosomes), CD63 (late endosomes/lysosomes), and Syntaxin13 (Rab4 and Rab11 recycling endosomes).

While we do observe some Rap2a at the Golgi, consistent with preliminary reports (Pizon et al. 1994), most intracellular GFP-Rap2a is clearly present within the endolysosomal compartment, where it co-localizes with both EEA1-positive early endosomes and CD63-positive late endosomes/lysosomes (Figure 2A-C, E). Consistent with at least some of the Rap2 pool being trafficked to lysosomes, we observe an increase in Rap2 co-localization with CD63 when cells are treated with Bafilomycin A1, an inhibitor of lysosomal degradation (Figure 2F). Finally, we note that GFP-Rap2a is rarely found in the cytosolic pool. This is supported by cell fractionation analysis, where we observe no detectable endogenous Rap2 in the cytosol fraction and robust signal in the membrane fraction, suggesting that Rap2 is predominantly membrane bound in the cell (Figure 2G).

**Figure 2.**
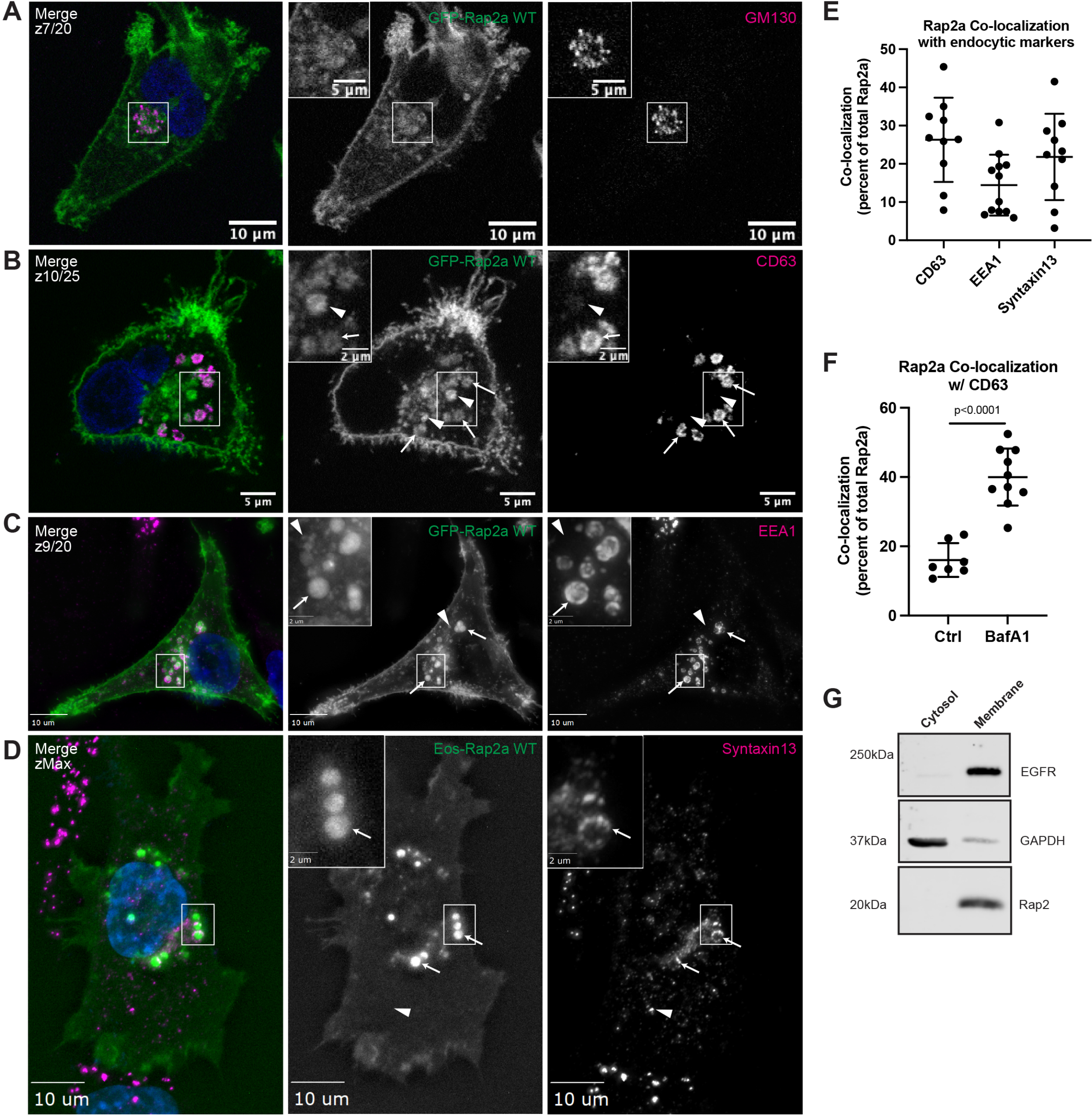
Rap2 localizes to the plasma membrane and endolysosomal compartment in MDA-MB-231s. (A) Co-localization of GFP-Rap2a and GM130. MDA-MB-231 cells stably expressing GFP-Rap2a were fixed and stained with the Golgi marker GM130 (magenta) and DAPI (blue). Inset shows GFP-Rap2a and GM130 overlap. Scale bars = 10µm, 5µm. (B) Co-localization of GFP-Rap2a and CD63. MDA-MB-231 cells stably expressing GFP-Rap2a were fixed and stained with the lysosomal marker CD63 (magenta) and DAPI (blue). Arrows indicate examples of GFP-Rap2a and CD63 overlap. Arrowheads point to GFP-Rap2a organelles that are not CD63 positive. Scale bars = 5µm, 2µm. (C) Co-localization of GFP-Rap2a and EEA1. MDA-MB-231 cells stably expressing GFP-Rap2a were fixed and stained with the early endosome marker EEA1 (magenta) and DAPI (blue). Arrows indicate examples of GFP-Rap2a and EEA1 overlap. Arrowheads point to GFP-Rap2a organelles that are not EEA1 positive. Widefield microscope. Scale bars = 10µm, 2µm. (D) Co-localization of Eos-Rap2a and Syntaxin13. MDA-MB-231 cells stably expressing Eos-Rap2a were fixed and stained with the recycling endosome marker Syntaxin13 (magenta) and DAPI (blue). Arrows indicate examples of Eos-Rap2a and Syntaxin13 overlap. Arrowhead points to Syntaxin13 endosome that is not Rap2a positive. Widefield microscope. Scale bars = 10µm, 2µm. (E) Rap2a co-localization analysis with endolysosomal compartments. 3i SlideBook6 software was used to calculate the percent of total GFP-Rap2a that co-localizes with the markers indicated (see methods). Two biological replicates were performed, with roughly 5 cells imaged for each replicate. Mean ± SD. n=10 for GFP-Rap2a/CD63, n=12 for GFP-Rap2a/EEA1, n=10 for Eos-Rap2a/Syntaxin13. (F) Rap2a and CD63 co-localization analysis in Ctrl cells vs cells treated with lysosomal inhibitor Bafilomycin A1. MDA-MB-231 cells stably expressing GFP-Rap2a were treated with either DMSO (Ctrl) or 200nM Bafilomycin A1 for 16 hrs. Cells were then fixed and stained for CD63, following co-localization analysis. 3i SlideBook6 software was used to calculate the percent of total GFP-Rap2a that co-localizes with CD63 (see methods). One biological replicate was performed. Mean ± SD. n=7 cells for Ctrl, n=10 cells for BafA1. Unpaired t-test. p=<0.0001. (G) Cell fractionation analysis cytosol vs membrane. MDA-MB-231 parental cells were fractionated (see methods) to determine subcellular distribution of endogenous Rap2. Cytosol and membrane fractions were collected, separated by SDS-PAGE, and western blotted against known cytosol (GAPDH) and membrane (EGFR) markers. 30µg of sample was loaded for each fraction.

The co-localization with Syntaxin13 suggests that Rap2 may recycle from early endosomes, likely via Rab4 and/or Rab11 recycling endosomes (Figure 2D & E) (Zerial and McBride 2001). To investigate this further, we co-transfected MDA-MB-231 cells with mCherry-Rap2a and either YFP-Rab4 or YFP-Rab11a to visualize potential co-localization at recycling endosomes (Figure 3A & B). Indeed, we do detect Rap2-Rab4 and Rap2-Rab11 positive organelles, which supports a model in which Rap2 gets recycled to the lamellipodia plasma membrane via the Rab4 and Rab11 pathways (Figure 3C). We note that the co-localization is not as robust as other endocytic markers like EEA1 and CD63, which again is consistent with Rap2 being quickly recycled via the Rab4 and Rab11 pathways, thus, at steady state only a small portion of Rap2 is present within recycling endosome compartments.

**Figure 3.**
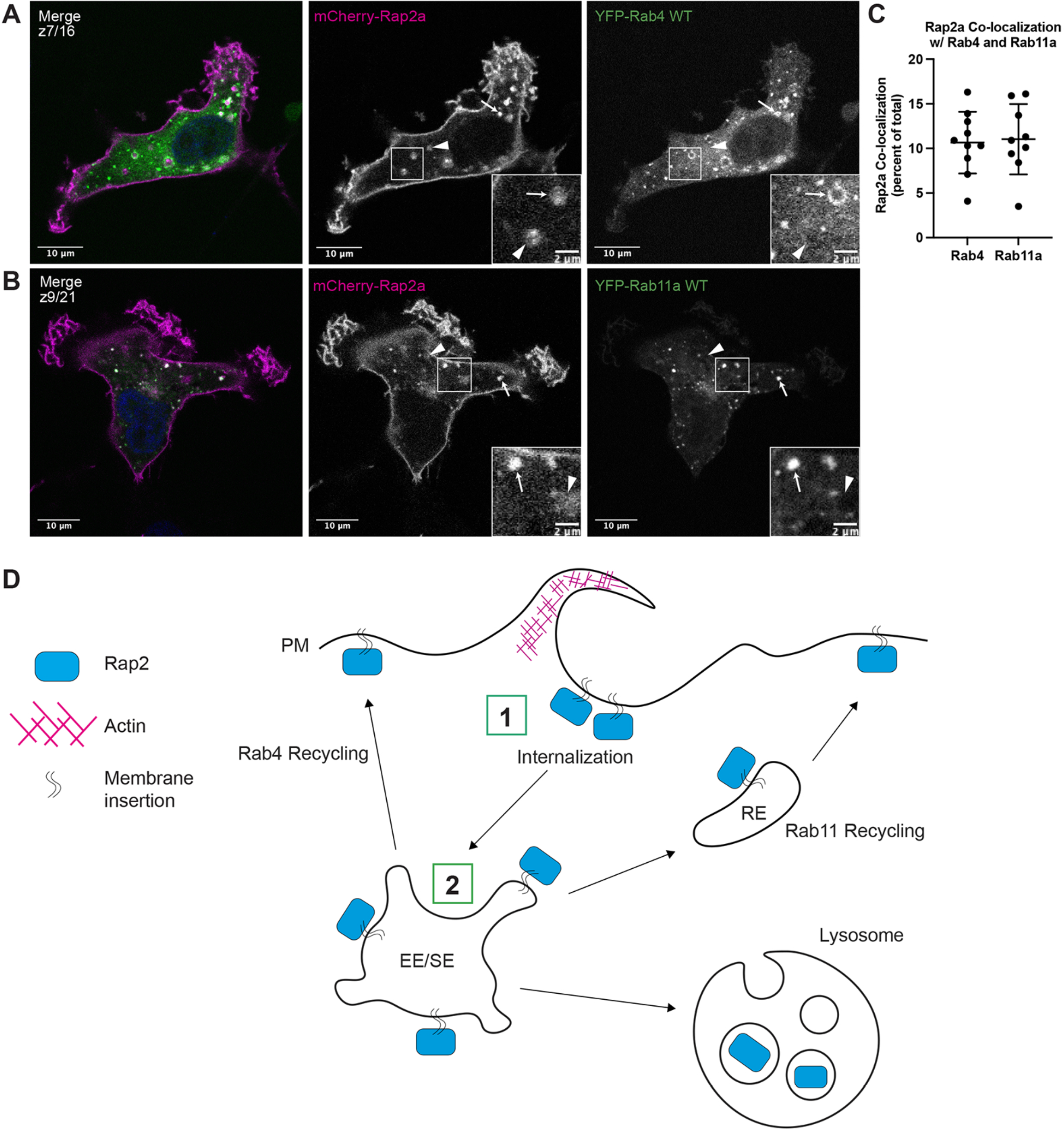
Rap2 is dynamically trafficked through the endocytic pathway. (A) Co-localization of mCherry-Rap2a and YFP-Rab4 WT. MDA-MB-231 cells were transiently co-transfected with mCherry-Rap2a and YFP-Rab4 WT, followed by fixation and staining for DAPI (blue). Arrows indicate examples of mCherry-Rap2a and YFP-Rab4 WT overlap. Arrowheads point to mCherry-Rap2a organelles that are not Rab4 positive. Scale bars = 10µm, 2µm. (B) Co-localization of mCherry-Rap2a and YFP-Rab11a WT. MDA-MB-231 cells were transiently co-transfected with mCherry-Rap2a and YFP-Rab11a WT, followed by fixation and staining for DAPI (blue). Arrows indicate examples of mCherry-Rap2a and YFP-Rab11a WT overlap. Arrowheads point to mCherry-Rap2a organelles that are not Rab11a positive. Scale bars = 10µm, 2µm. (C) Rap2a co-localization analysis with Rab4 WT and Rab11a WT. 3i SlideBook6 software was used to calculate the percent of total mCherry-Rap2a that co-localizes with Rab4 or Rab11a (see methods). One biological replicate was performed. Mean ± SD. n=10 for mCherry-Rap2a/YFP-Rab4 WT, n=9 for mCherry-Rap2a/YFP-Rab11a WT. (D) Model for Rap2 endocytic trafficking. 1. Rap2 is internalized at the lamellipodia plasma membrane via a pinocytosis-like mechanism 2. Internalization mediates the delivery of Rap2 to EEA1/Rab5-positive early endosomes where it is sorted into different fates, either recycling to the plasma membrane via the Rab4/Rab11 pathways or targeting to lysosome for degradation. We hypothesize that Rap2 recycling back to the leading-edge is needed to sustain cell migration.

To further test whether Rap2 recycles via the Rab4 and Rab11 pathways, we co-transfected MDA-MB-231 cells with mCherry-Rap2a and either YFP-Rab4-S27N (dominant-negative mutant) or GFP-FIP5-RBD (C-terminal fragment of Rab11-FIP5 that inhibits Rab11-mediated recycling pathway (Peden et al. 2004; Willenborg et al. 2011)). Expression of either mutant delays the exit of recycling cargo from endosomes, thus, accumulating cargo proteins in either YFP-Rab4-S27N or GFP-FIP5-RBD positive organelles. Consistent with our hypothesis, we observe mCherry-Rap2a present in both YFP-Rab4-S27N (affects Rab4-dependent recycling) and GFP-FIP5-RBD (affects Rab11-dependent recycling) organelles (Supplementary Figure 1A & B). In sum, our data suggests that Rap2 is dynamically trafficked through the endocytic pathway and is likely recycled through early endosomes/recycling pathway back to the leading-edge plasma membrane (Figure 3D, trafficking itinerary model for Rap2).

To further define the spatiotemporal properties of Rap2 localization in migrating cells, we next performed time-lapse analysis of GFP-Rap2a-expressing MDA-MB-231 cells. As shown in the Figure 4A, GFP-Rap2a is enriched at ruffling lamellipodia, consistent with its proposed role in regulating actin dynamics during cell migration. It is especially clear using live imaging that Rap2 is more dynamic at the leading-edge plasma membrane compared to the intracellular membrane bound population.

**Figure 4.**
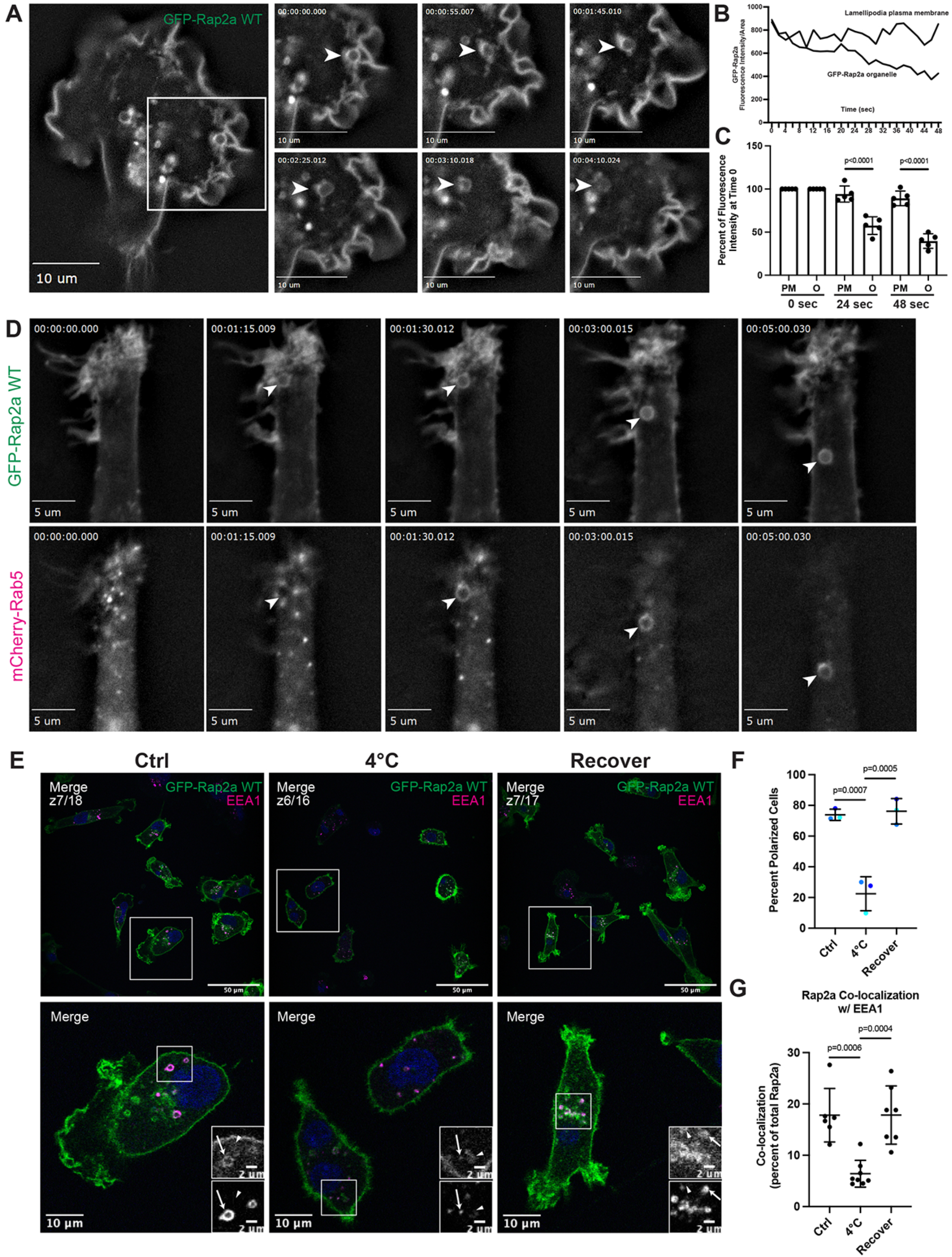
Rap2 is enriched at the lamellipodia leading-edge where its internalization and recycling to the plasma membrane is important for cell migration. (A) Live imaging of GFP-Rap2a in MDA-MB-231 cells. Cells stably expressing GFP-Rap2a were plated on a collagen-coated glass dish and imaged every 5 seconds using a 63x objective. Widefield microscope. Still images are shown. Arrowheads point to an example of GFP-Rap2a internalization over time from the plasma membrane in macropinosome-like vesicles. We also note that GFP-Rap2a organelle decreases in fluorescence intensity over time. See Video 4. Scale bars = 10µm. (B) Line-scan analysis of internalized GFP-Rap2a organelle in (A). A circular line was drawn around the GFP-Rap2a organelle marked in Figure 4A, and fluorescence intensity/area was measured at each timepoint (every 2 seconds). The same thing was done with a line-scan at the plasma membrane. The fluorescence intensity of GFP-Rap2a around the organelle line and at the plasma membrane is plotted as fluorescence intensity/area (µ^2^) across 48 seconds. (C) Same analysis as Figure 4B, but more organelles quantified. Graph includes original organelle from Figure 4A/4B, plus four more organelles from different cells. As in Figure 4B, line-scans were draw both around the GFP-Rap2a organelles (see methods for criteria) and at the lamellipodia plasma membrane. The intensity of GFP-Rap2a in each pixel along these lines was determined with either ImageJ or 3i Slidebook imaging software. Fluorescence intensity/area (µ^2^) at 0 seconds (start of time-lapse), 24 seconds (half of time-lapse), and 48 seconds (end of time-lapse) was plotted for all 5 organelles as a % of time 0. Statistical analysis (one-way ANOVA with Tukey’s multiple comparisons test) was used to compare fluorescence intensity changes between the plasma membrane and internalized organelles. Mean ± SD. PM=Plasma membrane. O=organelle. PM vs O at 24 sec p=<0.0001. PM vs O at 48 sec p=<0.0001. (D) Live imaging of GFP-Rap2a and mCherry-Rab5 in MDA-MB-231 cells. Cells stably expressing GFP-Rap2a were transiently transfected with mCherry-Rab5, plated on a collagen-coated glass dish, and imaged every 5 seconds using a 63x objective. Widefield microscope. Still images are shown. Arrowhead points to one example of a GFP-Rap2a internalized organelle over time, which becomes mCherry-Rab5 positive soon after internalization from the plasma membrane. See Video 6. Scale bars = 5µm. (E) Endocytosis cold block experiment. Three conditions were set up using MDA-MB-231 cells stably expressing GFP-Rap2a: 1. 37°C (Ctrl) 2. 4°C for 60 minutes and 3. 4°C for 60 minutes followed by 37°C for 40 minutes (Recover). Cells were fixed and stained for EEA1 (magenta) and DAPI (blue). Scale bars = 50µm, 10µm, 2µm. Arrows indicate examples of GFP-Rap2a and EEA1 overlap. Arrowheads point to GFP-Rap2a organelles that are not EEA1 positive. (F) Percent polarized cells quantification from cold block experiment in (E). Ctrl, 4°C, and Recover cells were scored for polarized vs non-polarized cells. Briefly, polarized cells had enrichment of Rap2 at the lamellipodia (see methods). Three biological replicates were performed. 8 fields of view were taken for each n, with roughly 6 cells in each field. Graph shows percent of polarized cells for each condition (color coded). Mean ± SD. One-way ANOVA with Tukey’s multiple comparisons test. Ctrl vs 4°C p=0.0007. 4°C vs Recover p=0.0005. (G) Quantification of co-localization between GFP-Rap2a and EEA1 in Ctrl vs 4°C vs Recover cells. Cells stably expressing GFP-Rap2a were fixed and stained with EEA1. 3i SlideBook6 software was used to calculate the percent of total GFP-Rap2a that co-localizes with EEA1. One biological replicate was performed, n=6 for Ctrl cells, n=8 for 4°C cells, n=7 for Recover cells. One-way ANOVA with Tukey’s multiple comparisons test. Ctrl vs 4°C p=0.0006. 4°C vs Recover p=0.0004.

Interestingly, we also observe that during lamellipodia ruffling GFP-Rap2a gets internalized in large pinocytosis-like organelles that originate at lamellipodia and subsequently move in a retrograde direction toward the retracting end of the cell (Figure 4A, Video 4, for additional cell see Video 5). Importantly, similar pinocytosis-like internalization of leading-edge plasma membrane has been reported before and is known to be required for cell migration (Moreau et al. 2019). Further, GFP-Rap2a containing organelles can be observed decreasing in fluorescence intensity over time, (Figure 4B shows quantification for organelle depicted in Figure 4A/Video 4, Figure 4C shows quantification for 4 additional organelles). Though some of this can be attributed to whole cell bleaching, we observe a noticeable decrease in fluorescence intensity over time when comparing internalized organelles to the plasma membrane (Figure 4B & C). This hints that Rap2 may be recycled back to the plasma membrane, especially given our previous data suggesting Rap2 may be trafficked through the Rab4/Rab11 recycling pathways. If this is the case, newly internalized GFP-Rap2a organelles should co-stain with a known early endosome/sorting endosome marker such as Rab5. Indeed, when we co-transfect MDA-MB-231 cells with GFP-Rap2a and mCherry-Rab5, we observe that newly internalized GFP-Rap2a organelles become Rab5 positive soon after internalization (Figure 4D, Video 6). In sum, we propose that pinocytosis-like GFP-Rap2a internalization mediates the delivery of Rap2 to EEA1/Rab5-positive early endosomes and that Rap2 is likely recycled back to the leading-edge in order to sustain cell migration (Figure 3D).

Finally, to evaluate whether dynamic recycling of Rap2 is needed for leading-edge formation and ultimate migration, we performed a cold block experiment aimed at inhibiting endocytosis. Previous work in the membrane trafficking field has shown that incubation of cells at 4°C slows down endocytosis from the plasma membrane, without a complete block. Thus, we incubated GFP-Rap2a expressing cells at 4°C for 60 minutes following fixation and staining. Compared to control cells, we observe that 4°C treatment significantly decreases the percent of polarized cells (cells with a defined leading-edge lamellipodia containing enriched GFP-Rap2a), suggesting that rapid endocytosis and recycling may be needed to maintain Rap2 lamellipodia enrichment (Figure 4E & F). Additionally, we find that GFP-Rap2a co-localizes with EEA1 at a lower frequency when the temperature is reduced to 4°C, indicative of slower endocytosis and less Rap2 in early endosomes/sorting endosomes (Figure 4G). Importantly, both these findings can be rescued when we place 4°C treated cells back at 37°C for 40 minutes (“Recover”, Figure 4E-G). Altogether, these data support our working model that Rap2 is dynamically trafficked, where quick re-distribution and recycling to the leading-edge plasma membrane is important for driving cell migration.

### Rap2 subcellular localization and activation state are closely intertwined

Based on previous Rap1 observations, we speculated that these two populations of Rap2 (plasma membrane vs endolysosomal compartment) might be correlated with nucleotide status and GTPase activity (Ohba, Kurokawa, and Matsuda 2003; Bivona et al. 2004; Jeon et al. 2007). To test this, we generated stable cell lines expressing either constitutively active (G12V; GTP-bound) or dominant-negative forms (S17N; GDP-bound) of Rap2a (Supplementary Figure 1D) and performed localization analysis on these mutants (Feig and Cooper 1988; Gibbs et al. 1989; Feig 1999). We find that Rap2a-G12V localizes primarily to the lamellipodia plasma membrane, whereas Rap2a-S17N shifts to a more endosome/lysosome localization (Figure 5A & B). Consistent with this observation, quantification of the intracellular pool compared to the whole cell (fluorescence signal) reveals a significant difference between Rap2a-G12V and Rap2a-S17N, where the dominant-negative mutant has decreased GFP-Rap2a signal at the plasma membrane (Figure 5C, Supplementary Figure 2C). Like Rap2a-WT, Rap2a-G12V appears to traffic between early endosomes, lysosomes, and the lamellipodia plasma membrane (Supplementary Figure 1F & G). However, Rap2a-S17N loses plasma membrane enrichment and is found predominantly in lysosomes and occasionally early endosomes (Supplementary Figure 1H & I). Though there is a difference in expression level of these constructs (Supplementary Figure 1C), we posit that this phenotypic shift of S17N away from the lamellipodia plasma membrane could be due to direct changes in Rap2 activation. There is a possibility that these active/inactive constructs alter cellular dynamics in such a way that their own subcellular localization is indirectly affected. However, because we observe this same phenotype (decreased plasma membrane bound Rap2) throughout future figures, we would argue that this is a real direct effect on Rap2 subcellular localization.

**Figure 5.**
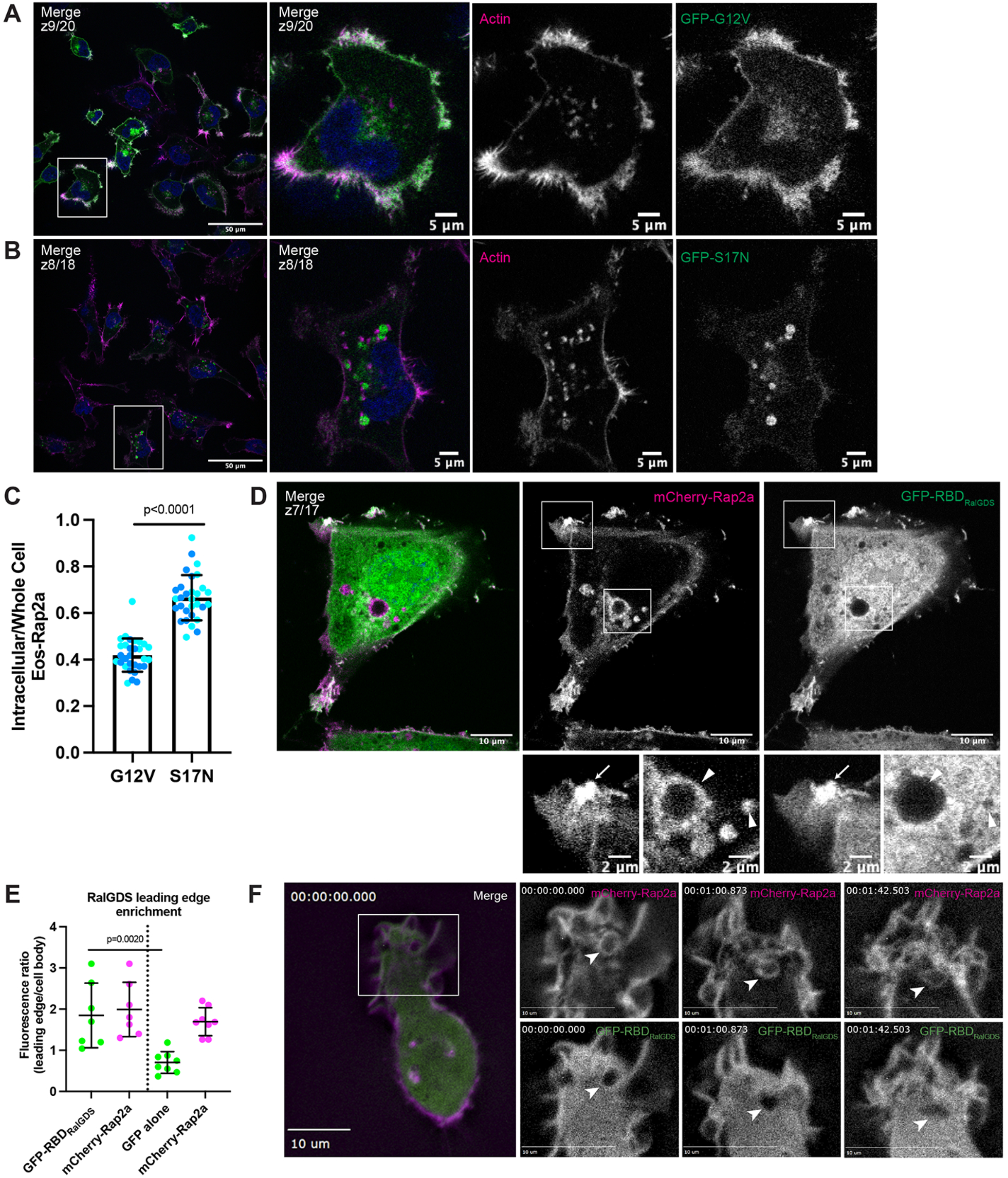
Rap2 activation is closely linked with its subcellular localization. (A) GFP-Rap2a-G12V localization. MDA-MB-231 cells stably expressing GFP-Rap2a-G12V were fixed and stained with Phalloidin (magenta) and DAPI (blue). Scale bars = 50µm, 5µm. (B) GFP-Rap2a-S17N localization. MDA-MB-231 cells stably expressing GFP-Rap2a-S17N were fixed and stained with Phalloidin (magenta) and DAPI (blue). Scale bars = 50µm, 5µm. (C) Intracellular to whole cell fraction quantification, Rap2a-G12V vs Gap2a-S17N. To quantify GFP-Rap2a localization changes (ie. decreased Eos-Rap2a at the plasma membrane in S17N background), intracellular/whole cell fractions were defined and calculated as the total fluorescence intensity of the intracellular GFP-Rap2a pool divided by the total fluorescence intensity of the whole cell GFP-Rap2a pool (see methods). Two biological replicates were performed for each cell line. In each biological replicate, 5 fields of view were imaged, with 3 cells analyzed from each field (total of 15 cells per n). The two shades of blue represent each set of 15 cells from n1 and n2. Each cell was treated as its own data point, and statistical analysis (unpaired t-test) was performed with n=30 total for each condition. Mean ± SD. G12V vs S17N p=<0.0001. (D) Co-localization of mCherry-Rap2a and GFP-RBD_RalGDS_. MDA-MB-231 cells were transiently transfected with mCherry-Rap2a and GFP-RBD_RalGDS_, then fixed. Arrow points to example of mCherry-Rap2a and GFP-RBD_RalGDS_ overlap at the plasma membrane. Arrowheads point to mCherry-Rap2a intracellular organelles, where GFP-RBDRalGDS is not present. Scale bars = 10µm, 2µm. (E) Quantification of GFP-RBD_RalGDS_ enrichment at the leading edge. In addition to the experiment in 3D, where cells were transfected with mCherry-Rap2a and GFP-RBDRalGDS, another experiment was performed side-by-side. MDA-MB-231 cells were transfected with mCherry-Rap2a and Free GFP (images not shown) to control for free FP localization at the leading edge. Fluorescence intensity ratios (leading edge/cell body ie. enrichment at ruffles) were calculated for individual cells (see methods) in both conditions. One biological replicate was performed, n=7 cells for GFP-RBD_RalGDS_ condition and n=8 cells for Free GFP condition. Mean ± SD. The two conditions are separated by the dotted line in the center of the graph. Green dots represent GFP-RBDRalGDS or Free GFP calculations, magenta represents mCherry-Rap2a. One-way ANOVA with Tukey’s multiple comparisons test. GFP-RBD_RalGDS_ vs GFP alone p=0.0020. (F) Live imaging of mCherry-Rap2a and GFP-RBD_RalGDS_ in MDA-MB-231 cells. Cells were transiently transfected with both mCherry-Rap2a and GFP-RBD_RalGDS_, plated on a collagen-coated glass dish, and imaged every 3 seconds using a 63x objective. Widefield microscope. Arrowheads point to an example of an mCherry-Rap2a positive organelle being internalized from the lamellipodia. GFP-RBD_RalGDS_ overlaps with mCherry-Rap2a primarily at the plasma membrane, and is not found present on the internalized organelle. See Video 7. Scale bars = 10µm.

To further investigate how Rap2 localization and activation are interconnected, we visualized Rap2 simultaneously with a known GTP-dependent effector, RalGDS (Spaargaren and Bischoff 1994; Franke, Akkerman, and Bos 1997; Nancy et al. 1999; Ohba, Kurokawa, and Matsuda 2003; Jeon et al. 2007; Liu et al. 2010). Based on our previous data, we hypothesized that a Rap2-effector complex would co-localize at the plasma membrane where the majority of Rap2 is active. MDA-MB-231s were transiently transfected with mCherry-Rap2a and GFP-RBD_RalGDS_ (Ras Binding Domain of RalGDS) and imaged using fluorescence microscopy. As shown in Figure 5D, GFP-RBD_RalGDS_ localizes to the leading edge, but is otherwise largely cytosolic. To firmly establish GFP-RBDRalGDS enrichment at ruffling lamellipodia, we used GFP alone as a control (Free GFP and mCherry-Rap2a co-transfection) and quantified the leading edge to cell body fluorescence ratio (Figure 5E). Our statistical analysis gives confidence to the co-occurrence and enrichment of Rap2a and RBDRalGDS at the leading edge. Importantly, mCherry-Rap2a and GFP-RBD_RalGDS_ overlap at the lamellipodia plasma membrane, but not at intracellular organelles (Figure 5D). These observations further support a model in which Rap2 is active at the plasma membrane and inactive when internalized, though still membrane bound.

Lastly, to characterize the spatiotemporal properties of Rap2 activation, we performed time-lapse analysis of mCherry-Rap2a concurrently with GFP-RBD_RalGDS_ (Figure 5F). MDA-MB-231 cells transiently expressing both constructs were imaged every second for several minutes, providing the fast dynamic information that is lost from fixed-cell imaging. As shown in Figure 5F, mCherry-Rap2a and GFP-RBD_RalGDS_ primarily co-localize at the leading-edge lamellipodia. Over time, when mCherry-Rap2a gets internalized from the plasma membrane, we observe that the mCherry-Rap2a organelle is not GFP-RBD_RalGDS_ positive. Though RalGDS is just one example of a Rap2 effector, we propose based on our collective data that Rap2 is active primarily at the plasma membrane.

Overall, these results suggest that endosomal trafficking is critical for Rap2 placement and that Rap2 activation is closely linked to its localization. Importantly, the high co-occurrence of Rap2 with actin supports a model in which Rap2 needs to be localized at the plasma membrane of lamellipodia in order to control actin dynamics. How Rap2 is continuously recycled between endocytic organelles and the plasma membrane, as well as how Rap2 is specifically targeted and activated at the leading-edge of migrating cells is an important next question.

### Targeting of Rap2 to the plasma membrane is dependent on the Rab40b/Cul5 complex

Ubiquitylation of small GTPases has emerged as a possible mechanism for regulating their localization, activation, and signaling (Nethe and Hordijk 2010; de la Vega, Burrows, and Johnston 2014; Dohlman and Campbell 2019). Notably, cumulative data suggests that mono-ubiquitylation regulates compartmentalization of the Ras protein family (HRas, KRas, and NRas), ultimately controlling location-specific signaling (Jura et al. 2006). However, how ubiquitylation regulates the Rap family of GTPases, and whether this is different from the Ras proteins, is essentially unknown. We hypothesized that ubiquitylation might serve to modulate spatiotemporal dynamics of Rap2 during cell migration. Importantly, Rap2 has been proposed as a putative substrate of the Rab40c/Cul5 complex in a *Xenopus* model (Lee et al. 2007). This study proposed that the XRab40c/Cul5 CRL governs poly-ubiquitylation of Rap2 in order to regulate the noncanonical Wnt signaling pathway, although how poly-ubiquitylation regulates Rap2 function and how this translates in a pathogenic context remains unclear. Furthermore, Rab40b KO in MDA-MB-231 cells leads to defects that partially phenocopy Rap2 KO, namely, decreased chemotactic migration and invasion (Linklater et al. 2021). Given these collective studies, we wondered whether the Rab40b/Cul5 complex interacts with and ubiquitylates Rap2 in mammalian cells and what role this ubiquitylation might play in regulating Rap2 localization and activation during mammalian cell migration (Figure 6A).

**Figure 6.**
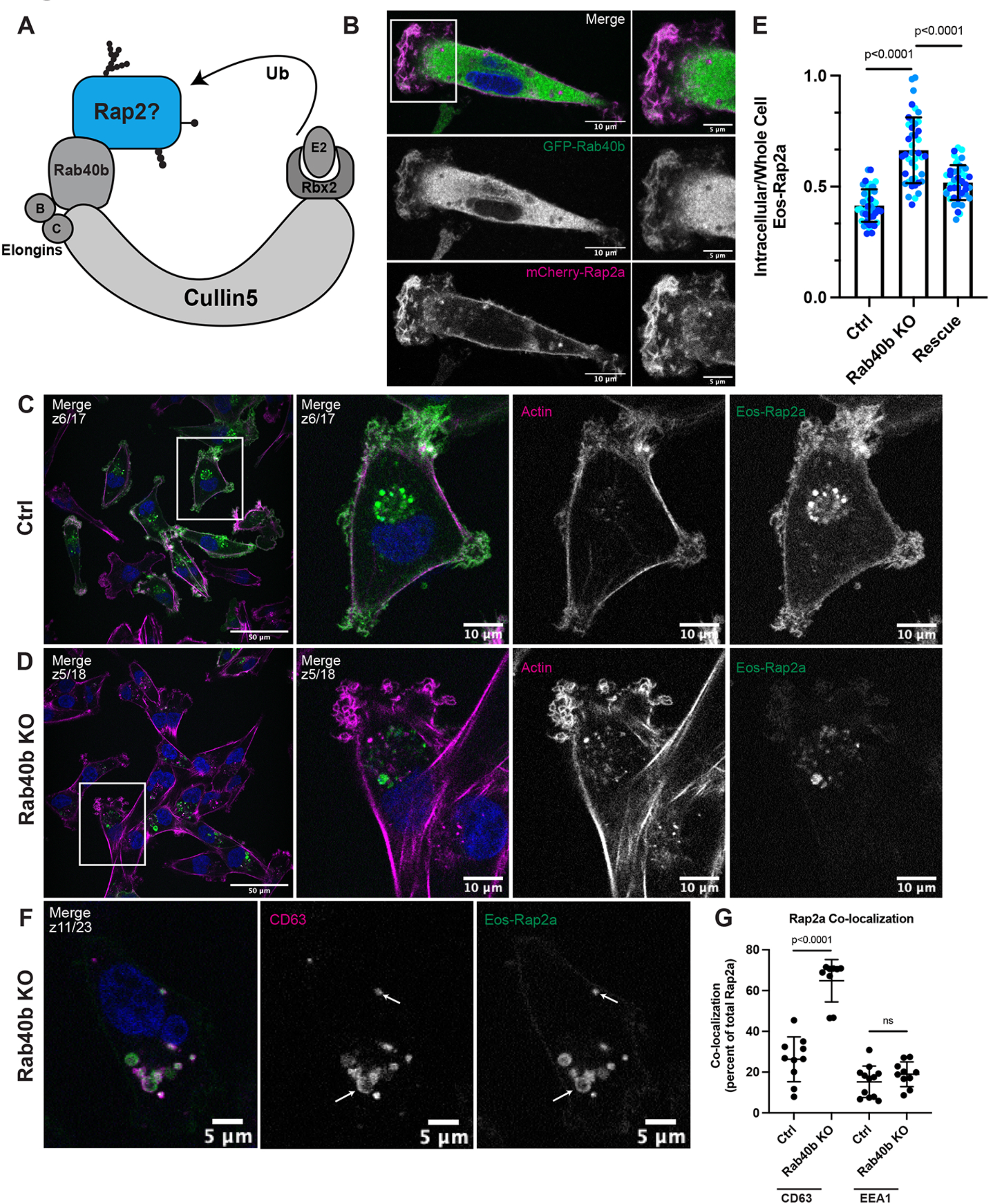
Loss of Rab40b results in a Rap2 sorting defect which leads to decreased Rap2 at the plasma membrane. (A) Rap2 is a putative substrate of the mammalian Rab40b/Cul5 complex. (B) Co-localization of GFP-Rab40b and mCherry-Rap2a. MDA-MB-231 cells stably expressing GFP-Rab40b were transiently transfected with mCherry-Rap2a, then fixed and stained with DAPI (blue). Scale bars = 10µm, 5µm. (C) Eos-Rap2a localization in Ctrl cells. Ctrl cells in this case are MDA-MB-231 parental cells. Eos-Rap2a has the same localization in MDA-MB-231 parental cells vs Cas9 cells (data not shown). MDA-MB-231 cells stably expressing Eos-Rap2a were fixed and stained with Phalloidin (magenta) and DAPI (blue). Scale bars = 50µm, 10µm. (D) Eos-Rap2a localization in Rab40b KO cells. Rab40b KO MDA-MB-231 cells stably expressing Eos-Rap2a were fixed and stained with Phalloidin (magenta) and DAPI (blue). Scale bars = 50µm, 10µm. (E) Intracellular to whole cell fraction quantification, Eos-Rap2a in Ctrl vs Rab40b KO vs Rab40b KO Rescue cells. Rab40b KO Rescue cells overexpress FLAG-Rab40b WT (stable population line, see Supplemental Figure 1E). To quantify Eos-Rap2a localization changes (ie. decreased Eos-Rap2a at the plasma membrane), intracellular/whole cell fractions were defined and calculated as the total fluorescence intensity of the intracellular Eos-Rap2a pool divided by the total fluorescence intensity of the whole cell Eos-Rap2a pool (see methods). Three biological replicates were performed for each cell line. In each biological replicate, 5 fields of view were imaged, with 3 cells analyzed from each field (total of 15 cells per n). The three shades of blue represent each set of 15 cells from n1, n2, and n3. Each cell was treated as its own data point, and statistical analysis (one-way ANOVA with Tukey’s multiple comparisons test) was performed with n=45 total for each condition. Mean ± SD. Eos-Rap2a in Ctrl cells vs Eos-Rap2a in Rab40b KO cells p=<0.0001. Eos-Rap2a in Rab40b KO cells vs Eos-Rap2a in Rab40b KO Rescue cells p=<0.0001. ROUT outlier test removed three outliers from Rab40b KO data set. Ctrl cells in this experiment are MDA-MB-231 parental cells. (F) Eos-Rap2a co-localization with CD63 in Rab40b KO cells. Rab40b KO MDA-MB-231 cells stably expressing Eos-Rap2a were fixed and stained with the lysosomal marker CD63 (magenta) and DAPI (blue). Arrows indicate examples of Eos-Rap2a and CD63 overlap. Scale bars = 5µm. (G) Quantification of co-localization between Rap2a and CD63/EEA1 in Ctrl vs Rab40b KO cells. Ctrl or Rab40b KO cells stably expressing Eos-Rap2a were fixed and stained with either CD63 (late endosomes/lysosomes) or EEA1 (early endosomes). 3i SlideBook6 software was used to calculate the percent of total GFP-Rap2a that co-localizes with the markers indicated (see methods). Two biological replicates were performed, with roughly 5 cells imaged for each replicate. n=10 Ctrl CD63, n=9 KO CD63, n=12 Ctrl EEA1, n=10 KO EEA1. One-way ANOVA with Tukey’s multiple comparisons test. Ctrl vs Rab40b KO CD63 p=<0.0001. Ctrl vs Rab40b KO EEA1 ns=non-significant.

To start dissecting whether the Rab40b/Cul5 complex regulates Rap2 during breast cancer cell migration, we first asked whether Rab40b and Rap2 co-localize in MDA-MB-231 cells. We have previously shown that GFP-Rab40b localizes to the leading-edge of migrating cells where it guides actin ruffling at lamellipodia (Jacob et al. 2013; Linklater et al. 2021). Since Rap2 appears to function at the leading-edge during cell migration, we hypothesized that Rab40b and Rap2 may co-localize at the leading-edge plasma membrane. To this end, MDA-MB-231 cells stably expressing GFP-Rab40b were transiently transfected with mCherry-Rap2a and analyzed by immunofluorescence. As predicted, we find that GFP-Rab40b and mCherry-Rap2a co-localize at the leading-edge lamellipodia of the cell (Figure 6B). The overlap of these two proteins at the plasma membrane support a possible link between Rab40b, Rap2, and cell migration.

Since Rap2 recycling between the endolysosomal compartment and lamellipodia plasma membrane appears to regulate its activity and function, we next wondered whether localization of Rap2 is affected in our Rab40b KO cell line. For this experiment, we used the fluorescent protein Eos, which we found less sensitive to low pH compared to GFP, thus allowing us to better visualize lysosomal Rap2 (Wiedenmann et al. 2004; Zhang et al. 2012). As shown in Figure 6C, Eos-Rap2a in MDA-MB-231 control cells localizes identically to GFP-Rap2a, with two major populations: plasma membrane bound and intracellular endolysosomal organelles (Figure 6C). Remarkably, Eos-Rap2a levels at the plasma membrane are greatly diminished in Rab40b KO cells, with increased Rap2a accumulation in intracellular organelles (Figure 6D). Quantification of Eos-Rap2a fluorescence intensity reveals a higher intracellular to whole cell fraction in Rab40b KO cells compared to control cells (Figure 6E, Supplementary 2C). This is not a result of decreased expression, as exogenous levels of Eos-Rap2a are equal in both control and Rab40b KO cell lines (Supplementary Figure 2A). Importantly, we can partially rescue Eos-Rap2a mis-localization with the addition of Rab40b-WT (Figure 6E, Supplementary Figure 1E).

We again co-stained these cells with known endocytic pathway markers, namely EEA1 (early endosome) and CD63 (early endosome/late endosome) and measured co-localization with Rap2a. The enriched intracellular Eos-Rap2a in Rab40b KO cells co-localizes with both EEA1 and CD63 positive organelles (Figure 6F, Supplementary Figure 1J). Interestingly though, we find that Eos-Rap2a has a higher co-occurrence with CD63 in Rab40b KO cells compared to control cells, whereas EEA1 overlap is unchanged (Figure 6G). Given our earlier experiments suggesting that Rap2 is dynamically trafficked and likely recycled to the plasma membrane, this increase in CD63 accumulation suggests that Rab40b/Cul5-dependent ubiquitylation may be required for Rap2 sorting at early endosomes, away from lysosomes, and back to the plasma membrane. Importantly, Rab40b depletion also leads to similar defects in Eos-Rap2b and Eos-Rap2c subcellular distribution (Supplementary Figure 2A, D-G), suggesting that the three Rap2 isoforms are regulated and function in a similar fashion. We propose, based on these results, that Rab40b regulates Rap2 sorting at early endosomes for recycling to the lamellipodia plasma membrane.

### Rap2 is a Rab40b binding protein

To begin understanding how Rab40b regulates Rap2 localization, we next assessed binding between these two proteins. Interestingly, both Rap2 and Rab40b are small monomeric GTPases, thus raising the question of whether their interaction is dependent on the nucleotide status of Rap2 or Rab40b (or both). To this end, we incubated purified recombinant GST-Rap2a with MDA-MB-231 cell lysates stably expressing FLAG-Rab40b and analyzed their interaction using GST pull-down assays. Importantly, we find that the binding between these two proteins is independent of Rap2a nucleotide status, (Figure 7A & C), but is enhanced when Rab40b is loaded with the non-hydrolyzable form of GTP, GMP-PNP (Figure 7B & D, left panels). This suggests that Rap2a predominately binds Rab40b-GTP. We note that the binding is fairly weak, suggesting that the interaction between these proteins is likely quite transient, as would be expected between an enzyme (E3 ligase complex) and its substrate (Rap2). Additionally, we tested binding between Rab40b and the other Rap2 isoforms and find that Rab40b also interacts with Rap2b and Rap2c, suggesting again that the three Rap2 isoforms may be regulated in a similar manner (Supplementary Figure 3A).

**Figure 7.**
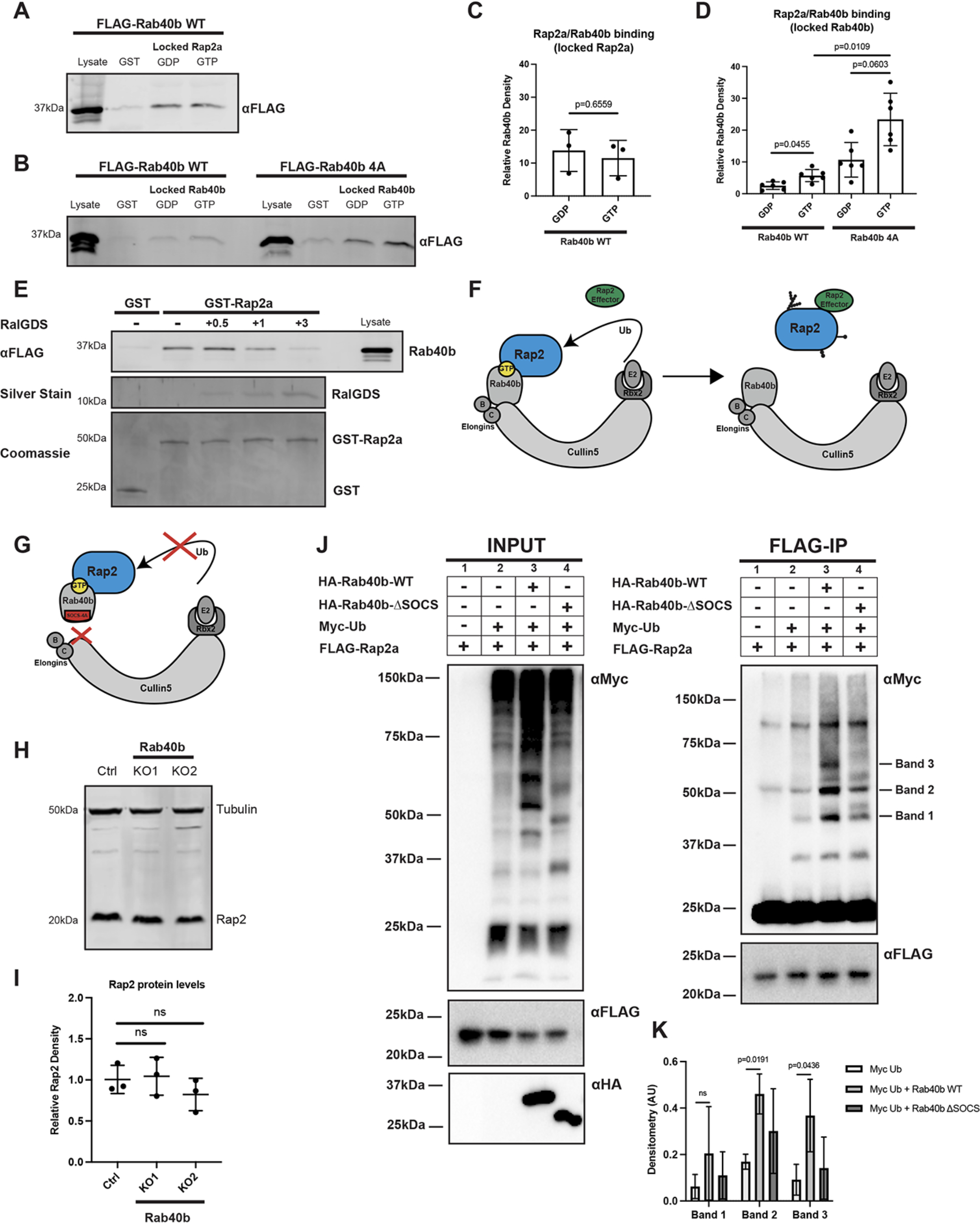
Rap2 is a Rab40b binding protein and is ubiquitylated by the Rab40b/Cul5 complex. (A) Rab40b binding to locked Rap2a. MDA-MB-231 lysates stably expressing FLAG-Rab40b WT were incubated with either GST or GST-Rap2a, followed by a GST pull-down assay. Before incubation, GST-Rap2a was loaded with either GDP or GMP-PNP (labeled as GTP for brevity). GST alone was used to control for GST binding to Rab40b. 25µg of lysate input was loaded as a positive control and used to estimate pull-down efficiency. Western blot was probed with αFLAG antibody. Coomassie gel showing equal levels of GST and GST-Rap2a is in Supplementary Figure 5. (B) Rap2a binding to locked Rab40b. MDA-MB-231 lysates stably expressing FLAG-Rab40b WT or SOCS-4A (left and right, respectively) were incubated with either GST or GST-Rap2a, followed by a GST pull-down. Before incubation, FLAG-Rab40b lysates were loaded with either GDP or GMP-PNP (labeled as GTP for brevity). GST alone was used to control for GST binding to Rab40b. 25µg of lysate input was loaded as a positive control and used to estimate pull-down efficiency. Western blot was probed with αFLAG antibody. Coomassie gel showing equal levels of GST and GST-Rap2a is in Supplementary Figure 5. (C) Quantification of GST pull-down in (A). Three biological replicates were performed. Mean ± SD. GST signal was subtracted from GDP/GTP, and “Relative Rab40b Density” was calculated by normalizing to lysate. Unpaired t-test. Rab40b WT GDP vs GTP (locked Rap2a) p=0.6559. GTP=GMP-PNP. (D) Quantification of GST pull-down in (B). Six biological replicates were performed. Mean ± SD. GST signal was subtracted from GDP/GTP, and “Relative Rab40b Density” was calculated by normalizing to lysate. Brown-Forsythe ANOVA with Dunnett’s T3 multiple comparisons test. Rab40b WT GDP vs GTP (locked Rab40b) p=0.0455. Rab40b SOCS-4A GDP vs GTP (locked Rab40b) p=0.0603. Rab40b WT GTP vs Rab40b SOCS-4A GTP (locked Rab40b) p=0.0109. GTP=GMP-PNP. (E) Competitive Binding Experiment between FLAG-Rab40b, GST-Rap2a, and untagged RalGDS (RBD). MDA-MB-231 lysates stably expressing FLAG-Rab40b WT (GMP-PNP loaded) were incubated with either GST or GST-Rap2a (GMP-PNP loaded) and increasing concentrations of RalGDS (0.5, 1, and 3 times the amount of GST/GST-Rap2a). GST alone was used to control for GST binding to Rab40b. 15µg of lysate input was loaded as a positive control and used to estimate pull-down efficiency. Western blot was probed with αFLAG antibody. Coomassie gel shows equal levels of GST and GST-Rap2a. Silver Stain was used to visualize RalGDS binding to GST-Rap2a. With increasing concentrations of RalGDS, we observe a concurrent decrease in FLAG-Rab40b binding to GST-Rap2a. (F) Model summarizing findings in A-D. Rap2 preferentially binds Rab40b-GTP (left). Ubiquitylation of Rap2 by Rab40b/Cul5 and subsequent complex dissociation is necessary for Rap2 to interact with its downstream effector (in a GTP dependent manner). (G) When the Rab40b/Cul5 complex is disrupted (SOCS-4A mutant), Rab40b binds more strongly to Rap2. We propose this is due to lack of ubiquitylation/complex dissociation. (H) Rap2 protein levels Ctrl vs Rab40b KO cells. Ctrl and Rab40b KO MDA-MB-231 lysates were probed for αRap2 and αTubulin (loading control). Ctrl cells are dox-inducible Cas9 MDA-MB-231s that were used to generate CRISPR lines. 50µg of lysate was loaded for each sample. (I) Quantification of western blot in (H). Three biological replicates were performed. Relative intensity of Rap2 was normalized to the levels of Tubulin and to Ctrl cells. Mean ± SD. One-way ANOVA with Tukey’s multiple comparisons test. ns=non-significant. (J) Rap2a ubiquitylation in HEK293T cells. 293Ts were transfected with pRK5-FLAG-Rap2a +/-pRK5-Myc-Ub, pRK7-HA-Rab40b or pRK7-HA-Rab40b ýSOCS. After 24hrs of transfection, cells were harvested, lysed, and immunoprecipitated with αFLAG. Left column shows lysates (input) probed for Myc, FLAG, and HA. Right column shows FLAG immunoprecipitates probed for Myc and FLAG. Tick marks on the right blot indicate the Rap2-ubiquitin bands that are increased in response to Rab40b addition. (K) Quantification of 293T ubiquitylation assay in (J). Raw densitometry was measured for Band 1, Band 2, and Band 3 across four independent experiments. These values (arbitrary unites) were then normalized to FLAG-Rap2a input levels and graphed for conditions 2, (Myc-Ub), 3 (Myc-Ub + Rab40b WT) and 4 (Myc-Ub + Rab40b ýSOCS). Four biological replicates were performed. Mean ± SD. Two-way ANOVA with Tukey’s multiple comparisons test. Band 1 Myc-Ub vs Myc-Ub + Rab40b WT ns=non-significant. Band 2 Myc-Ub vs Myc-Ub + Rab40b WT p=0.0191. Band 3 Myc-Ub vs Myc-Ub + Rab40b WT p=0.0436.

Since this interaction is independent of Rap2 nucleotide state, we wondered whether Rap2 can bind its downstream effector proteins in a GTP dependent manner while part of a Rab40b/Rap2 complex, or, whether these interactions are mutually exclusive. To test this, we performed a competitive binding experiment: FLAG-Rab40b lysates were incubated with GST-Rap2a as before, but with increasing concentrations of the Rap2 effector, RalGDS (Figure 7E). Importantly, we discover that there is indeed competition for Rap2 binding between Rab40b and RalGDS. With increasing concentrations of effector, we see a decrease in FLAG-Rab40b binding to GST-Rap2a and a concurrent increase in RalGDS pulled down with GST-Rap2a (Figure 7E), suggesting that Rap2 cannot bind both proteins simultaneously. Overall, this data suggests that Rap2 dissociation from the Rab40b/Cul5 complex, likely after ubiquitin modification, is required for its interaction with downstream effectors (Figure 7F).

To understand the role of Cul5 in regulating Rap2 function we next asked whether Rab40b binding to Rap2 is dependent on Cul5. To this end, we incubated purified GST-Rap2a with MDA-MB-231 cells stably expressing a FLAG-Rab40b SOCS-4A mutant, which disrupts binding to Cul5 and formation of the CRL complex (Figure 7G) (Linklater et al. 2021; Duncan et al. 2021). We find that the SOCS-4A mutation does not block Rap2a and Rab40b binding (Figure 7B, right panel), suggesting that Rab40b is able to bind its substrates independent of Cul5 presence. Though this was our original prediction, and was intended as an experimental control, we observe an intriguing result: the Rab40b SOCS-4A mutant binds to Rap2a stronger than Rab40b WT (Figure 7D, right panel). This observation would be consistent with the idea that Rap2 is a Rab40b/Cul5 complex substrate, since blocking ubiquitylation would likely prevent dissociation between the E3 ligase complex (Rab40b/Cul5) and its substrate protein (Rap2) (Figure 7G). Furthermore, this fits with the idea that Rap2 must be released from the Rab40b/Cul5 complex in order to bind its putative effectors (Figure 7F). Overall, these data support a model in which Rab40b-GTP binds Rap2 that Rap2 is a putative substrate for ubiquitylation by the Rab40b/Cul5 CRL.

### Rap2 is ubiquitylated by the Rab40b/Cul5 complex

Given that mammalian Rab40b and Rap2 interact and our data suggesting Rap2 is not targeted properly in the absence of Rab40b, we next asked whether the Rab40b/Cul5 complex regulates proteasomal degradation of Rap2 via poly-ubiquitylation. To do this, we measured total protein levels of Rap2 in two different Rab40b KO cell lines as well as two other cell lines expressing either a Rab40b SOCS-4A or Rab40b ΔSOCS mutant (Linklater et al. 2021; Duncan et al. 2021). Our previous studies using the Rab40b SOCS-4A mutant demonstrated only a partial loss of Cul5 binding, thus, we generated a Rab40b ΔSOCS mutant (residues 1-174 followed by residues 229-278, so that Rab40b is missing the SOCS box proper, but still contains the C-terminal prenylation site) to increase disruption of Rab40b/Cul5 complex formation (Supplementary Figure 3B). We hypothesized that if the Rab40b/Cul5 complex regulates proteasomal degradation of Rap2, that Rap2 levels should be increased in the absence of Rab40b as well as when the Rab40b/Cul5 complex is disrupted. This is based on the well accepted doctrine in the field that canonical poly-ubiquitin chains (typically K48-linked) signal for proteasomal degradation via the 26S proteasome (Pickart and Fushman 2004; Komander and Rape 2012). Surprisingly, we observe no significant change in Rap2 protein levels across our Rab40b KO cell lines nor the Rab40b SOCS mutants (Figure 7H & I, Supplementary Figure 3C & D). This is interesting, as it suggests that the Rab40b/Cul5 complex does not mediate canonical ubiquitin-dependent degradation of Rap2. This supports previous observations about Rap2 stability in *Xenopus*, where the authors did not observe any effect on Rap2 stability when cells were treated with XRab40c or XCul5 morpholinos (Lee et al. 2007). One caveat to the Rab40b KO cell line is the presence of the other Rab40 isoforms, which may compensate for the lack of Rab40b and prevent us from detecting protein level changes. To alleviate this, and since there is also evidence of Rab40c-dependent regulation of Rap2, we measured Rap2 protein levels in two different MDA-MB-231 Rab40 KO cell lines, where cells lack the three Rab40 family members Rab40a, Rab40b, and Rab40c (Supplementary Figure 3E) (Linklater et al. 2021). Importantly, we once again do not detect any significant change in global Rap2 protein levels in the Rab40 KO cell lines, suggesting that the Rab40 GTPases do not mediate ubiquitin-dependent proteasomal degradation of Rap2.

At this point, our data postulates that the Rab40b/Cul5 complex binds and regulates Rap2 localization, without affecting global protein levels of Rap2. Additionally, our localization data suggests a potential early endosome sorting defect in Rab40b KO cells, leading to increased targeting of Rap2 to lysosomes. Based on these observations and what we know about the ubiquitin code, we speculated that Rap2 might be subjected to a non-canonical ubiquitin tag (ie. not a ubiquitin chain signaling for proteasomal degradation) that may regulate its sorting and targeting to the lamellipodia plasma membrane. (Swatek and Komander 2016; Akutsu, Dikic, and Bremm 2016). To begin testing this idea, we utilized HEK293T cells to test direct ubiquitylation of Rap2 by the Rab40b/Cul5 complex.

Using 293T cells, we set up four different transfection conditions: 1. FLAG-Rap2a alone 2. FLAG-Rap2a + Myc-Ub 3. FLAG-Rap2a + Myc-Ub + HA-Rab40b WT and 4.

FLAG-Rap2a + Myc-Ub + HA-Rab40b ΔSOCS (Figure 7J, left panel). In each case, we immunoprecipitated with anti-FLAG and western blotted for anti-Myc (Figure 7J, right panel). Importantly, we observe stimulation of Rap2 ubiquitylation by Rab40b WT (condition 3) compared to just Myc-Ub addition (condition 2). We focused on Rap2-Ub species that were stimulated by Rab40b in all four repeats. First, the Rap2-Ub species labeled as “Band 1” appears to be a specific band (not present in condition 1), where we detect an upward trend when Rab40b-WT is added (condition 3), but less enhanced by Rab40b ΔSOCS (condition 4) (Figure 7J & K). Based solely on the molecular weight ladder, we hypothesize that “Band 1” corresponds to a di-Ub Rap2 species, but further work will be needed to directly prove this. Next, we also see “Band 2” and “Band 3” are enhanced by the addition of Rab40b-WT, but decreased in the case of Rab40b ΔSOCS (Figure 7J & K). When analyzing all four experiments, “Band 2” and “Band 3” seem to be the strongest increase with a concurrent trend downward in the Rab40b ΔSOCS condition (Figure 7K). Again, based on size, we hypothesize that these bands correspond to the addition of three-five ubiquitin moieties on Rap2. One caveat is that “Band 2” appears in the FLAG-Rap2a only control (far left column, condition 1). Despite our troubleshooting efforts to increase wash stringency and use a secondary antibody specific to the light chain of IgG, we believe that these are non-specific bands that our secondary antibody detects, likely remnants of IgG heavy chain. Nonetheless, data from four independent experiments strongly suggest that “Band 2” and “Band 3” are stimulated by Rab40b-WT compared to control. Finally, we see little indication of Rab40b-mediated poly-ubiquitylation of Rap2, which would appear as a high molecular weight shmear instead of distinct bands. This is consistent with our data showing that loss of Rab40b does not mediate proteasomal targeting of Rap2 (Figure 7H & I). Given this, we hypothesize that these non-poly chains (ranging from two to five ubiquitin moieties) are likely linkages with a unique downstream signal, possibly to regulate sorting and targeting to the plasma membrane. Though further work will be needed to define the composition of these ubiquitin species (multi-mono, etc), collectively, these results support Rap2 as a substrate of the Rab40b/Cul5 complex.

### Ubiquitylation of specific Rap2 lysines regulates its subcellular localization

Summarizing our data thus far, we propose that the Rab40b/Cul5 complex regulates Rap2 function via direct ubiquitylation. To start dissecting the molecular consequence of Rap2 ubiquitylation, we sought to identify the specific lysine(s) on Rap2 that are modified by ubiquitin. Previous work on Rap2a in neurons identified four lysines of Rap2a that are putative targets for Nedd4-1-mediated ubiquitylation (Kawabe et al. 2010). The authors originally identified nine lysines of Rap2a as solvent exposed, and found that mutation of four of these lysines (K5, K94, K148, and K150) was able to efficiently reduce Nedd4-1-mediated Rap2a ubiquitylation. Additionally, the initial study implicating Rap2 as a possible Rab40c/Cul5 substrate proposed a single lysine mutation at position 117 was sufficient to reduce ubiquitylation of Rap2 (Lee et al. 2007). Combining these observations, we decided to generate a Rap2a-K5R construct (K5, K94, K117, K148, K150), with the goal of understanding how localization, sorting, and activation are affected by mutation of putative ubiquitylation sites (Supplementary Figure 3F).

One potential concern is that mutation of K5, K94, K117, K148, and K150 alters Rap2 folding and blocks GTP loading. To refute this possibility, we used an endpoint phosphate assay to measure GTP hydrolysis as a readout for proper Rap2 folding.

Purified GST-Rap2a-WT and GST-Rap2a-K5R were incubated side-by-side with CytoPhos reagent, resulting in a colorimetric change upon inorganic phosphate release. We find that Rap2a-K5R is capable of binding and hydrolyzing GTP, suggesting that the K5R mutation does not affect overall Rap2a folding (Supplementary Figure 3G). In fact, we observe that Rap2a-K5R may in fact hydrolyze GTP faster than its WT counterpart, though it is unclear if this ∼2-fold change is biologically relevant.

Moving forward, we first asked whether Rap2a-K5R localization is altered compared to Rap2a-WT. To this end, MDA-MB-231 cells stably expressing Eos-Rap2a-K5R (Supplementary Figure 3H) were analyzed by immunofluorescence. Somewhat striking, we find that Eos-Rap2a-K5R is predominately localized to the endolysosomal compartment (Figure 8A). We immediately noticed the similarity between Eos-Rap2a-K5R localization and what we observe when Eos-Rap2a-WT is expressed in a Rab40b KO background. Indeed, fluorescence intensity analysis of Rap2a-K5R reveals a higher intracellular to whole cell fraction compared to Rap2a-WT (Figure 8B, Supplementary 2C).

**Figure 8.**
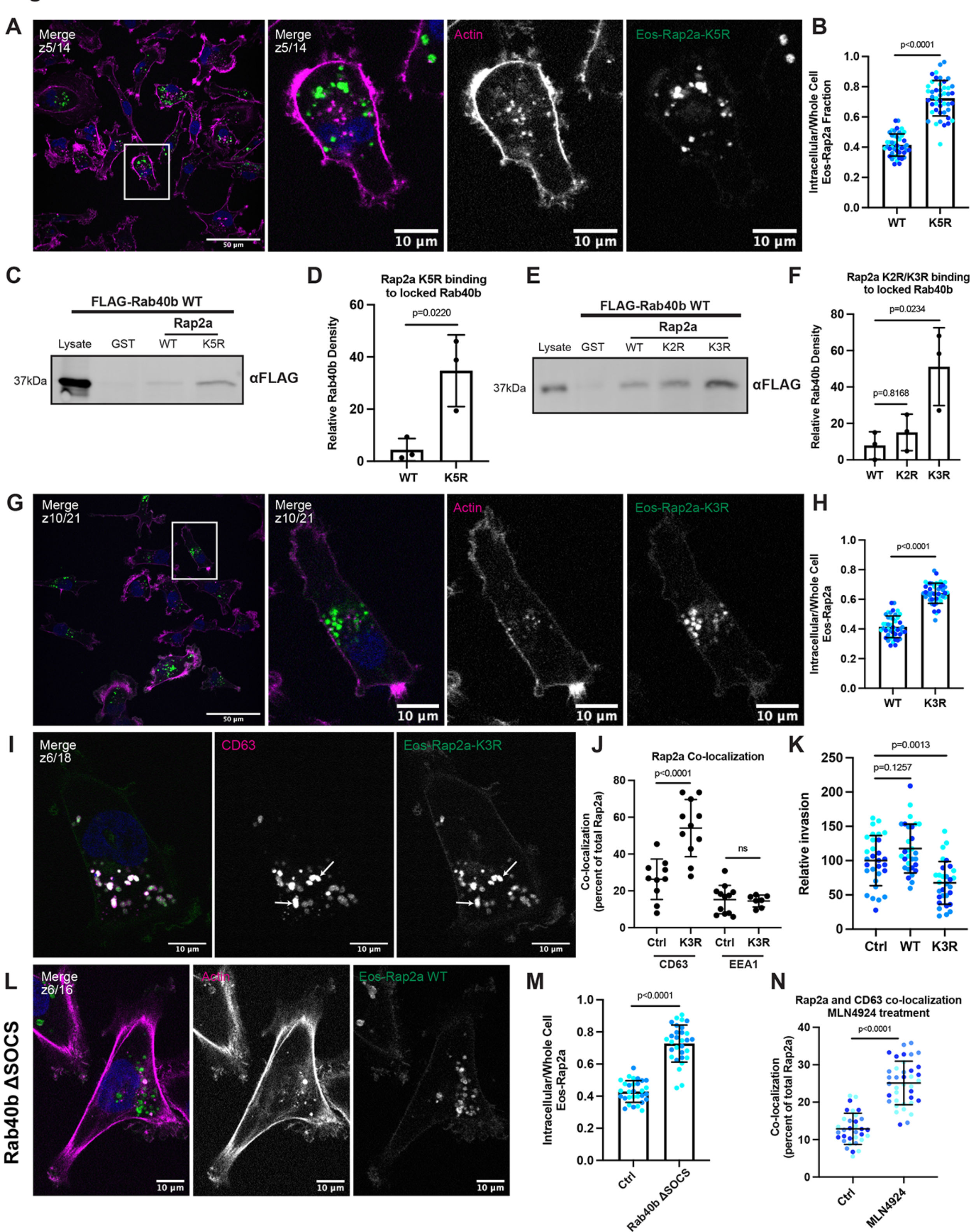
Mutation of five putative ubiquitylation sites within Rap2 affects recycling, subcellular localization, binding to Rab40b, and cell invasion. (A) Eos-Rap2a-K5R localization. MDA-MB-231 cells stably expressing Eos-Rap2a-K5R were fixed and stained with Phalloidin (magenta) and DAPI (blue). Scale bars = 50µm, 10µm. (B) Intracellular to whole cell fraction quantification, Eos-Rap2a-WT vs Eos-Rap2a-K5R. To quantify Eos-Rap2a localization changes, intracellular/whole cell fractions were defined and calculated as the total fluorescence intensity of the intracellular Eos-Rap2a pool divided by the total fluorescence intensity of the whole cell Eos-Rap2a pool (see methods). Three biological replicates were performed for each cell line. In each biological replicate, 5 fields of view were imaged, with 3 cells analyzed from each field (total of 15 cells per n, color coded). Each cell was treated as its own data point, and statistical analysis (unpaired t-test) was performed with n=45 total for each condition. Mean ± SD. Eos-Rap2a-WT vs Eos-Rap2a-K5R p=<0.0001. Eos-Rap2a-WT data points are from Figure 6E. (C) Rap2a-K5R binding to Rab40b. MDA-MB-231 lysates stably expressing FLAG-Rab40b WT were incubated with either GST, GST-Rap2a-WT or -K5R, followed by a GST pull-down. Before incubation, FLAG-Rab40b lysates were loaded with GMP-PNP. GST alone was used to control for GST binding to Rab40b. 25µg of lysate input was loaded as a positive control and used to estimate pull-down efficiency. Western blot was probed with αFLAG antibody. Coomassie gel showing equal levels of GST, GST-Rap2a-WT and -K5R is in Supplementary Figure 5. (D) Quantification of GST pull-down in (C). Three biological replicates were performed. Mean ± SD. GST signal was subtracted from experimental lanes, and “Relative Rab40b Density” was calculated by normalizing to lysate. Unpaired t-test. Rap2a-WT vs Rap2a-K5R p=0.0220. (E) Rap2a-K2R and Rap2a-K3R binding to Rab40b. MDA-MB-231 lysates stably expressing FLAG-Rab40b WT were incubated with either GST, GST-Rap2a-WT, -K2R, or -K3R followed by a GST pull-down. Before incubation, FLAG-Rab40b lysates were loaded with GMP-PNP. GST alone was used to control for GST binding to Rab40b. 25µg of lysate input was loaded as a positive control and used to estimate pull-down efficiency. Western blot was probed with αFLAG antibody. Coomassie gel showing equal levels of GST, GST-Rap2a-WT, -K2R, and -K3R is in Supplementary Figure 5. (F) Quantification of GST pull-down in (E). Three biological replicates were performed. Mean ± SD. GST signal was subtracted from experimental lanes, and “Relative Rab40b Density” was calculated by normalizing to lysate. One-way ANOVA with Tukey’s multiple comparisons test. Rap2a-WT vs Rap2a-K2R p=0.8168. Rap2a-WT vs Rap2a-K3R p=0.0234. (G) Eos-Rap2a-K3R localization. MDA-MB-231 cells stably expressing Eos-Rap2a-K3R were fixed and stained with Phalloidin (magenta) and DAPI (blue). Scale bars = 50µm, 10µm. (H) Intracellular to whole cell fraction quantification, Eos-Rap2a-WT vs Eos-Rap2a-K3R. To quantify Eos-Rap2a localization changes, intracellular/whole cell fractions were defined and calculated as the total fluorescence intensity of the intracellular Eos-Rap2a pool divided by the total fluorescence intensity of the whole cell Eos-Rap2a pool (see methods). Three biological replicates were performed for each cell line. In each biological replicate, 5 fields of view were imaged, with 3 cells analyzed from each field (total of 15 cells per n, color coded). Each cell was treated as its own data point, and statistical analysis (unpaired t-test) was performed with n=45 total for each condition. Mean ± SD. Eos-Rap2a-WT vs Eos-Rap2a-K3R p=<0.0001. Eos-Rap2a-WT data points are from Figure 6E. (I) Eos-Rap2a-K3R co-localization with CD63. MDA-MB-231 cells stably expressing Eos-Rap2a-K3R were fixed and stained with the lysosomal marker CD63 (magenta) and DAPI (blue). Arrows indicate examples of Eos-Rap2a-K5R and CD63 overlap. Scale bars = 10µm. (J) Quantification of co-localization between Rap2a and CD63/EEA1 in Ctrl vs K3R cells. MDA-MB-231s stably expressing either Eos-Rap2a WT or Eos-Rap2a K3R were fixed and stained with either CD63 (late endosomes/lysosomes) or EEA1 (early endosomes). 3i SlideBook6 software was used to calculate the percent of total GFP-Rap2a that co-localizes with the markers indicated (see methods). Two biological replicates were performed, with roughly 5 cells imaged for each replicate. n=10 Ctrl CD63, n=11 K3R CD63, n=12 Ctrl EEA1, n=7 K3R EEA1. One-way ANOVA with Tukey’s multiple comparisons test. Mean ± SD. Ctrl vs K3R CD63 p=<0.0001. Ctrl vs K3R EEA1 ns=non-significant. (K) Invasion assay quantification, Ctrl vs Eos-Rap2a WT vs Eos-Rap2a K3R. Three biological replicates were performed, with technical duplicates in each experiment. 5 fields of view were imaged for each Matrigel insert (resulting in 10 fields of view per experiment per condition, color coded). Raw number of cells invaded per field of view were normalized to Ctrl (“Relative invasion”). Ctrl cells are MDA-MB-231 parental cells. Eos-Rap2a WT and K3R are stable overexpressed lines. Each field of view was treated as its own data point (n=30). One-way ANOVA with Tukey’s multiple comparisons test. Mean ± SD. Ctrl vs Eos-Rap2a WT p=0.1257. Ctrl vs Eos-Rap2a K3R p=0.0013. (L) Eos-Rap2a localization in Rab40b ΔSOCS cells. Rab40b ΔSOCS MDA-MB-231 cells expressing Eos-Rap2a were fixed and stained with Phalloidin (magenta) and DAPI (blue). Scale bars = 50µm, 10µm. (M) Intracellular to whole cell fraction quantification, Eos-Rap2a in Ctrl cells vs Rab40b ΔSOCS cells. Two biological replicates were performed for each cell line. In each biological replicate, 5 fields of view were imaged, with 3 cells analyzed from each field (total of 15 cells per n, color coded). Each cell was treated as its own data point, and statistical analysis (unpaired t-test) was performed with n=30 total for each condition. Mean ± SD. Eos-Rap2a Ctrl vs Eos-Rap2a ΔSOCS cells p=<0.0001. Eos-Rap2a-WT data points are from Figure 6E. (N) Rap2a and CD63 co-localization analysis in Ctrl cells vs cells treated with neddylation/Cul5 inhibitor MLN4924. MDA-MB-231 cells stably expressing GFP-Rap2a were treated with either DMSO (Ctrl) or 300nM MLN4924 for 24 hrs. Cells were then fixed and stained for CD63, following co-localization analysis. 3i SlideBook6 software was used to calculate the percent of total GFP-Rap2a that co-localizes with CD63 (see methods). Three biological replicates were performed, with roughly 12 cells imaged for each replicate (color coded for each n). Mean ± SD. n=33 for Ctrl/DMSO treated cells, n=38 for MLN4924 treated cells. Unpaired t-test. p=<0.0001.

So far, the data reveal that mutation of Rap2 at five lysines dramatically affects its subcellular localization. To begin tying this back to Rab40b and cell migration, we next wondered how mutation of these five lysines might affect Rap2 binding to the Rab40b/Cul5 complex. To test this, we incubated purified GST-Rap2a-K5R with MDA-MB-231 FLAG-Rab40b lysates and performed a GST pull-down assay. Notably, we discover that Rap2a-K5R binds to Rab40b much stronger than Rap2a-WT (Figure 8C & D). Again, this result reminded us of our previous observation: increased binding between Rap2a-WT and Rab40b SOCS-4A. If we hypothesize that increased binding is due to lack of dissociation, one prediction is that the Rab40b/Cul5 complex binds Rap2a-K5R but cannot ubiquitylate and dissociate from one another. This would suggest that Rab40b/Cul5 may target at least one or more of the five lysines within the Rap2a-K5R mutant.

After examining the crystal structure of Rap2a (PDB 2RAP), we noticed that the five lysines in question are clustered into two general regions: three lysines near the GTP pocket (K117, K148, and K150), and two lysines removed from the GTP pocket (K5 and K94). To begin dissecting the function of the five lysines within our Rap2a-K5R mutant, we split the mutations into two groups: K2R (K5, K94) and K3R (K117, K148, K150) (Supplementary Figure 3F). Given the increased binding of Rap2a-K5R to Rab40b, we began with this binding assay as a read-out of which mutant (K2R or K3R) might house the relevant Rap2 lysine(s) targeted by the Rab40b/Cul5 complex. To this end, we incubated purified GST-Rap2a-K2R or GST-Rap2a-K3R with FLAG-Rab40b MDA-MB-231 lysates and assessed binding using GST pull-down assays. Notably, we observe a significant increase in the ability of Rap2a-K3R to bind Rab40b compared to Rap2a-WT (Figure 8E & F), suggesting that the K3R mutant likely contains the most pertinent Rap2 lysine(s) ubiquitylated by Rab40b/Cul5.

Of course, the goal in any ubiquitylation study is to identify the specific lysine responsible for changes in downstream signaling. This can be very difficult, however, as previous studies have shown that ubiquitin machinery can be promiscuous and modify nearby lysines even when single point mutations are made at the site of interest.

Furthermore, our 293T ubiquitylation assays suggest that Rap2 is potentially modified on several lysines. Nonetheless, we split the K3R mutant into four individual mutants: K117R, K148R, K150R, and K148R/K150R. We then used our GST-Rap2a and FLAG-Rab40b binding assay to test whether any of these single (or double) mutations could phenocopy the increased GST-Rap2a-K3R binding to Rab40b (presumably due to lack of ubiquitylation and dissociation of enzyme/substrate complex). Importantly, none of these single mutants recapitulated the increased binding to Rab40b as seen by Rap2a-K5R and Rap2a-K3R (Supplementary Figure 4A & B), thus, we hypothesize that Rab40b/Cul5 may ubiquitylate all three lysines (K117, K148, and K150) within Rap2.

We do note a slight but non-significant increase in Rap2a-K117R binding to FLAG-Rab40b compared to Rap2a-WT (Supplementary Figure 4B). This is consistent with previous reports in *Xenopus*, where K117 was suggested to be a putative Rap2 ubiquitylation site (Lee et al. 2007). Based on these cumulative data, we propose that all three Rap2 lysines K117, K148, and K150 are regulated by the Rab40b/Cul5 complex. In particular, our 293T ubiquitylation experiment strongly suggests that Rab40b stimulates the addition of two-five ubiquitin moieties on Rap2 and while it is possible that all of these ubiquitin modifications are conjugated to one single lysine, it’s also equally possible for multiple Rap2 lysines to be modified, possibly multi-mono-ubiquitylated. Future work will be needed to dissect whether ubiquitylation at these sites (K117, K148, and K150) control distinct downstream signaling or whether all three work in concert with each other.

Moving forward with Rap2a-K3R, we next assessed whether mutation of these three lysines affects localization in a manner similar to the K5R mutant. Thus, MDA-MB-231 cells stably expressing Eos-Rap2a-K3R (Supplementary Figure 3H) were analyzed by immunofluorescence. Like the K5R mutant, we find that Eos-Rap2a-K3R is predominately localized intracellularly, within the endolysosomal compartment (Figure 8G). Fluorescence intensity analysis of Rap2a-K3R compared to Rap2a-WT demonstrates a higher intracellular to whole cell fraction (Figure 8H, Supplementary 2C), phenocopying the effect of Rap2a-K5R. Similar to how Rap2a-WT behaves in Rab40b KO cells, Rap2a-K3R has increased presence in lysosomes (CD63) compared to control cells, but unchanged co-localization with early endosomes (EEA1) (Figure 8I & J). This again supports a model in which ubiquitylation of Rap2 is needed for sorting and recycling to the leading-edge plasma membrane. To test this model further, we asked whether recruitment of Rap2 to the plasma membrane is dependent on Rab40b/Cul5 complex formation. To this end, we examined Rap2 localization in a Rab40b ΔSOCS background (Figure 8L). Rab40b KO cells stably expressing FLAG-Rab40b ΔSOCS were transfected with Eos-Rap2a WT lentivirus and flow sorted (Supplementary Figure 4C). Consistent with our hypothesis, we discover that Eos-Rap2a levels at the plasma membrane are noticeably reduced in Rab40b ΔSOCS cells (Figure 8L). Indeed, our results demonstrate a higher endolysosomal Rap2 fraction in Rab40b ΔSOCS compared to control cells (Figure 8M). Collectively, these data define a model wherein ubiquitylation by the Rab40b/Cul5 complex is required for sorting and recycling of Rap2 to the leading-edge plasma membrane of migrating breast cancer cells.

Although the mis-localization of Rap2 in Rab40b ΔSOCS strongly suggests that the Rab40b/Cul5 complex is necessary for proper Rap2 targeting to the leading-edge (given the lack of binding between Rab40b ΔSOCS and Cul5), we also tested whether direct inhibition of Cul5 resulted in the same phenotype. To this end, we used MLN4924, a small molecular inhibitor of NEDD8-Activating Enzyme. Because CRLs require Cullin neddylation for activity, MLN4924 therefore acts as an inhibitor of CRLs. MDA-MB-231 cells stably expressing GFP-Rap2a were treated with MLN4924 for 24hrs, followed by fixation and staining for CD63. Similar to what we observe in Rab40b KO cells, Rab40b ΔSOCS cells, and Rap2a K-R mutant cells, Rap2 accumulates in lysosomes with MLN4924 treatment (Figure 8N, Supplementary Figure 4D).

Additionally, MLN4924 appears to impact Rap2 enrichment at the leading-edge. Compared to control cells were Rap2 polarization is clear at the lamellipodia plasma membrane, inhibitor treated cells lack Rap2 enrichment at any one location and seem evenly dispersed across the membrane (Supplementary Figure 4D). This suggests that Cul5, and specifically ubiquitylation by Rab40b/Cul5, is needed for dynamic re-distribution of Rap2 at the lamellipodia during cell migration.

Finally, to bridge these findings with Rap2’s putative role in regulating cell movement, we wondered whether the K3R mutation, where Rap2 is lost at the plasma membrane, would have a corresponding effect on cell migration. To this end, we measured the invasiveness of MDA-MB-231 parental cells compared to cells stably expressing either Eos-Rap2a WT or Eos-Rap2a K3R. Remarkably, cells expressing the K3R mutant have decreased invasion overall, suggesting that this mutant perhaps act as a dominant-negative (Figure 8K). Future work will be needed to fully dissect how the K3R mutant adversely affects cell migration. However, what is clear is that modification of these Rap2 lysines are critical for its endosomal trafficking, leading-edge localization, and ultimate function during cell migration and invasion.

### Ubiquitylation of Rap2 at critical lysines is required for its activation

Up until now, we have primarily focused on the role of ubiquitylation in regulating Rap2 localization. The accumulation of Rap2 in the endolysosomal compartment in Rab40b KO and ΔSOCS cells (Figure 6E and Figure 8M) raises an interesting question about the activation state of Rap2 in these cells, given that the Rap2 dominant-negative mutant localizes almost exclusively to endosomes and lysosomes (Figure 5B, Supplementary Figure 1H & I). Additionally, recent work has shown that mono-ubiquitylation of Ras isoforms at K117 and K147 (Rap2-K148 equivalent) directly regulates activation by enhancing intrinsic GDP-to-GTP exchange rates (K117) and inhibiting GAP binding (K147), respectively (Sasaki et al. 2011; Baker et al. 2012, 2013). This raises an interesting possibility that ubiquitylation of Rap2 may also directly regulate its activation.

To investigate this further, we revisited the known Rap2 effector RalGDS and took advantage of a previously described method to measure Rap2 activation status (Spaargaren and Bischoff 1994; Franke, Akkerman, and Bos 1997; Nancy et al. 1999; Ohba, Kurokawa, and Matsuda 2003; Jeon et al. 2007). Specifically, purified GST-RBDRalGDS was incubated with cell lysates expressing various Rap2 mutants and the levels of active Rap2 (bound to GST-RBDRalGDS) was determined by GST pull-down assays followed by western blotting against Rap2. First, as a proof of principle, MDA-MB-231 cells transfected with either GFP-Rap2a-WT, Rap2a-G12V, or Rap2a-S17N were lysed and incubated with GST-RBDRalGDS (Figure 9A). We then defined active Rap2a as the fraction of GFP-Rap2a pulled down by GST-RBDRalGDS compared to the input (GFP-Rap2a levels in the lysate) (Figure 9D). We find that the results are consistent with the constitutively active and dominant-negative mutations, were Rap2a-G12V is more active and Rap2a-S17N is less active than Rap2a-WT, respectively (Figure 9D).

**Figure 9.**
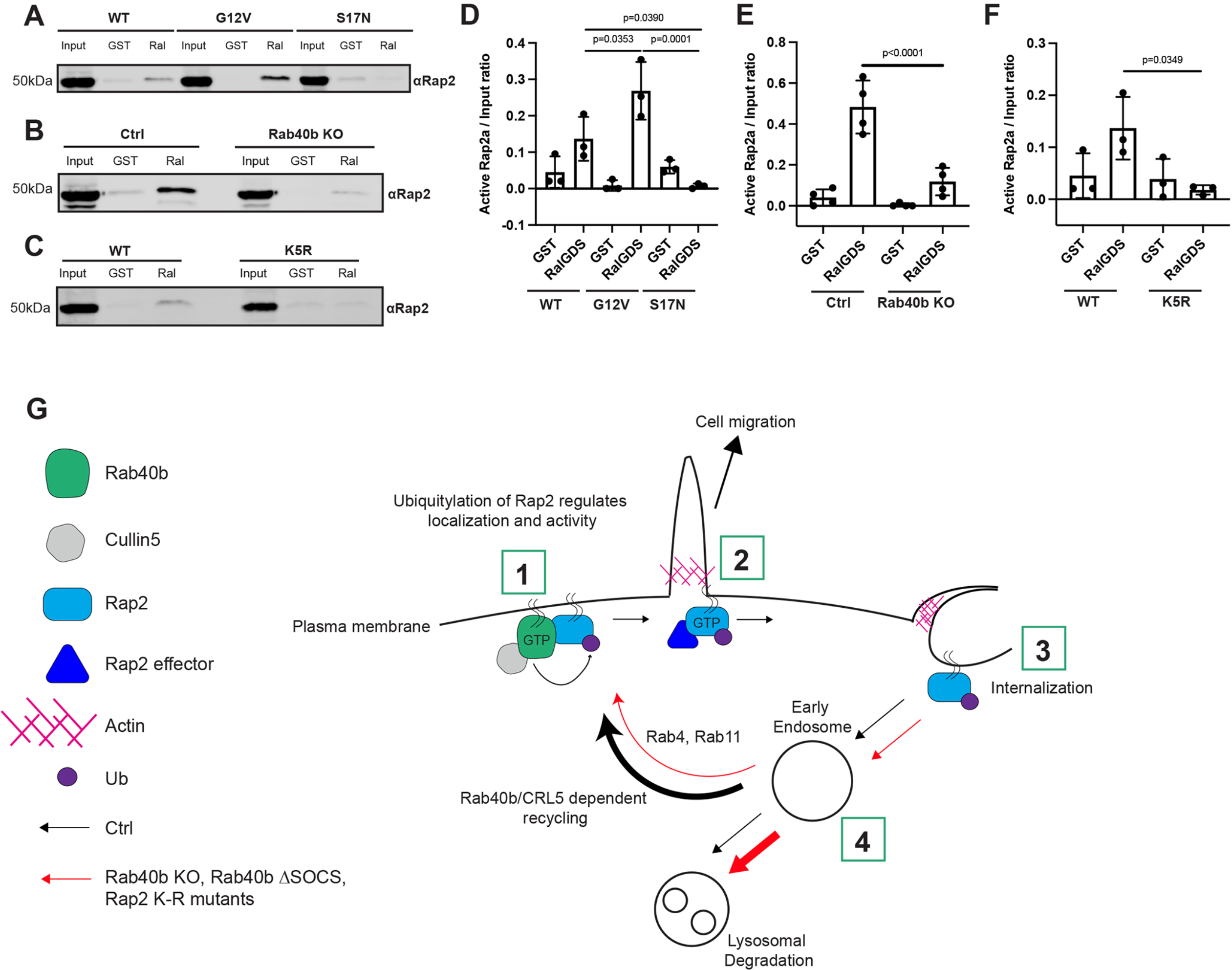
Activation of Rap2 is regulated by Rab40b and is linked to its ubiquitylation. (A) Activation assay Rap2a-WT, -G12V, and -S17N. MDA-MB-231 parental cells were transiently transfected with GFP-Rap2a-WT, -G12V, or -S17N (pLVX-GFP constructs). Lysates were incubated with either GST or GST-RBDRalGDS, followed by a GST pull-down assay. GST alone was used to control for GST binding to Rap2a. 25µg of lysate input was loaded as a positive control and used to calculate active levels of Rap2a. Western blot was probed with αRap2 antibody. Coomassie gel showing equal levels of GST and GST-RBDRalGDS is in Supplementary Figure 5. (B) Activation assay Rap2a-WT in Ctrl vs Rab40b KO cells. MDA-MB-231 parental cells or Rab40b KO cells were transiently transfected with GFP-Rap2a-WT (pEGFP-construct). Lysates were incubated with either GST or GST-RBDRalGDS, followed by a GST pull-down assay. GST alone was used to control for GST binding to Rap2a. 25µg of lysate input was loaded as a positive control and used to calculate active levels of Rap2a. Western blot was probed with αRap2 antibody. Coomassie gel showing equal levels of GST and GST-RBDRalGDS is in Supplementary Figure 5. (C) Activation assay Rap2a-WT and -K5R. MDA-MB-231 parental cells were transiently transfected with GFP-Rap2a-WT, -K5R (pLVX-GFP constructs). Lysates were incubated with either GST or GST-RBDRalGDS, followed by a GST pull-down assay. GST alone was used to control for GST binding to Rap2a. 25µg of lysate input was loaded as a positive control and used to calculate active levels of Rap2a. Western blot was probed with αRap2 antibody. Coomassie gel showing equal levels of GST and GST-RBDRalGDS is in Supplementary Figure 5. (D) Quantification of activation assay in (A). Three biological replicates were performed. Mean ± SD. Active Rap2a/Input ratio was defined as active Rap2 signal/lysate signal. One-way ANOVA with Tukey’s multiple comparisons test. GST signal was not subtracted. WT vs G12V p=0.0353. WT vs S17N p=0.0390, G12V vs S17N p=0.0001. (E) Quantification of activation assay in (B). Three biological replicates were performed. Mean ± SD. Active Rap2a/Input ratio was defined as active Rap2 signal/lysate signal. One-way ANOVA with Tukey’s multiple comparisons test. GST signal was not subtracted. Ctrl vs Rab40b KO p<0.0001. (F) Quantification of activation assay in (C). Three biological replicates were performed. Mean ± SD. Active Rap2a/Input ratio was defined as active Rap2 signal/lysate signal. One-way ANOVA with Tukey’s multiple comparisons test. GST signal was not subtracted. WT vs K5R p=0.0349. (G) Working model for how Rab40b/Cul5 modulates Rap2 subcellular localization and activation via ubiquitylation. 1. Rab40b/Cul5 and Rap2 primarily interact at the plasma membrane, where binding is enhanced when Rab40b is GTP bound. Ubiquitylation by the Rab40b/Cul5 complex causes dissociation. 2. Active Rap2 is then able to interact with its effector proteins at the plasma membrane where it regulates leading-edge actin dynamics and ultimately drives cell migration. 3. To quickly re-distribute Rap2 to other locations at the lamellipodia plasma membrane, Rap2 is internalized via a pinocytosis-like mechanism. Internalization mediates the delivery of Rap2 to early endosomes. 4. Once at early endosomes, previous ubiquitylation by the Rab40b/Cul5 complex serves as a signal for Rap2 to be recycled back to the leading-edge plasma membrane via the Rab4/Rab11 recycling pathways. Without this ubiquitin signal, Rap2 is sorted to lysosomes for degradation. We hypothesize that Rap2 is continuously recycled to and from the plasma membrane to regulate its localization and activity. Ultimately, we propose that dynamic re-distribution of Rap2 is required to maintain its enrichment at the leading-edge, where it can regulate actin dynamics and promote cell migration.

Next, we asked whether Rap2 activity is affected in cells lacking Rab40b. Based on the localization changes detailed in Figure 6, we predicted that Rab40b KO cells would have less active Rap2 overall. Indeed, we detect less active Rap2a in Rab40b KO cells compared to control cells, suggesting that Rab40b, and in particular the Rab40b/Cul5 complex, is required for both Rap2 activation and localization at the plasma membrane (Figure 9B & E). Overall, this data points at a complex system whereby Rap2 subcellular localization and GTP state are interconnected and regulated by Rab40b/Cul5-dependent ubiquitylation.

Delving further, we next questioned whether Rap2a-K5R activation is altered compared to Rap2a-WT. Again, based on our previous observations and recent studies on Ras regulation, we speculated that mutation of these critical lysines would affect GTP activity. Like our previous experiments, we incubated purified GST-RBDRalGDS with lysates expressing Rap2a-K5R and measured levels of GTP-bound Rap2a.

Remarkably, we find that the K5R mutant severely abrogates Rap2 activation, as shown by a lack of Rap2a-K5R binding to GST-RBDRalGDS (Figure 9C & F). This data suggests that ubiquitylation of these lysines is critical for both Rap2 localization and activation.

Interestingly, the K5R mutation appears to have a much stronger negative effect on Rap2 activation compared to the loss of Rab40b, suggesting that the Rab40b/Cul5 complex only ubiquitylates certain lysines within these chosen five and that other E3 ligases are likely also involved in Rap2 regulation. In conclusion, our data hint at a complex system whereby the Rab40b/Cul5 complex co-regulates Rap2 localization and activation. Though it is clear that subcellular distribution and GTP activity are closely intertwined, future investigations will be needed to tease apart these pathways and uncover the direct molecular consequences of ubiquitin modification on Rap2.

## Discussion

One of the most fundamental questions in cell biology is how cells coordinate highly complex processes such as actin polymerization, membrane trafficking, and adhesion remodeling to drive cell migration. In this study, we set out to better characterize the function and regulation of the small GTPase Rap2 during cell migration. Here, we define a novel molecular mechanism by which Rab40b/Cul5-dependent ubiquitylation positively regulates Rap2 subcellular localization, as well as targeting and activation at the leading-edge plasma membrane.

### Rap2 subcellular localization and activation are tightly controlled during cell migration

While emerging evidence from this and earlier studies strongly suggest that Rap2 GTPases are directly involved in promoting cell migration, how Rap2 is regulated has remained essentially unknown. If Rap2 functions by regulating integrin-based adhesion and actin dynamics at the leading-edge, one would expect that cells have fine-tuned mechanisms to control the spatiotemporal dynamics of Rap2 in migrating cells. Tight control of Rap2 localization and activation is likely crucial to maintain highly dynamic lamellipodia formation, extension, and retraction during cell movement. Notably, we find that Rap2 localizes in two primary locations: the lamellipodia plasma membrane and the endolysosomal compartment. This largely aligns with previous reports on Rap1 localization (Pizon et al. 1994; Jeon et al. 2007). Using both fixed and live cell imaging, we discover that Rap2 is dynamically trafficked through the endocytic pathway.

Importantly, we observe that Rap2 is internalized via large macropinosome-like organelles, that eventually become Rab5 positive. Once on Rab5 endosomes (also known as sorting or early endosomes), Rap2 is then sorted between two pathways, Rab4/Rab11 dependent recycling back to the plasma membrane or, lysosomal degradation. Sorting into the Rab4 and Rab11 recycling pathways from early endosomes appears to be a key event that allows rapid and dynamic targeting of internalized Rap2 back to the lamellipodia plasma membrane of migrating cells. Taken together, our collective data suggests a model where localization of Rap2 is determined by active sorting between lysosomes or recycling to the plasma membrane, thus, allowing for dynamic re-distribution of Rap2 as cells migrate in order to maintain its enrichment at the leading-edge (Figure 9G).

Like other small GTPases, Rap2 is active when GTP bound. What is of particular interest is that active Rap2 appears to predominantly localize at the plasma membrane, whereas inactive Rap2 is localized to endosomes and lysosomes. Interestingly, similar observations have been made with Rap1, suggesting that regulation of subcellular distribution may be a key step in modulating activity and function of all Rap subfamily members (Rebstein, Weeks, and Spiegelman 1993; Ohba, Kurokawa, and Matsuda 2003; Bivona et al. 2004; Jeon et al. 2007). Consistent with GTP state and localization being interconnected, we show that Rap2 and its known effector RalGDS co-localize at the plasma membrane. There is some evidence that this is common amongst all Rap2 effectors, however, more work will be needed to uncover additional Rap2 effectors and whether they all localize and function at the plasma membrane (Taira et al. 2004; Machida et al. 2004; Myagmar et al. 2005; Nonaka et al. 2008). Given these cumulative observations, we propose that Rap2 signaling is tightly controlled by regulating its targeting and activation at the leading-edge plasma membrane, and that this is key to driving cell migration. This raises an intriguing question of how cells are able to co-regulate recruitment and activation of Rap2 at the plasma membrane and what molecular machinery is involved.

### The Rab40b/Cul5 complex modulates Rap2 recycling to the leading-edge plasma membrane via ubiquitylation

Post-translational modifications (PTMs) of small monomeric GTPases are known to regulate localization and activity. Often, these modifications are found within the C-terminal hypervariable region (HVR), namely prenylation. Phosphorylation has also been implicated in small GTPase signaling (Shinde and Maddika 2018). Aside from these, ubiquitylation has emerged as an additional layer of signal regulation, especially for the Ras proteins (Jura et al. 2006; Dohlman and Campbell 2019). We wondered whether ubiquitylation of Rap2 might play an important role in regulating its spatiotemporal dynamics and activation in migrating cells.

The Rab family of small GTPases represent the largest branch of the Ras superfamily, governing various membrane transport pathways (Schwartz et al. 2008). Our previous work uncovered the role of one specific Rab, Rab40b, during breast cancer cell migration and led us to also characterize its interaction with the E3-Ubiquitin Ligase scaffold protein Cul5 (A. Jacob et al. 2013; Abitha Jacob et al. 2016; Linklater et al. 2021; Duncan et al. 2021). The Rab40 GTPases (Rab40a, Rab40al, Rab40b, and Rab40c in humans) are unique amongst other Rabs because of their extended C-terminal SOCS box, which facilities binding to Cul5, forming a CRL5 ubiquitylation complex. Importantly, our previous work demonstrates that Rab40b depletion partially phenocopies the cell migration and invasion defects in Rap2 KO cells that we report in this study (Linklater et al. 2021). Furthermore, recent evidence suggested that ubiquitylation of the *Xenopus* ortholog of Rap2 by the XRab40c/Cul5 complex regulates Wnt signaling, though the exact consequence of this ubiquitylation on Rap2 localization, activation, and function remain unclear (Lee et al. 2007). All of these studies combined raise an interesting possibility that Rab40b/Cul5-dependent ubiquitylation may regulate Rap2 localization and activity during mammalian cell migration.

Interestingly, we find that in cells lacking Rab40b, Rap2 is decreased at the plasma membrane and instead accumulates intracellularly, primarily in lysosomes. Consistent with this observation, the levels of Rap2 protein expression decrease with passage number, suggesting that Rap2 undergoes increased lysosomal degradation in Rab40b KO cells (Supplementary Figure 2B). Importantly, we can partially rescue this localization defect with the addition of Rab40b-WT but not with Rab40b-ΔSOCS, a mutant unable to bind Cul5, suggesting that ubiquitylation by the Rab40b/Cul5 complex is necessary for Rap2 recycling to the plasma membrane. Furthermore, we discover the same phenotype for Rap2b and Rap2c, suggesting that at least in MDA-MB-231 cells, all three Rap2 isoforms are regulated by Rab40b.

In an effort to understand how loss of Rab40b/Cul5 binding affects localization, we indeed find that the Rab40b/Cul5 complex mediates direct ubiquitylation of Rap2. Consistent with Rap2 being a substrate of the Rab40b/Cul5 complex, we discover that Rap2 binds Rab40b stronger when Rab40b is unable to bind Cul5, pointing at a lack of Rap2 ubiquitylation and subsequent turnover. The Rab40b-induced ubiquitylation banding pattern we observe is consistent with two to five ubiquitin moieties attached to Rap2. We observe little to no stimulation of Rap2-poly-ubiquitin shmear by Rab40b, leading us to believe that the primary Rab40b-driven signal is the addition of smaller ubiquitin species (two-five). Consistent with this, we find no evidence that Rab40b/Cul5-mediated ubiquitylation of Rap2 signals for proteasomal degradation. Additional studies will be needed to fully define the Rab40b-dependent ubiquitin marks on Rap2 (ie. multi-mono vs poly-linkages). Importantly, these findings contrast with the recent study which suggested that Rab40b strictly mediates poly-ubiquitylation of Rap2 in *Xenopus* embryos. Whether these differences in Rap2 ubiquitylation are the result of different experimental systems (mammalian vs *Xenopus* cells) or different Rap2 functions (cell migration vs Wnt signaling) remains to be determined. Finally, since we see no indication that Rab40b/Cul5 stimulates the first mono-ubiquitin mark on Rap2, it is possible that Rab40b/Cul5 works with other E3s, where Rab40b/Cul5 triggers additional ubiquitylation of Rap2 after it is originally primed with ubiquitin by another E3 Ligase.

### Ubiquitylation at critical lysines within Rap2 promotes its plasma membrane targeting

To begin linking the localization defects of Rap2 in Rab40b mutant cells and our evidence suggesting Rap2 is a substrate of Rab40b/Cul5, we turned to mutation of Rap2 lysines to understand how ubiquitylation regulates Rap2 function. “K-to-R” mutants have been canonically used to study loss of ubiquitylation and the downstream signaling effects. We generated a Rap2a construct with five putative lysines mutated to arginines (K5R, K94R, K117R, K148R, and K150R). Mutation of K5, K94, K148, and K150 were shown to inhibit Nedd4-1 mediated ubiquitylation, whereas mutation of K117 appeared to abrogate ubiquitylation of Rap2 in *Xenopus* cells (Kawabe et al. 2010; Lee et al. 2007). Importantly, three of these lysines are conserved within the Ras family (K5, K117, and K148) (Supplementary Figure 4D). K94 appears to be specific to the Rap2 family and may underlie a unique mechanism to regulate Rap2, whereas K150 is conserved within the Rap1 and Rap2 families (Supplementary Figure 4D). We find that mutation of K5R affects Rap2 subcellular localization in a similar manner to loss of Rab40b. Notably, we observe that Rap2a-K5R is primarily localized within lysosomes. The increased Rap2 in lysosomes suggests that ubiquitylation of Rap2 is required for its recycling to the lamellipodia plasma membrane.

Aside from localization changes, we also note that Rap2a-K5R binds stronger to Rab40b compared to Rap2a-WT. Going back to our earlier findings, we hypothesize that the Rab40b/Cul5 complex may target one or more of these lysines, as mutation prevents dissociation. To begin teasing apart the Rap2a-K5R mutant, we separated the mutations into two clusters: K2R (K5 and K94) and K3R (K117, K148, and K150). We find that the K3R mutant phenocopies the localization and Rab40b binding effect of K5R. Delving further, we split the K3R mutant into single and double mutations.

Interestingly, none of the single/double mutants seem to repeat the Rab40b binding effect we observe with K5R/K3R. K117R does appear to have a slight increase in binding to Rab40b, and while we plan to investigate this further, we ultimately propose that all three lysines within K3R are ubiquitylated and regulated by Rab40b/Cul5.

However, there are clear caveats with using the binding as a read-out for which sites are ubiquitylated and we cannot fully rule out the possibility that these mutants affect Rap2 and Rab40b binding at a structural level and that a more complex mechanism is at play. Future work will be needed to tease apart exactly how each of these three Rap2 lysines are regulated by Rab40b/Cul5 specifically.

### The versatile and complex role of ubiquitylation during protein trafficking

Summarizing so far, we propose that Rab40b/Cul5-mediated ubiquitylation of Rap2 promotes its recycling from early endosomes to the lamellipodia plasma membrane. This is supported by our data demonstrating that mutation of putative Rap2 ubiquitylation sites results in a loss of Rap2 enrichment at the plasma membrane and a concurrent increase in lysosomes. This is quite surprising, as it challenges the current dogma that ubiquitylation of plasma membrane bound proteins leads to lysosomal degradation. Canonically, when proteins at the plasma membrane need to be removed, ubiquitylation serves as an internalization signal, where proteins are then trafficked and sorted into lysosomes for degradation (Mukhopadhyay and Riezman 2007; Piper, Dikic, and Lukacs 2014). Our data suggests that ubiquitylation serves as a targeting signal to recycle Rap2 back to the plasma membrane once internalized. While there is at least one recent example of ubiquitin functioning as a recycling signal rather than degradation, this phenomenon remains elusive and poorly studied (Xu et al. 2017).

Many future questions remain: is Rap2 internalized and recycled by two different types of ubiquitin signals? Does Rab40b/Cul5 tag Rap2 right before internalization? What are the readers responsible for recognizing this recycling ubiquitin mark? Is ubiquitin removed by DUBs before it returns to the plasma membrane to start the cycle again? While future work will be needed to answer these intriguing questions, our study defines a new pathway that regulates localization and function of Rap2, and possibly Rap1, small GTPases during cell migration.

### Differential regulation between Rap and Ras family members

Though the Rap subfamily shares some similarities with the Ras family, our data draw attention to the differences and highlight the need for studying this differential regulation further. First, it is particularly important to compare Rap2 localization with Ras isoform localization: HRas and NRas are localized at both the plasma membrane and the Golgi, whereas KRas predominately sits at the plasma membrane (Hancock 2003). The enrichment and active trafficking through the endolysosomal compartment is a unique feature of the Rap family, suggesting distinct regulation and function compared to Ras proteins. Further, our data propose that Rap2 interacts with its effectors predominately at the plasma membrane, whereas Ras has been shown to signal from the Golgi and the endoplasmic reticulum (Chiu et al. 2002). Second, Rap2 seems to be regulated by ubiquitin differently than the Ras family. For instance, ubiquitin attached to HRas stabilizes its interaction with endosomes (Jura et al. 2006), whereas we observe that Rap2 ubiquitylation positively controls its membrane targeting and signaling. In both cases, it is apparent that cells need continuous trafficking of Rap and Ras for proper signaling, but the mechanism by which ubiquitylation governs these pathways appears unique.

Despite the differences, previous work in the Ras ubiquitylation field allows us to think more critically about our Rap2 findings and future directions (Baker et al. 2012, 2013; Yin et al. 2020; Thurman, Siraliev-Perez, and Campbell 2020). Specifically, mono-ubiquitylation of KRas-K147 impedes GAP-mediated GTP hydrolysis and promotes effector binding (Sasaki et al. 2011; Baker et al. 2012, 2013). Additionally, mono-ubiquitylation of HRas-K117 enhances the intrinsic rate of nucleotide exchange, promoting activation. This site specific ubiquitylation of Ras isoforms appears to differentially regulate downstream signaling. In the case of Rap2, we would argue that three lysines are putative sites for Rab40b/Cul5-mediated ubiquitylation: K117, K148, and K150. Future work will be aimed at uncovering the functions of each site and how this compares to previous Ras findings. Of note, both K148 and K150 are located within the HVR, a common region for PTMs, and K150 is specific to the Rap family.

Ubiquitylation of this residue could be a novel mechanism to differentially regulate the Rap family proteins. Overall, our data helps bolster the paradigm for differential regulation of Rap and Ras GTPases by ubiquitylation.

### The complicated link between Rap2 localization and activation

The similarity in localization between our various Rap2 lysine mutants and the dominant-negative mutant is noteworthy and suggests that ubiquitylation may also be critical for Rap2 GTPase function. As mentioned before, there is already precedence for this, as mutation of Ras isoforms at some of these lysines has been shown to affect GTP hydrolysis and GAP binding (Baker et al. 2012, 2013; Dohlman and Campbell 2019). Notably, we detect less active Rap2 overall in Rab40b KO cells, as well as the K5R mutant. These data argue that ubiquitylation by the Rab40b/Cul5 complex is critical not only for Rap2 recycling to the leading-edge, but also for Rap2 activation. However, it remains to be determined whether this occurs via direct regulation of GTP hydrolysis/GAP binding or simply a consequence of Rap2 not being properly targeted to the plasma membrane, where Rap2 GEFs may reside (Gloerich et al. 2012). Being able to tease apart the complex link between plasma membrane targeting and activation will be the focus of future studies. We also want to note that RalGDS binding assays may not be the best suited for this question, especially since RalGDS appears to also bind other Ras and Rap proteins (Spaargaren and Bischoff 1994; Herrmann et al. 1996). In future studies, it may be beneficial to use an effector more specific to Rap2 as well as optimize conditions for different cell types. Moreover, unlike binding assays where spatiotemporal information is lost, biosensors may be better suited for measuring Rap activation differences.

### The Rab40b/Cul5 complex as a pro-migratory machine

Taken together, our collective data suggests that the Rab40b/Cul5 complex ubiquitylates Rap2 to regulate its plasma membrane targeting and activity, ultimately promoting breast cancer cell migration (Figure 9G). Interestingly, the downstream signaling effect of Rap2 ubiquitylation seems to be dependent on the specific E3 ligase. For instance, Nedd4-1 mediated ubiquitylation of Rap2a inhibits its function and activation, affecting kinase activity and promoting dendrite growth (Kawabe et al. 2010). Similarly, in glioma cells, ubiquitylation of Rap2a by Nedd4-1 inhibits GTP activity, promoting migration and invasion (L. Wang et al. 2017). Uncovering the differences between these ubiquitylation events is critical to understand how Rap2 is fine-tuned during cell migration. With Rab40b/Cul5, we establish its role as a pro-migratory molecular machine, where the complex ubiquitylates and regulates a multitude of proteins, many of which directly mediate actin dynamics. Consistent with this model, we have already shown that Rab40b/Cul5 ubiquitylates and regulates the subcellular distribution of another actin regulator, EPLIN (Linklater et al. 2021). Curiously, Rab40b/Cul5 appears to drive poly-ubiquitylation and degradation of EPLIN, whereas here we find it stimulates recycling of Rap2 via a presumed non-poly ubiquitin mark.

How the same Rab40b/Cul5 complex can mediate such different processes, and to what extent these pathways overlap, are exciting future avenues to investigate. Given that ubiquitylation is a fast-acting post-translational modification pathway, as opposed to transcriptional regulation, this may be an innovative way for cells to quickly change actin and adhesion dynamics at the leading-edge. We are now poised to interrogate a number of potential substrates and understand how ubiquitylation may be important for their function, specifically during cell migration.

## Materials and methods

### Cell culture and generation of lentiviral stable cell lines

MDA-MB-231 cells were cultured in DMEM with 4.5g/L glucose, 5.84g/L L-glutamine, 1% sodium pyruvate, 1% non-essential amino acids, 1µg/mL insulin, 1% penicillin/streptomycin, and 10% fetal bovine serum (FBS). HEK293T cells were cultured in DMEM with 4.5g/L glucose, 5.84g/L L-glutamine, 1% penicillin/streptomycin, and 10% FBS. All GFP-Eos-and FLAG-MDA-MB-231 stable cells lines used in this study were generated using lentivirus transfection (puro). Once lentivirus was collected from 293T cells (see “Transfections” below), virus was transferred to MDA-MB-231 target cells and allowed to incubate for 24hrs before replacing with fresh MDA-MB-231 media. Stable, population cell lines were then selected using 1µg/mL of puromycin. Cell lines were routinely tested for mycoplasma. Additionally, all cell lines were authenticated in accordance with ATCC standards.

### Generating MDA-MB-231 CRISPR/Cas9 KO cell lines

MDA-MB-231 cells stably expressing Tet-inducible Cas9 (Horizon Discovery Edit-R lentiviral Cas9) were grown in a 12-well dish to ∼75% confluency. Cells were then treated with 1µg/mL doxycycline for 24hrs to induce Cas9 expression. After 24hrs, cells were co-transfected with crRNA:tracrRNA mix and DharmaFECT Duo transfection reagent as described by the Horizon Discovery DharmaFECT Duo protocol available online. Transfected cells were incubated for 24hrs, then trypsinized and plated for single clones. Clones were screened through genotyping PCR and sanger sequencing for Rab40b/Rab40 KOs and Rap2 KOs. For Rap2 KOs, original screening was done via western blot using the commercially available pan-Rap antibody. Guide RNAs were purchased from Horizon Discovery (Edit-R Predesigned synthetic crRNA), as well as synthetic tracrRNA (Cat# U-002005-xx). All guide RNAs and genotyping primers are listed in Table 2.

#### CRISPR KO genotyping results

* indicates where the mutant allele sequence diverges from WT due to frameshift <u>Rap2 KO1:</u>

Rap2a allele 1:

*GSGGVGKSALTVQFVTGTFIEKYDPTIEDFYRKEIEVDSSPSVLEILD

Rap2a allele 2:

*GSGGVGKSALTVQFVTGTFIEKYDPTIEDFYRKEIEVDSSPSVLEILD

Rap2b allele 1: MRE*SALTVQFVTGSFIEKYDPTIEDFYRKEIEVDSSPSVLEILDTAGTEQFASMRDLYIK NGQGFILVYSLVNQQSFQDIKPMRDQIIRVKRYERVPMILVGNKVDLEGEREVSYGEGKALAEEWSCPFMETSAKNKASVDELFAEIVRQMNYAAQPNGDEGCCSACVIL

Rap2b allele 2:

MRE*CWARAAWASPRSPCSS

Rap2c allele 1:

*VVLGSGGVGKSALTVQFVTGTFIEKYDPTIEDFYRKEIEVDSSPSVLEILDTAGTEQFA SMRDLYIKNGQGFILVYSLVNQQSFQDIKPMRDQIVRVKRYEKVPLILVGNKVDLEPER EVMSSEGRALAQEWGCPFMETSAKSKSMVDELFAEIVRQMNYSSLPEKQDQCCTTC VVQ

Rap2 KO2

Rap2a allele 1:

*VVLGSGGVGKSALTVQFVTGTFIEKYDPTIEDFYRKEIEVDSSPSVLEILDT

Rap2b allele 1: MREYK*GWCWARAAWASPRSPCSS

Rap2c allele 1:

*REYKVVVLGSGGVGKSALTVQFVTGTFIEKYDPTIEDFYRKEIEVDSSPSVLEILD

Rap2c allele 2:

*REYKVVVLGSGGVGKSALTVQFVTGTFIEKYDPTIEDFYRKEIEVDSSPSVLEILD

Rab40b KO1

Allele 1: MSALGSPVRAYDFLLKFLLVGDSDVGKGEILASLQDGAAESPYGHPAGIDYKTTTILLD GRRVKLQLWDTS*AREDFVPYSAPTPGAHRV

Allele 2: MSALGSPVRAYDFLLKFLLVGDSDVGKGEILASLQDGAAESPYGHPAGIDYKTTTILLD GRRVKLQLWDTSGQGRFCTIFRSYSRGAQGVILVYDIANRWSFDGIDRWIKEIDEHAP

GVPKILVGNRLHLAFKRQVPTEQAQAYAERLGVTFFEVSPLCNFNITESFTELARIVLLR HGMDRLWRPSK*C

Rab40b KO2

Allele 1: MSALGSPVRAYDFLLKFLLVGDSDVGKGEILASLQDGAAESPYGHPAGIDYKTTTILLD GRRVKLQLWDTS*AREDFVPYSAPTPGAHRV

Boyden chamber Matrigel™ invasion assay

MDA-MB-231 control and Rap2 KO cell lines were grown to ∼70% confluency prior to setting up the experiment. Cells were washed with PBS, trypsinized, and resuspended in 0.5% serum DMEM (serum-starved MDA-MB-231 media). While preparing cell suspension, Corning Matrigel™ Invasion Chambers (Corning #354481) were thawed and equilibrated with 500µL (top and bottom) of serum free DMEM for 1hr at 37°C. Next, 750µL of 10% serum DMEM (full MDA-MB-231 media) was added to the bottom of the Matrigel™ Invasion Chambers, and 200,000 cells (in 500µL) were plated in the top chamber (in 0.5% serum DMEM). Cells were incubated for 20hrs at 37°C followed by 4% PFA fixation for 10 min and 0.1% Crystal Violet staining for 10 min. Before fixation, sterile cotton swabs were used to gently scrape and discard non-invaded cells from the inner membrane. After Crystal Violet staining, sterile cotton swabs were again used to soak up excess dye. Matrigel™ inserts were then rinsed with sterile water and allowed to dry overnight, followed by brightfield imaging with a 20X air objective (Nikon Eclipse Ti2 Inverted A1 Confocal). Three biological replicates were performed, with technical duplicates in each set. For each Matrigel™ insert, five fields of view were captured, and cells were counted in Fiji (ten data points per condition, per biological replicate). The graph includes all of the data points, but statistical analysis was performed on the average number of cells/field for each biological replicate.

#### Chemotaxis assay

Chemotaxis cell migration was performed according to the “General IncuCyte Chemotaxis Cell Migration Protocol” available online. First, IncuCyte ClearView 96-Well Plate (Cat. 4582) was coated (top and bottom) with 1X collagen for 1hr at room temperature. MDA-MB-231 control and Rap2 KO cells were grown to ∼70% confluency before being washed with PBS, trypsinized, and resuspended in 0.5% serum DMEM (serum-starved MDA-MB-231 media). Serial dilutions were made in 0.5% serum DMEM so that cell suspension concentrations were ∼17cells/µL. Cells of interest were then seeded at 60µL/per well (1,000 cells total) in the collagen coated IncuCyte ClearView plate. This was based on the recommendation by Essen BioScience, where they found that 1,000 cells per well for adherent cell types was a reasonable starting point. The IncuCyte plate was then placed on a level surface, where cells were allowed to settle for 30 min at room temperature. Finally, the IncuCyte insert with seeded cells was placed into a pre-filled plate with 200µL of chemoattractant per well. The IncuCyte ClearView plate was then imaged every 2hrs for 48hrs using an IncuCyte S3 instrument equipped with a 10x objective (CU Cancer Center Cell Technologies Shared Resource). Three biological replicates were performed, with six technical replicates in each set. The IncuCyte Chemotaxis Analysis Software Module (Cat. 9600-0015) was used to extract “Total Phase Object Area Normalized to Initial Top Value” (ie. sum area of migrated cells normalized to time zero area). The graph shows the raw area (arbitrary units) at each timepoint (averaged between the six technical replicates). We selected the 8hr timepoint as time “zero”, so cells had ample time to adhere to the porous membrane before area measurements were analyzed. This reduced noise in the data and allowed us to express “Relative Chemotactic Migration” as a fold change over the 8hr timepoint. Statistical analysis was performed on the Relative Chemotactic Migration at 48hrs.

#### Transfections

Polyplus jetPRIME transfection reagent (Cat. 114-xx) was used for all MDA-MB-231 transient transfections (aside from DharmaFECT Duo reagent used for generating CRISPR KO cell lines). For a 10cm dish, 7.5µg of DNA was mixed with 500µL of jetPRIME buffer followed by 25µL of jetPRIME transfection reagent. For a 6-well plate, 2µg of DNA was mixed with 200µL of jetPRIME buffer followed by 5µL of jetPRIME transfection reagent Transfection mixture was incubated for 10 min at room temperature and added to cells with full-serum medium. Cells were harvested and/or analyzed 24hrs after transfection. For a more detailed protocol and guidelines on scaling up or down, see protocol available on the Polyplus transfection website. Lipofectamine 2000 transfection reagent was used for all HEK293T transfections (both for lentivirus generation and for ubiquitin experiments).

#### Flow cytometry

For cases where Eos-Rap2a was stably expressed in a background already expressing a lentiviral puromycin resistant construct (Figure 6E & 8M), cell sorting was used to select for Eos positive cells. For Figure 6E & 8M, Rab40b KO cells were first transfected with FLAG-Rab40b WT or ΔSOCS, respectively, generating a lentiviral stable cell line (populational). Next, these FLAG-Rab40b WT/ΔSOCS cells were transfected with Eos-Rap2a WT via lentivirus and flow sorted. Due to low levels of FLAG-Rab40b WT/ΔSOCS and FLAG-antibody sensitivity issues, we were unable to co-stain these cells to show both FLAG and Eos, however, our western blot shows robust Eos-Rap2a signal in both Rab40b WT and Rab40b ΔSOCS cells (Supplementary Figure 1E & 4C). For flow sorting, cells were washed in PBS, trypsinized, and resuspended in sort buffer (PBS containing 30mM HEPES pH 7.4, 1mM EDTA, and 0.1g BSA) after lentivirus transfection. Cells were then sorted on a GFP positive gate using a Beckman Coulter MoFlo XDP100 (CU Cancer Center Flow Cytometry Shared Resource).

#### Immunofluorescence staining

MDA-MB-231 cells were seeded on collagen coated glass coverslips and grown in full media for at least 24hrs. Coverslips used for all experiments were Number 1 thickness. Unless otherwise stated, cells were fixed with 4% paraformaldehyde for 20 min at room temperature and quenched for 5 min at room temperature with quenching buffer (375mg Glycine in 50mL PBS). Cells were blocked for 30 min at room temperature with block buffer containing PBS, 1mL of FBS, 200mg saponin, and 0.1mg bovine serum albumin. Primary antibodies were diluted in block buffer and incubated for 1hr at room temperature. When co-staining for actin, Alexa Fluor^TM^ 568 Phalloidin (Cat. #a12380) was added with primary antibody solution. See all antibody dilutions in Table 1. Cells were washed with block buffer before adding secondary antibodies (also diluted in block) for 30 min at room temperature. Cells were washed again with block buffer before mounting in Vectashield and sealing with nail polish. During last wash, Hoechst stain was used at 1:2000 for 5 min to visualize nuclei.

**Table 1:** Antibodies and reagents.

| **Name** | **Company/Product Number** | **Dilution** |
| --- | --- | --- |
| FLAG M2 | Sigma F3165 | 1:1000 |
| Rap2 | BD Biosciences 610215 | 1:1000 |
| Tubulin | Li-Cor 926-42211 | 1:2500 |
| GM130 | BD Biosciences 610822 | 1:100 |
| CD63 | Gift from Andrew Peden | 1:100 |
| EEA1 | Gift from Andrew Peden | 1:100 |
| Syntaxin13 | Prekeris Lab (Prekeris et al. 1998) | 1:100 |
| Ubiquitin P4D1 | Santa Cruz sc-8017 | 1:200 |
| GFP | Invitrogen A-11122 | 1:100 |
| GAPDH | CST 5174 | 1:1000 |
| EGFR | CST 2232 | 1:1000 |
| Myc 9E10 | Santa Cruz sc-40 | 1:1000 |
| HA | Santa Cruz sc-7392 | 1:1000 |
| FLAG-HRP conjugate | CST 86861 | 1:1000 |
| AffiniPure Goat Anti-Mouse IgG,<br>light chain specific | Jackson ImmunoResearch 211-032-171 | 1:1000 |
| IRDye 680RD Anti-Mouse<br>Secondary | Li-Cor 926-68072 | 1:2500 |
| IRDye 800CW Anti-Rabbit<br>Secondary | Li-Cor 926-32213 | 1:2500 |
| Alexa 568 Phalloidin | Thermo Fisher a12380 | 1:40 |
| Alexa 488 Anti-Mouse secondary | Jackson ImmunoResearch 715-545-150 | 1:100 |
| Alexa 594 Anti-Mouse secondary | Jackson ImmunoResearch 715-585-150 | 1:100 |
| Alexa 594 Anti-Rabbit secondary | Jackson ImmunoResearch 711-585-152 | 1:100 |
| Hoechst 33342 | AnaSpec AS-83218 | 1:2000 |
| jetPRIME transfection reagent | Polyplus 114-xx | N/A |
| DharmaFECT Duo transfection<br>reagent | Horizon Discovery<br>T-2010-xx | N/A |
| Lipofectamine 2000 transfection reagent | Invitrogen 11668027 | N/A |
| tracrRNA | Horizon Discovery U-002005-xx | N/A |
| Bafilomycin A1 | Selleckchem S1413 | 200nM |
| MLN4924 | Selleckchem S7109 | 300nM |
| Protein G Sepharose 4 Fast Flow | Cytiva 17-0618-01 | 50% |
| Bio-Rad Protein Assay Dye Reagent (Bradford method) | Bio-Rad 5000006 | 1:5 |
| Pierce Silver Stain Kit | Thermo Fisher 24612 | N/A |
| Intercept TBS Blocking Buffer | Li-Cor 927-60001 | 1:3 |
| CytoPhos Phosphate Assay | Cytoskeleton BK054 | N/A |

**Table 2:**
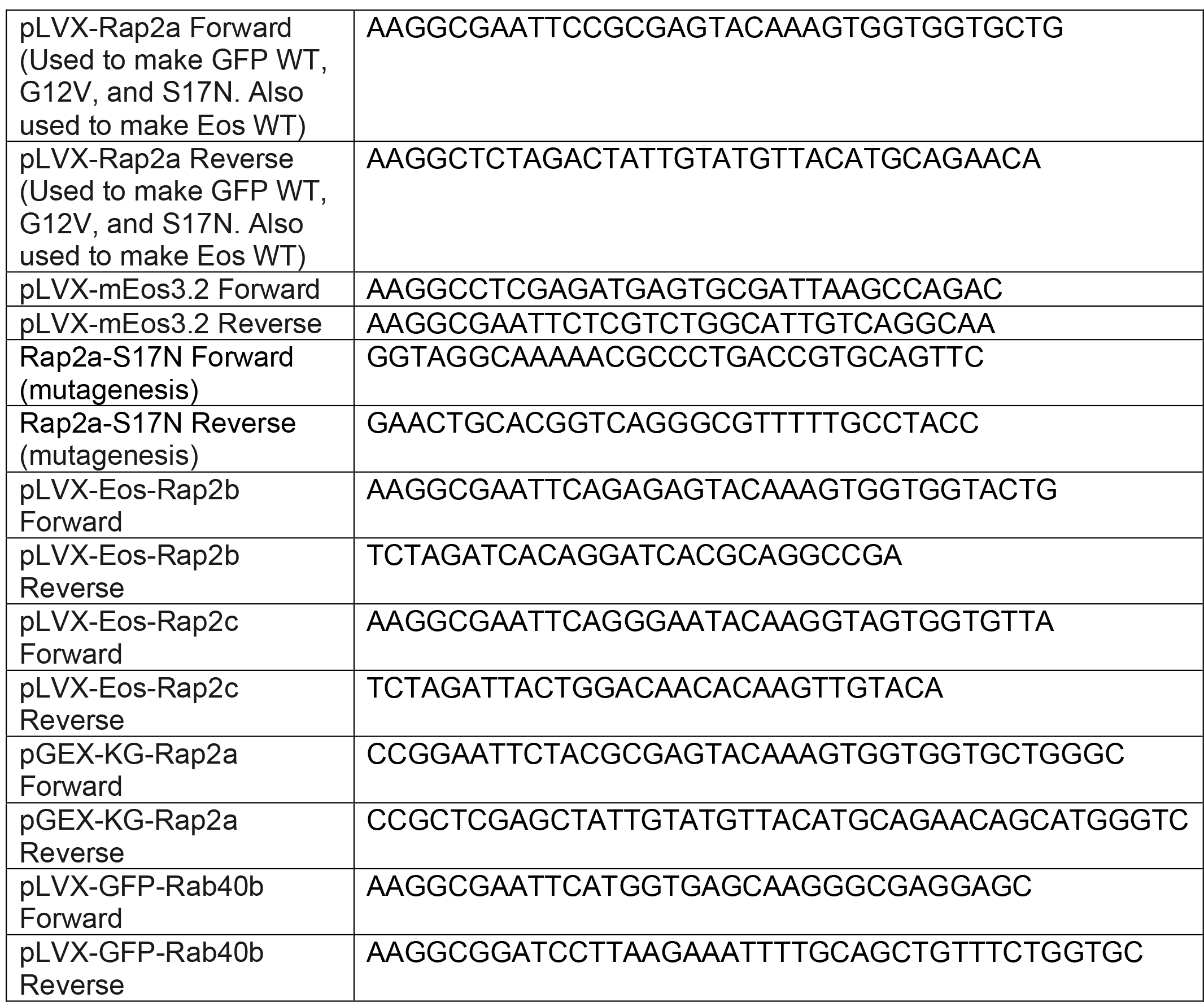
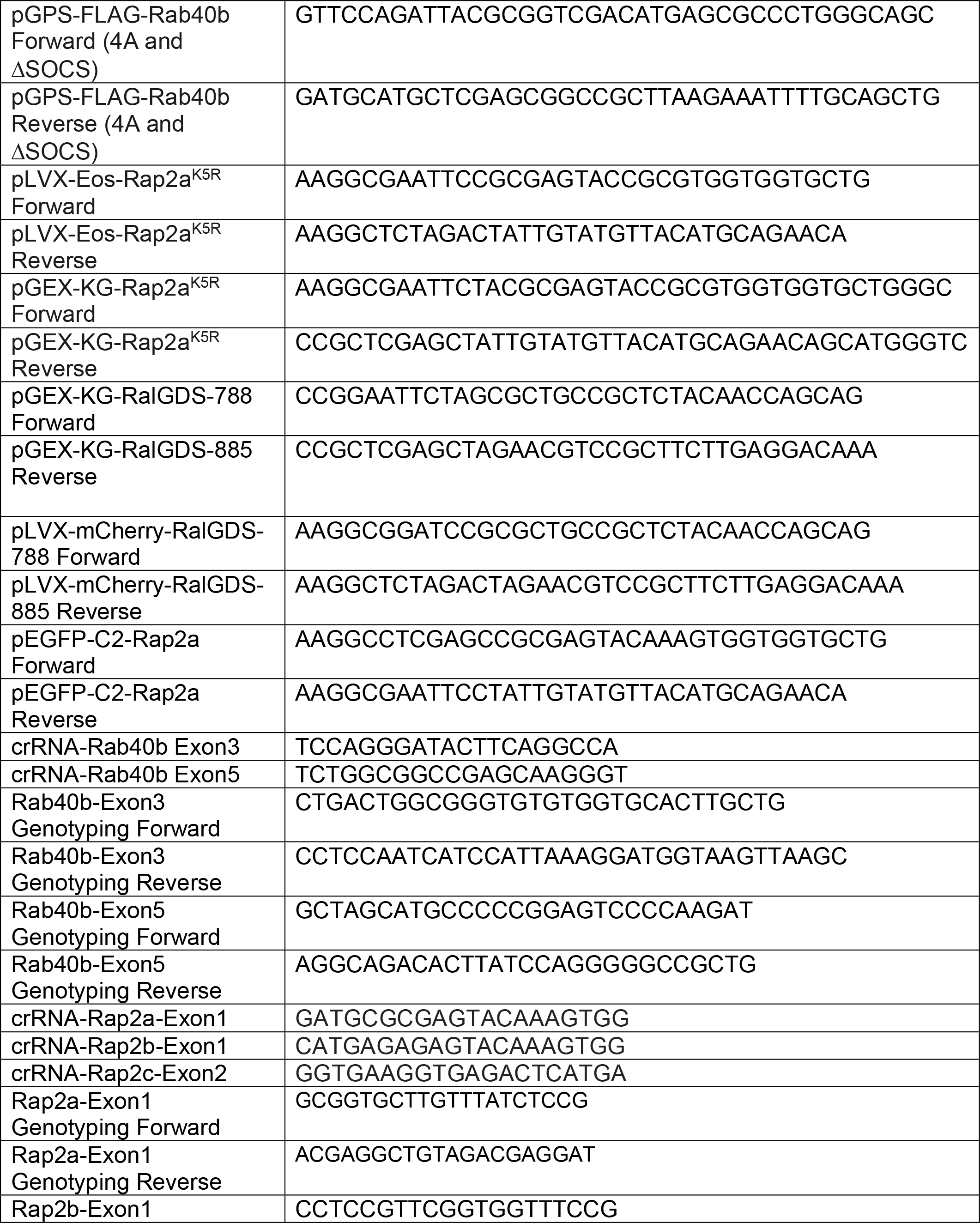
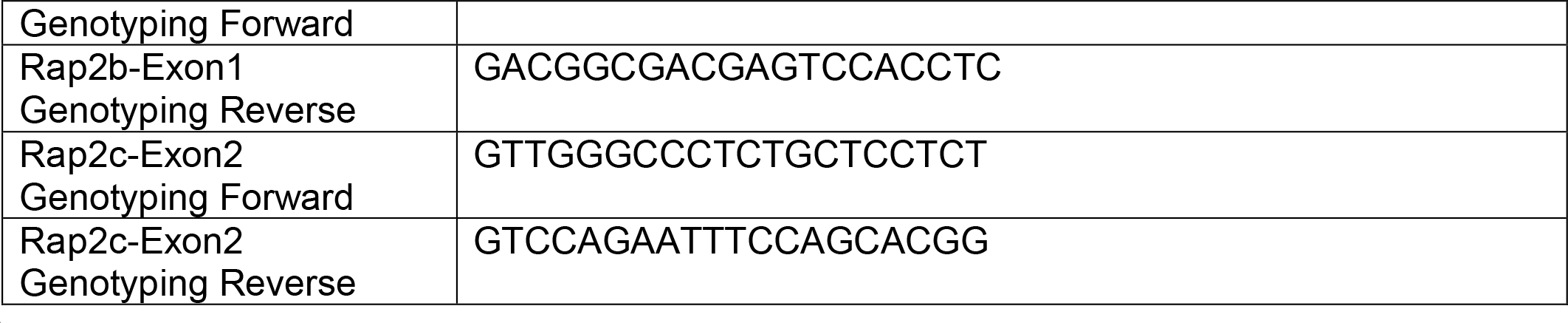
Primer Sequences.

### Microscopy

Unless otherwise noted, all images were acquired using a Nikon Eclipse Ti2 inverted A1 Confocal with a 63x oil objective with a z-step size of 0.5µm. Time-lapse imaging for Videos 1-3 was performed using the same Nikon Eclipse Ti2 inverted A1 Confocal, but with a 40x oil objective equipped with a humidified chamber and temperature-controlled stage. Widefield images (noted in Figure legends) were acquired with an inverted Zeiss Axiovert 200M microscope using a 63x oil objective, QE charge-coupled device camera (Sensicam), and Slidebook 6.0 software (Intelligent Imaging Innovations). Time-lapse imaging for Videos 4-7 was performed on the same Zeiss Widefield, with a 63x oil objective and a temperature-controlled stage. Images were processed in Fiji or 3i software.

### Detailed image analysis for specific experiments

*Figure 1B/C:* Time-lapse imaging was performed using a Nikon Eclipse Ti2 inverted A1 Confocal, with a brightfield 40x oil objective equipped with a humidified chamber and temperature-controlled stage. All time-lapses were taken at 10 min intervals with 10-15 z-stacks (z-step size 0.5µm). Max projections were used for cell tracking (Figure 1B shows max). 95 frames were taken, resulting in a total time-lapse of ∼16hrs. Videos are 5 frames per second. For tracking, cells were manually tracked using the “Manual Tracking” plugin in Fiji. 0.31µm/pixel constant was used to calculate velocity. Careful effort was made to select the geometric center of the cell when manually tracking. Three biological replicates were performed for each cell line (from three passages). In each biological replicate, at least 10 fields of view (xy positions) were imaged. From each of the 10 xy positions, 2-4 random cells were chosen for manual tracking, totaling 30 cells per biological replicate. For each n, the velocity was averaged for all 30 cells. Statistical analysis (unpaired t-test) was performed on the averages of the 3 biological replicates.

*Figure 1I:* To analyze co-localization between GFP-Rap2a and Actin, thresholded Mander’s coefficients (tM) were calculated using the Fiji Coloc 2 plugin. Costes threshold regression was used. PSF 3.0, Costes randomizations 10. Careful effort was made to pick cells without over-exposed signal. One biological replicate was performed, with 5 fields of view and 5 cells in each field (n=25). tM1 was defined as the fraction of Actin overlapping with Rap2a (mean=0.8678) and tM2 was defined as the fraction of Rap2a overlapping with Actin (mean=0.8612).

*Figure 2E & F, Figure 3C, Figure 4G, Figure 6G, Figure 8J & N:* To analyze co-localization between GFP/Eos-Rap2a and endolysosomal markers, 3i SlideBook6 software was used to calculate the percent of total Rap2a that co-localizes with the markers indicated. Briefly, images were fragmented based on fluorescence intensity and masks were created for both the GFP/Eos-Rap2a channel and the Alexa Fluor 594 channel (endolysosomal marker). Then, the percent of total Rap2a intensity within the Alexa Fluor 594 channel mask was calculated for individual cells. Figure 2E: Two biological replicates were performed, with roughly 5 cells imaged for each replicate.

n=10 for GFP-Rap2a/CD63, n=12 for GFP-Rap2a/EEA1, n=10 for Eos-Rap2a/Syntaxin13. Figure 2F: One biological replicate was performed, n=7 for Ctrl cells, n=10 for BafA1 treated cells. Figure 3C: One biological replicate was performed, n=10 for mCherry-Rap2a/YFP-Rab4 WT, n=9 for mCherry-Rap2a/YFP-Rab11a WT. Figure 4G: One biological replicate was performed, n=6 for Ctrl cells, n=8 for 4°C cells, n=7 for Recover cells. Figure 6G: Two biological replicates were performed, with roughly 5 cells imaged for each replicate. n=10 for Ctrl/CD63, n=9 for Rab40b KO/CD63, n=12 for Ctrl/EEA1, n=10 for Rab40b KO/EEA1. Figure 8J: Two biological replicates were performed, with roughly 5 cells imaged for each replicate. n=10 for Ctrl/CD63, n=11 for K3R/CD63, n=12 for Ctrl/EEA1, n=7 for K3R/EEA1. Figure 8N: Three biological replicates were performed, with roughly 12 cells imaged for each replicate. n=33 for Ctrl/DMSO treated cells, n=38 for MLN4924 treated cells.

*Figure 4B:* Line-scans were draw both around the designated GFP-Rap2a organelle (arrow in Figure 4A) and at the lamellipodia plasma membrane. The intensity of GFP-Rap2a in each pixel along these lines was determined with either ImageJ or 3i Slidebook imaging software. The fluorescence intensity of GFP-Rap2a around the organelle line and at the plasma membrane was plotted as fluorescence intensity/area (µ^2^) across 48 seconds.

*Figure 4C:* Same analysis as Figure 4B, but more organelles quantified. Graph includes original organelle from Figure 4A/4B, plus four more organelles from different cells. Criteria for organelle selection was as follows: 1. Organelle must be seen being formed at lamellipodia plasma membrane (this is time 0). 2. Organelle must be visualized in the same plane throughout the entire time-lapse series. So, in sum, only organelles that were seen internalized (time 0) and followed for 48 seconds (end of time-lapse) in the same plane were traced/quantified. As in Figure 4B, line-scans were draw both around the GFP-Rap2a organelles (following criteria above) and at the lamellipodia plasma membrane. The intensity of GFP-Rap2a in each pixel along these lines was determined with either ImageJ or 3i Slidebook imaging software. Instead of plotting the fluorescence intensity/area (µ^2^) across 48 seconds for each organelle, the fluorescence intensity of GFP-Rap2a around the organelle line and at the plasma membrane was graphed as the percent of fluorescence intensity at time 0.

Fluorescence intensity/area (µ^2^) at 0 seconds (start of time-lapse), 24 seconds (half of time-lapse), and 48 seconds (end of time-lapse) was plotted for all 5 organelles as a % of time 0. Statistical analysis (one-way ANOVA) was used to compare fluorescence intensity changes between the plasma membrane and internalized organelles (ie. organelles do decrease in fluorescence intensity over time).

*Figure 4F*: For the cold-block experiment, polarized vs non-polarized cells were defined as follows: Polarized cells have an enrichment of Rap2 at the leading-edge plasma membrane. In other words, cells with a polarized Rap2 distribution somewhere along the lamellipodia were scored as polarized. Everything else was scored as non-polarized. Three biological replicates were performed. 8 fields of view were taken for each n, with roughly 6 cells in each field. Graph shows percent of polarized cells for each condition. Ctrl n1=44 polarized/61 total, n2=30 polarized/42 total, n3=32 polarized/41 total. 4°C n1=4 polarized/41 total, n2=12 polarized/40 total, n3=13 polarized/47 total. Recover n1=30 polarized/40 total, n2=27 polarized/40 total, n3= 37 polarized/44 total.

*Figure 5E:* Fluorescence intensity ratios (leading edge/cell body) were calculated for individual cells under two conditions: MDA-MB-231 cells transiently expressing mCherry-Rap2a + GFP-RBD_RalGDS_ or MDA-MB-231 cells transiently expressing mCherry-Rap2a + Free GFP. The two conditions are separated by the dotted line in the center of the graph. Areas/cells of interest were determined and marked using mCherry-Rap2a localization, choosing cells that had a clearly defined Rap2a enrichment/polarization at one of the cell edges. Then, an area was marked at the leading edge and just behind the leading edge (cell body). Background was subtracted, and sum intensity per area was calculated. For each cell, this was done for both the red channel (mCherry-Rap2a), and the green channel (Free GFP or GFP-RBD_RalGDS_). The ratio of fluorescence at the leading edge versus the area behind the edge was then calculated, giving “enrichment at the leading edge” for each cell. One biological replicate was performed, n=7 cells for GFP-RBD_RalGDS_ condition and n=8 cells for Free GFP condition.

*Figure 5C, 6E, 8B, 8H, and 8M:* To quantify Eos-Rap2a localization changes (ie. decreased Eos-Rap2a at the plasma membrane), intracellular/whole cell fractions were calculated. In Fiji, the images were first split into 3 channels—Eos-Rap2a, Actin, and DAPI. The Eos-Rap2a channel was max projected. Using the z-slices of the Actin channel to find the cell outlines, whole cells were manually traced with the Fiji polygon tool. Individual whole cell traces (including area inside) were defined as ROI-1. Next, another manual trace was made for the inside of the cell (just inside the plasma membrane marked by Actin). Individual intracellular traces (including area inside) were defined as ROI-2. ROI-1 and ROI-2 were individually projected onto the Eos-Rap2a channel. Analyze>Measure was first executed on the corresponding Eos-Rap2a cell, then the ROI was moved to an empty region in the same field of view and executed again (background measurement). This sequence was done for both ROI-1 and ROI-2. Integrated density (area x mean fluorescence intensity) was extracted from Analyze>Measure. To generate intracellular/whole cell fluorescence intensity fractions, background integrated density measurements were first subtracted from ROI-1 and ROI-2. Finally, the intracellular fraction was defined as the fluorescence intensity of ROI-2 (inner cell) divided by the fluorescence intensity of ROI-1 (whole cell) (Supplementary Figure 2C). Two to three biological replicates were performed for each experiment (2-3 coverslips from at least 2 different passages, noted in figure legends). In each biological replicate, 5 fields of view were imaged, with 3 cells analyzed from each field (total of 15 cells per n). Criteria for the 3 cells chosen to analyze were as follows: 1. Clearly defined cell outline 2. No over-exposed signal and 3. Enough empty space in the field of view to acquire a corresponding background measurement. Each cell was treated as its own data point, and statistical analysis (unpaired t-test or one-way ANOVA with post-hoc test) was performed for each condition. The same data points for Eos-Rap2a-WT (in Ctrl cells) were used for Figure 4E, 6C, 6I, and 6K. For each data set, a ROUT outlier test was performed (GraphPad Prism) to identify any outliers. Three outliers were removed from the ‘Eos-Rap2a in Rab40b KO cells’ data set (Figure 4E). No other outliers were found. The same p-value was calculated with and without outliers for Figure 4E, outliers were removed for purposes of graphing.

### Antibodies and reagents

See Table 1 for a list of primary and secondary antibodies as well as specific reagents.

### DNA constructs

YFP-Rap2a was a gift from Johannes L. Bos. Rap2a was cloned from the YFP vector into pLVX-puro with an N-terminal GFP tag to generate pLVX-GFP-Rap2a stable cell lines. Rap2a S17N was generated via site-directed mutagenesis using primers in Table 2 and Rap2a G12V was synthesized by Integrated DNA Technologies (IDT) before cloning into pLVX-puro-GFP (N-term). For Eos-Rap2a constructs, mEos3.2-N1 was purchased from Addgene (#54525) and then cloned into pLVX-puro. Rap2a was amplified from YFP-Rap2a and cloned into pLVX-mEos3.2, generating an N-terminal Eos tag. Rap2a K5R (K5R, K94R, K117R, K148R, and K150R), K3R (K117R, K148R, and K150R), K2R (K5R and K94R), and single K-R mutations (K117R, K148R, K150R, K148R/K150R) were also synthesized by IDT and subsequently cloned into pLVX-puro-Eos or pGEX-KG. For GST-Rap2a, Rap2a was cloned into pGEX-KG (N-term GST tag). GST-Rap2b and GST-Rap2c pGEX-2T plasmids were purchased from Addgene (#55667 and #55668, respectively). For mCherry-Rap2a, mCherry was first cloned into pLVX-puro, then Rap2a was cloned in downstream of mCherry (N-term tag). FLAG-Rab40b WT and FLAG-Rab40b SOCS-4A pLVX-puro constructs were generated as previously described (Linklater et al. 2021). To generate the GFP-Rab40b WT construct, Rab40b WT was amplified from the previously used pLVX-puro-FLAG construct and cloned into pLVX-puro-GFP (N-term). In Supplementary Figure 3B, Rab40b SOCS-4A and Rab40b ΔSOCS were cloned into the FLAG lentivirus vector pGPS: SOCS-4A was amplified from the previous pLVX-FLAG construct, Rab40b ΔSOCS was synthesized by IDT and subsequently cloned into FLAG-pGPS. Rab40b ΔSOCS includes resides 1-174 followed by residues 229-278, so that Rab40b is missing the SOCS box proper, but still contains the C-terminal prenylation site. For RalGDS, GFP-RalGDS was purchased from Addgene (#118315). From this construct, amino acid residues 788-884 as well as residue 885 (incorporated via primer) were cloned into pGEX-KG to generate GST-RalGDS (primers listed in Table 2). This 788-885 stretch includes the defined Ras-binding domain (798-885 based on UniProt KB Q12967). For some of the transient transfections (see Figure legends), Rap2a was cloned into the mammalian expression vector pEGFP-C2 (N-terminal GFP tag). For ubiquitylation experiments in HEK293T cells, the following mammalian expression vectors were used: pRK5-FLAG-Rap2a, pRK5-Myc-Ub, pRK7-HA-Rab40b and pRK7-HA-Rab40b ΔSOCS (all cloned from previous constructs). YFP-Rab4 WT and YFP-Rab11a WT were gifts from Alexander Sorkin. GFP-FIP5-RBD was cloned previously by the Prekeris Lab (Willenborg et al. 2011).

### Cell lysis and western blot analysis

Unless otherwise stated, cells were lysed on ice in buffer containing 20mM HEPES pH 7.4, 150mM NaCl, 1% Triton X-100, and 1mM PMSF. After 30 min, lysates were clarified at 15,000xg in a pre-chilled microcentrifuge. Supernatants were collected and analyzed via Bradford assay (Bio-Rad Protein Assay, Cat. #5000006). 50µg lysate samples were prepared (unless otherwise stated) in 4X SDS loading dye, boiled for 5 min at 95°C, and separated via SDS-PAGE. Gels were transferred onto 0.45µm PVDF membrane (Cat. #IPFL00010), followed by blocking for 30 min in Intercept Blocking Buffer diluted in TBST 1:3 (Cat. #927-60001). Primary antibodies (made in diluted Intercept Blocking Buffer) were incubated overnight at 4°C. The next day, blots were washed in TBST followed by incubation with IRDye fluorescent secondary antibody (diluted Intercept Blocking Buffer) for 30 min at room temperature. Blots were washed once again with TBST before final imaging on a Li-Cor Odyssey CLx. See Table 1 for primary and secondary antibody product numbers and dilutions.

### Cell fractionation

For cell fractionation (Figure 2G), MDA-MB-231 parental cells were grown to confluency (∼5x10cm plates). Cells were then trypsinized and washed in PBS before resuspending the cell pellet in 800µL of 10mM HEPES, pH 7.4 (200µL per plate). Pooled cells were lysed with a Dounce homogenizer (20 strokes) and put on ice before adding NaCl to 100mM final, MgCl2 to 1mM final, and PMSF at 1mM. Cells were then spun at 1,000xg for 5 min at 4°C to sediment unbroken cells and nuclei. Supernatant was moved to an ultracentrifuge tube, and spun at 100,000xg for 1hr in a fixed angle TLS-55 ultracentrifuge (Beckman). Supernatant (cytosol fraction) was collected. The remaining pellet was resuspended in 300µL of ice-cold resuspension buffer (10mM HEPES, pH 7.4, 100mM NaCl, 1mM MgCl2, 1% Triton X-100, 1mM PMSF). Pellet was broken up with a 1cc syringe needle, and then incubated at 4°C for 30 min while rotating.

Suspension was then spun at 100,000xg for 1hr in a fixed angle TLS-55 ultracentrifuge (Beckman). Supernatant (membrane fraction) was collected. The remaining pellet (cytoskeleton fraction) was not collected for this particular experiment. 30µg protein samples were made for both the cytosol and membrane fraction. Samples were separated via SDS-PAGE and western blotted for Rap2, GAPDH (cytosol marker), and EGFR (membrane marker).

### Cold-block experiment

To slow endocytosis (Figure 4E), cells were incubated at 4°C for 60 minutes. To buffer the media at low temperature, 20mM HEPES pH 7.4 was added. For “Recover” condition, cells were incubated at 4°C for 60 minutes with HEPES buffering, then placed back in the 37°C incubator for 40 minutes before fixation.

### Protein expression and purification

GST-Rap2a was purified from BL21-Gold (DE3) competent cells (Cat. #230130). Briefly, cultures were grown at 37°C to OD of ∼0.6, then induced with 0.25mM IPTG final overnight at 16°C. Cells were lysed via French Press in buffer containing 20mM HEPES pH 7.4, 150mM NaCl, 0.1% Tween, 1mM PMSF, 1mM MgCl2, and 1mM 2-Mercaptoethanol. Post-centrifugation lysates were incubated with Glutathione beads for 2hrs at 4°C, then washed with buffer containing 20mM HEPES pH 7.4, 300mM NaCl, and 0.1% Tween. After washing, GST-Rap2a was eluted off the beads using 25mM Glutathione made in buffer containing 20mM HEPES pH 7.4 and 150mM NaCl. Finally, eluted protein was dialyzed into either PBS or 20mM HEPES pH 7.4 and 150mM NaCl. Protein purity was analyzed via SDS-PAGE and protein concentration was measured using Bradford (Bio-Rad protein assay). GST-Rap2a Lysine mutants (K2R, K3R, K5R, single mutants), GST-RalGDS, GST-Rap2b, and GST-Rap2c were purified using the same protocol.

### GST pull-down assays

For GST-Rap2a pull-down assays, MDA-MB-231 cells stably expressing FLAG-Rab40b (WT or 4A) were lysed in buffer containing 20mM HEPES pH 7.4, 150mM NaCl, 1% Triton X-100, 1mM PMSF, and 5mM Iodoacetamide (DUB inhibitor). For GMP-PNP/GDP loading, three steps were followed: 1. Added 5mM EDTA to lysate, incubated 10 min at room temperature, 2. Added 5mM either GMP-PNP/GDP, incubated 10 min at room temperature, 3. Added 15mM MgCl2, incubated 10 min at room temperature.

0.5mg of GMP-PNP/GDP loaded lysate was mixed with 10µg of either GST or GST-Rap2a and incubated for 1hr at room temperature while rotating end-over-end. 75µL of glutathione beads (50% in PBS) were added and incubated for another 30 min at room temperature while rotating end-over-end. Beads were washed 5X with 1mL of buffer containing 20mM HEPES pH 7.4, 300mM NaCl, and 0.1% Triton X-100. Protein was eluted with 1X SDS sample loading dye, separated by SDS-PAGE, and analyzed via western blot. 2/3 of elution was used for western blot, while 1/3 of elution was used for Coomassie to check for presence of GST/GST-Rap2a.

### Competition binding assay

MDA-MB-231 cells stably expressing FLAG-Rab40b WT were lysed in buffer containing 20mM HEPES pH 7.4, 150mM NaCl, 1% Triton X-100, 1mM PMSF, and 5mM Iodoacetamide (DUB inhibitor). Both FLAG-Rab40b WT lysates and GST-Rap2a (recombinant) were loaded with GMP-PNP. For GMP-PNP loading, three steps were followed: 1. Added EDTA to 5mM, incubated 10 min at room temperature, 2. Added GMP-PNP to 5mM, incubated 10 min at room temperature, 3. Added MgCl2 to 15mM, incubated 10 min at room temperature. Once lysates and GST-Rap2a were loaded with GMP-PNP, the following reactions were set up: GST + lysate, GST-Rap2a + lysate, GST-Rap2a + lysate + 5µg RalGDS, GST-Rap2a + lysate + 10µg RalGDS, and GST-Rap2a + lysate + 30µg of RalGDS. In all reactions, GST/GST-Rap2a was at 10µg.

RalGDS was untagged. Reactions were incubated for 1hr at room temperature while rotating end-over-end. 75µL of glutathione beads (50% in PBS) were added and incubated for another 30 min at room temperature while rotating end-over-end. Beads were washed 5X with 1mL of buffer containing 20mM HEPES pH 7.4, 300mM NaCl, and 0.1% Triton X-100. Protein was eluted with 1X SDS sample loading dye, separated by SDS-PAGE, and analyzed via western blot. 1/2 of elution was used for western blot (FLAG), 1/4 of elution was used for Coomassie to check for presence of GST/GST-Rap2a, and remaining 1/4 of elution was used for Silver Stain to analyze RalGDS binding.

### GST-RalGDS activation assay

To measure the levels of active Rap2a, we performed GST pull-down assays with GST-RalGDS (see *DNA constructs* for plasmid details). MDA-MB-231 cells transiently transfected with GFP-or Eos-Rap2a were lysed in buffer containing 20mM HEPES pH 7.4, 150mM NaCl, 1% Triton X-100, 1mM PMSF, and 5mM Iodoacetamide (DUB inhibitor). GMP-PNP/GDP loading was not performed for these assays. 0.5mg of lysate was mixed with 10µg of either GST or GST-RalGDS and incubated for 1hr at room temperature. Following this, 75µL of glutathione beads (50% in PBS) were added and lysate/protein/bead solution was incubated for 30 min at room temperature on a nutator. Beads were washed 5X with 1mL of buffer containing 20mM HEPES pH 7.4, 300mM NaCl, and 0.1% Triton X-100. Protein was eluted with 1X SDS sample buffer, separated by SDS-PAGE, and analyzed via western blot. 2/3 of elution was used for western blot, while 1/3 of elution was used for Coomassie to check for presence of GST/GST-RalGDS.

### GTP hydrolysis assay

CytoPhos Endpoint Phosphate Assay (Cytoskeleton, BK054) was used to measure GTP hydrolysis, specifically the inorganic phosphate (Pi) generated during the enzymatic hydrolysis of GTP. In general, the assay was performed according to the manufacturer protocol. GST-Rap2a-WT and -K5R were purified as described above and prepared in a final buffer of 20mM HEPES pH 7.4 and 150mM NaCl (eg. a non-phosphate buffer). 50µL of 0.4mg/mL protein was first loaded with GTP. For GTP loading, three steps were followed: 1. Added EDTA to 5mM, incubated 10 min at room temperature, 2. Added GTP to 1mM, incubated 10 min at room temperature, 3. Added MgCl2 to 15mM, incubated 10 min at room temperature. Then, 5µL of GTP-loaded protein was added to 25µL of HEPES buffer in a 96-well plate (2µg of protein total). The 96-well plate was then incubated at 37°C for 1hr. After 37°C incubation, 70µL of CytoPhos Reagent was added to each well, resulting in a 100µL reaction. After 10 min of CytoPhos incubation, colorimetric change at 650nm was measured using a plate reader. Blank samples were done side by side (50µL of buffer loaded with GTP).

Absorbance from the blank samples was subtracted from Rap2a positive samples.

Three biological replicates were performed, with three technical replicates in each set. A standard curve for inorganic phosphate (Pi) release was generated using the phosphate standard supplied in the kit. Phosphate release (nmoles) for GST-Rap2a-WT and -K5R was calculated using this standard curve.

### 293T ubiquitin assays

Briefly, 293T cells (∼80% confluency) were transfected with plasmids expressing pRK5-FLAG-Rap2a with or without pRK5-Myc-Ub, pRK7-HA-Rab40b or pRK7-HA-Rab40b ýSOCS using Lipofectamine 2000. After 24hrs, cells were harvest and lysed in 1mL lysis buffer (20mM Tris-HCl, pH 7.6, 150mM NaCl, 2mM EDTA, 1% Triton X-100, 10% glycerol) with 1% SDS. Cell lysates were collected and boiled at 95°C for 10 min.

Supernatants were then diluted with lysis buffer to reduce SDS concentration to 0.1% and incubated with 5µg anti-FLAG M2 antibody and 60µl 50% protein G bead slurry (Cytiva 17-0618-01) overnight. Beads were pelleted by centrifugation and washed three times with lysis buffer plus 0.5 M NaCl. Bound proteins were eluted in 50µl 1X SDS sample buffer. Eluates (20µl) were resolved via SDS-PAGE and transferred to nitrocellulose membranes for immunoblotting. To remove the background IgG heavy chain and light chain after immunoprecipitation, we used an IgG light chain specific secondary antibody (Jackson 211-032-171) and FLAG antibody directly conjugated with HRP (CST 86861S). Blot images were captured using a ChemiDoc MP Imaging system (Bio-Rad).

### Statistical analysis

All statistical analyses were performed using GraphPad Prism Software. Unless described otherwise, statistics were performed using an unpaired student’s t-test (two-tailed) or a one-way ANOVA with post-hoc test assuming normal gaussian distribution. In all cases data were collected from at least from three independent experiments. For microscopy analysis, experiments were performed from at least two different cell passages.

## Supplemental material

Supplemental Figures 1-5 Videos 1-7

## Acknowledgements

We thank Johannes L. Bos for gifting us YFP-Rap2a. We thank the University of Colorado Cancer Center Flow Cytometry Shared Resource and Cell Technologies Shared Resource for their assistance with flow sorting and IncuCyte analysis, respectively (P30CA046934). We are also thankful to members of the Prekeris lab for their continued support and critical feedback. This work was supported by NIH-T32-GM008730 to E.D.D, Bolie Scholarship to E.D.D. and NIH-1R01-GM122768 to R.P.

## Author Contributions

E.D.D. performed experiments with help from K.J.H, M.A.T, and R.P. E.D.D. and R.P. conceived experiments, wrote the manuscript, and secured funding.

---

**Supplemental Figure 1.**
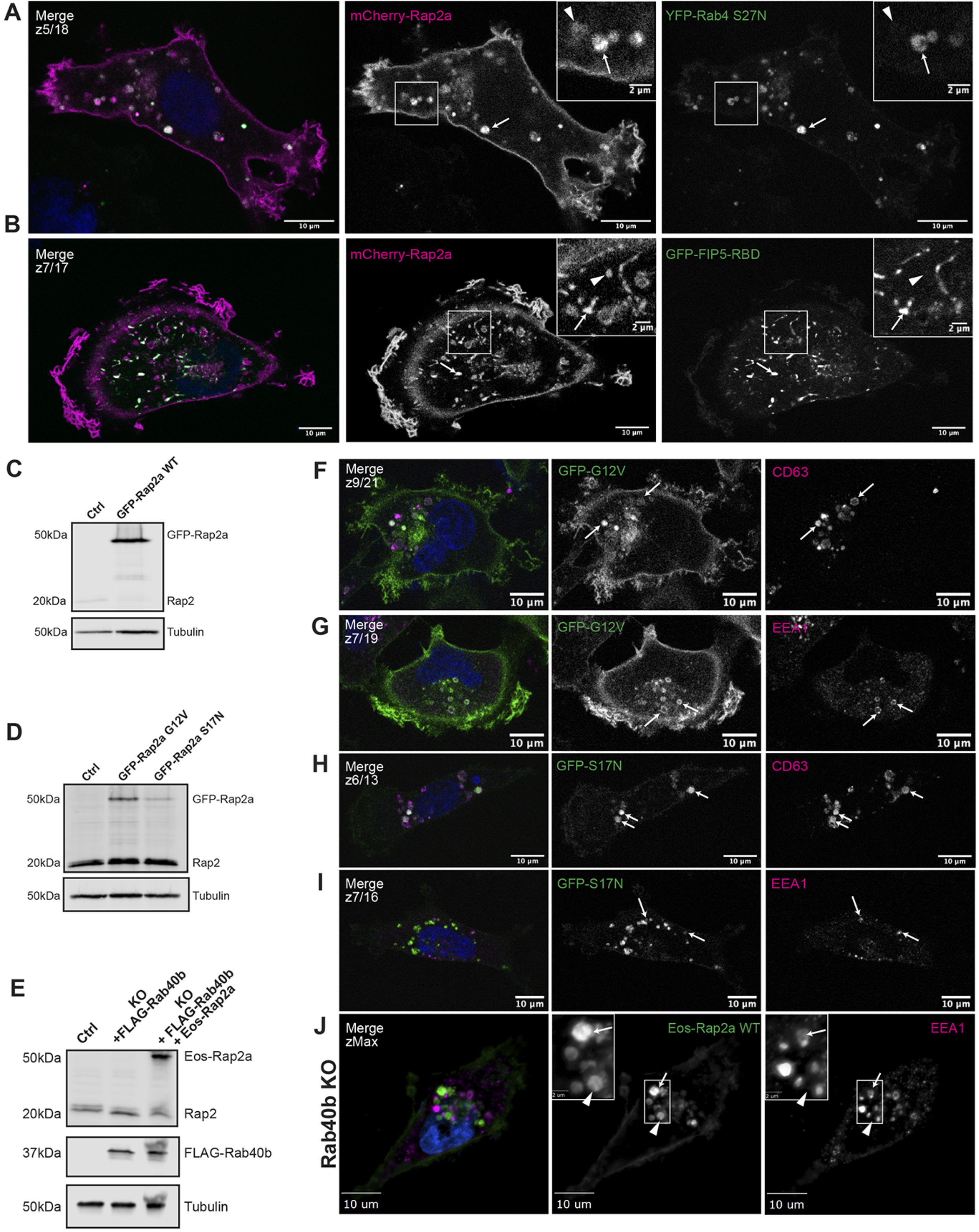
Further characterization of Rap2a subcellular localization in MDA-MB-231 cells and analysis of Eos-Rap2a in Rab40b KO cells. (A) Co-localization of mCherry-Rap2a and YFP-Rab4 S27N. MDA-MB-231 cells were transiently co-transfected with mCherry-Rap2a and YFP-Rab4 S27N, followed by fixation and staining for DAPI (blue). Arrows indicate examples of mCherry-Rap2a and YFP-Rab4 S27N overlap. Arrowheads point to mCherry-Rap2a organelles that are not Rab4 S27N positive. Scale bars = 10µm, 2µm. (B) Co-localization of mCherry-Rap2a and GFP-FIP5-RBD. MDA-MB-231 cells were transiently co-transfected with mCherry-Rap2a and GFP-FIP5-RBD, followed by fixation and staining for DAPI (blue). Arrows indicate examples of mCherry-Rap2a and GFP- FIP5-RBD overlap. Arrowheads point to mCherry-Rap2a organelles that are not FIP5- RBD. Scale bars = 10µm, 2µm. (C) Western blot showing stable overexpression of GFP-Rap2a WT in MDA-MB-231s (lentivirus). Ctrl cells are MDA-MB-231 parentals. 50µg of lysate was loaded for each sample. (D) Western blot showing stable overexpression of GFP-Rap2a-G12V and -S17N in MDA-MB-231s (lentivirus). Ctrl cells are MDA-MB-231 parentals. 50µg of lysate was loaded for each sample. (E) Western blot showing generation of rescue line in Figure 6E. Rab40b KO cells were first stably transfected with FLAG-Rab40b WT (lentivirus, second column). Then, these cells were transfected with Eos-Rap2a and flow sorted (lentivirus, third column, flow sort instead of selection). Ctrl cells are dox-inducible Cas9 MDA-MB-231s that were used to generate Rab40b KO CRISPR line. 50µg of lysate was loaded for each sample. (F) Co-localization of GFP-Rap2a-G12V and CD63. MDA-MB-231 cells stably expressing GFP-Rap2a-G12V were fixed and stained with the lysosomal marker CD63 (magenta) and DAPI (blue). Arrows indicate examples of Rap2a-G12V and CD63 overlap. Scale bars = 10µm. (G) Co-localization of GFP-Rap2a-G12V and EEA1. MDA-MB-231 cells stably expressing GFP-Rap2a-G12V were fixed and stained with the early endosome marker EEA1 (magenta) and DAPI (blue). Arrows indicate examples of Rap2a-G12V and EEA1 overlap. Scale bars = 10µm. (H) Co-localization of GFP-Rap2a-S17N and CD63. MDA-MB-231 cells stably expressing GFP-Rap2a-S17N were fixed and stained with the lysosomal marker CD63 (magenta) and DAPI (blue). Arrows indicate examples of Rap2a-S17N and CD63 overlap. Scale bars = 10µm. (I) Co-localization of GFP-Rap2a-S17N and EEA1. MDA-MB-231 cells stably expressing GFP-Rap2a-S17N were fixed and stained with the early endosome marker EEA1 (magenta) and DAPI (blue). Arrows indicate examples of Rap2a-S17N and EEA1 overlap. Scale bars = 10µm. (J) Eos-Rap2a co-localization with EEA1 in Rab40b KO cells. Rab40b KO MDA-MB-231 cells stably expressing Eos-Rap2a were fixed and stained with the lysosomal marker CD63 (magenta) and DAPI (blue). Widefield microscope. Arrows indicate examples of Eos-Rap2a and EEA1 overlap. Arrowheads point to Eos-Rap2a organelles that are not EEA1 positive. Scale bars = 10µm, 2µm.

**Supplemental Figure 2.**
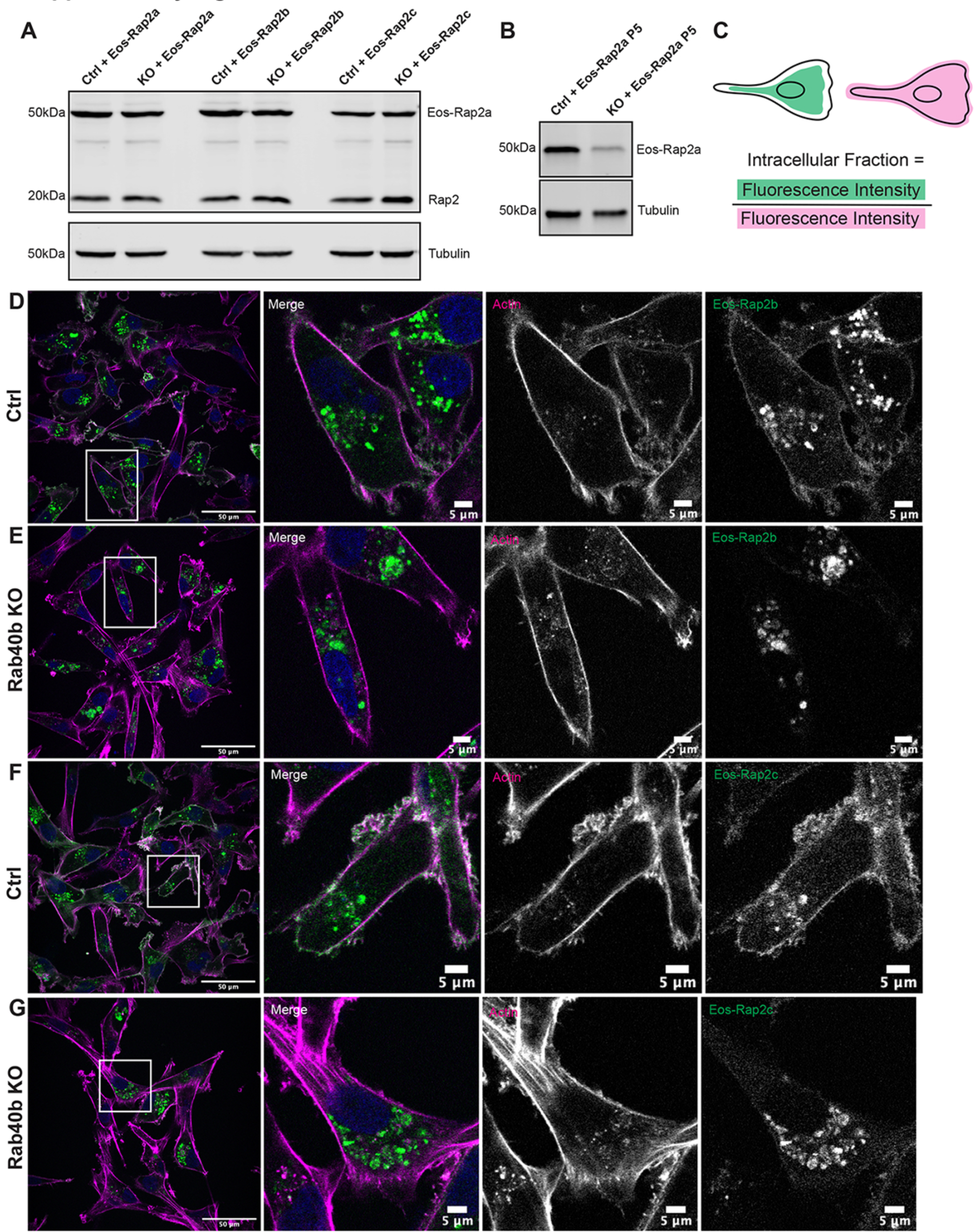
Loss of Rab40b also affects Rap2b and Rap2c subcellular localization. (A) Western blot showing stable overexpression of Eos-Rap2a, Eos-Rap2b, and Eos-Rap2c in MDA-MB-231 Ctrl and Rab40b KO cells (lentivirus). Ctrl cells are MDA-MB-231 parentals. 50µg of lysate was loaded for each sample. (B) Western blot showing stable overexpression of Eos-Rap2a in MDA-MB-231 Ctrl and Rab40b KO cells at passage five (lentivirus). Ctrl cells are MDA-MB-231 parentals. 50µg of lysate was loaded for each sample. (C) Cartoon representation of intracellular/whole cell fraction calculation (see methods). (D) Eos-Rap2b localization in MDA-MB-231 Ctrl cells. MDA-MB-231 cells stably expressing Eos-Rap2b were fixed and stained with Phalloidin (magenta) and DAPI (blue). Scale bars = 50µm, 5µm. (E) Eos-Rap2b localization in Rab40b KO MDA-MB-231 cells. Rab40b KO cells stably expressing Eos-Rap2b were fixed and stained with Phalloidin (magenta) and DAPI (blue). Scale bars = 50µm, 5µm. (F) Eos-Rap2c localization in MDA-MB-231 Ctrl cells. MDA-MB-231 cells stably expressing Eos-Rap2c were fixed and stained with Phalloidin (magenta) and DAPI (blue). Scale bars = 50µm, 5µm. (G) Eos-Rap2c localization in Rab40b KO MDA-MB-231 cells. Rab40b KO cells stably expressing Eos-Rap2c were fixed and stained with Phalloidin (magenta) and DAPI (blue). Scale bars = 50µm, 5µm.

**Supplemental Figure 3.**
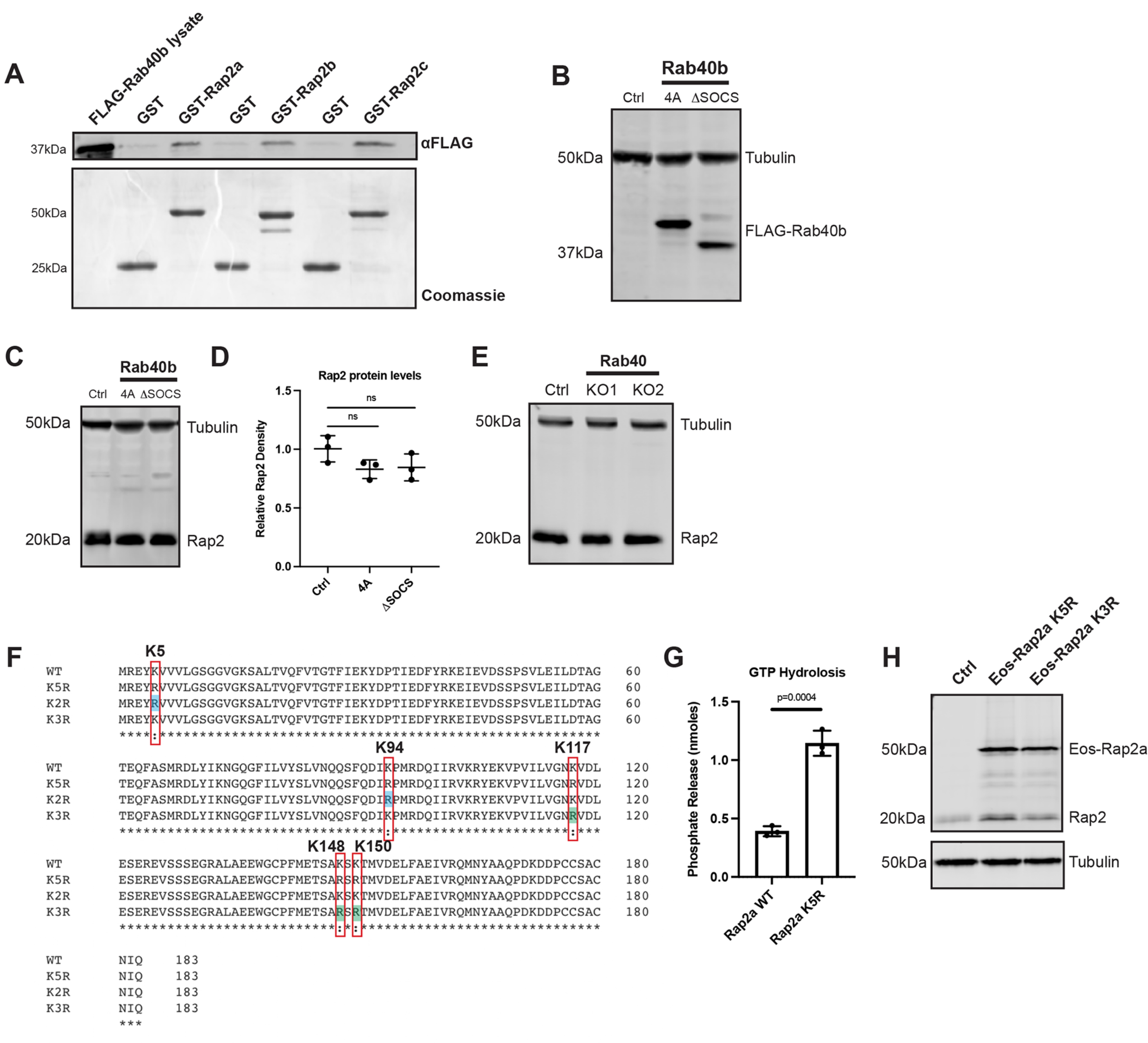
Rap2 protein levels across Rab40 mutant cell lines and generation of Rap2 lysine mutants. (A) Rab40b binding to Rap2a, Rap2b, and Rap2c. MDA-MB-231 lysates stably expressing FLAG-Rab40b WT were incubated with either GST, GST-Rap2a, GST-Rap2b, or GST-Rap2c followed by a GST pull-down assay. 25µg of lysate input was loaded as a positive control and used to estimate pull-down efficiency. Western blot was probed with αFLAG antibody. Coomassie gel shows equal levels of GST and GST-Rap2. (B) Western blot showing stable overexpression of FLAG-Rab40b SOCS-4A and FLAG-Rab40b ΔSOCS in MDA-MB-231s. Ctrl cells are dox-inducible Cas9 MDA-MB-231s that were used to generate CRISPR lines. FLAG-Rab40b SOCS-4A and FLAG-Rab40b ΔSOCS were made in a Rab40b KO background. 50µg of lysate was loaded for each sample. (C) Rap2 protein levels Ctrl vs Rab40b SOCS-4A vs Rab40b ΔSOCS cells. Ctrl, Rab40b SOCS-4A, and Rab40b ΔSOCS cell lysates were probed for αRap2 and αTubulin (loading control). Ctrl cells are dox-inducible Cas9 MDA-MB-231s that were used to generate CRISPR lines. FLAG-Rab40b SOCS-4A and FLAG-Rab40b ΔSOCS were made in a Rab40b KO background. 50µg of lysate was loaded for each sample. (D) Quantification of western blot in (D). Three biological replicates were performed. Relative intensity of Rap2 was normalized to the levels of Tubulin and to Ctrl cells. Mean ± SD. One-way ANOVA with Tukey’s multiple comparisons test. ns=non-significant. (E) Rap2 protein levels Ctrl vs Rab40 KO cells. Ctrl and Rab40 KO cell lysates (lacking Rab40a, Rab40b, and Rab40c) were probed for αRap2 and αTubulin (loading control). Ctrl cells are dox-inducible Cas9 MDA-MB-231s that were used to generate CRISPR lines. 50µg of lysate was loaded for each sample. (F) Protein sequence alignment of Rap2a-WT, K5R, K3R, and K2R. Red boxes denote the five lysines within K5R, blue shades represent K2R, green shades represent K3R. Alignment made using Clustal Omega. (G) GTP Hydrolysis assay Rap2a-WT vs Rap2a-K5R. CytoPhos assay was performed using purified GST-Rap2a-WT and GST-Rap2a-K5R. Colorimetric change at 650nm was measured and analyzed against a standard curve to determine inorganic phosphate release (nmoles). Three biological replicates were performed. Mean ± SD. Unpaired t-test. WT vs K5R p=0.0004. (H) Western blot showing stable overexpression of Eos-Rap2a-K5R and -K3R in MDA-MB-231s (lentivirus). Ctrl cells are MDA-MB-231 parentals. 50µg of lysate was loaded for each sample.

**Supplemental Figure 4.**
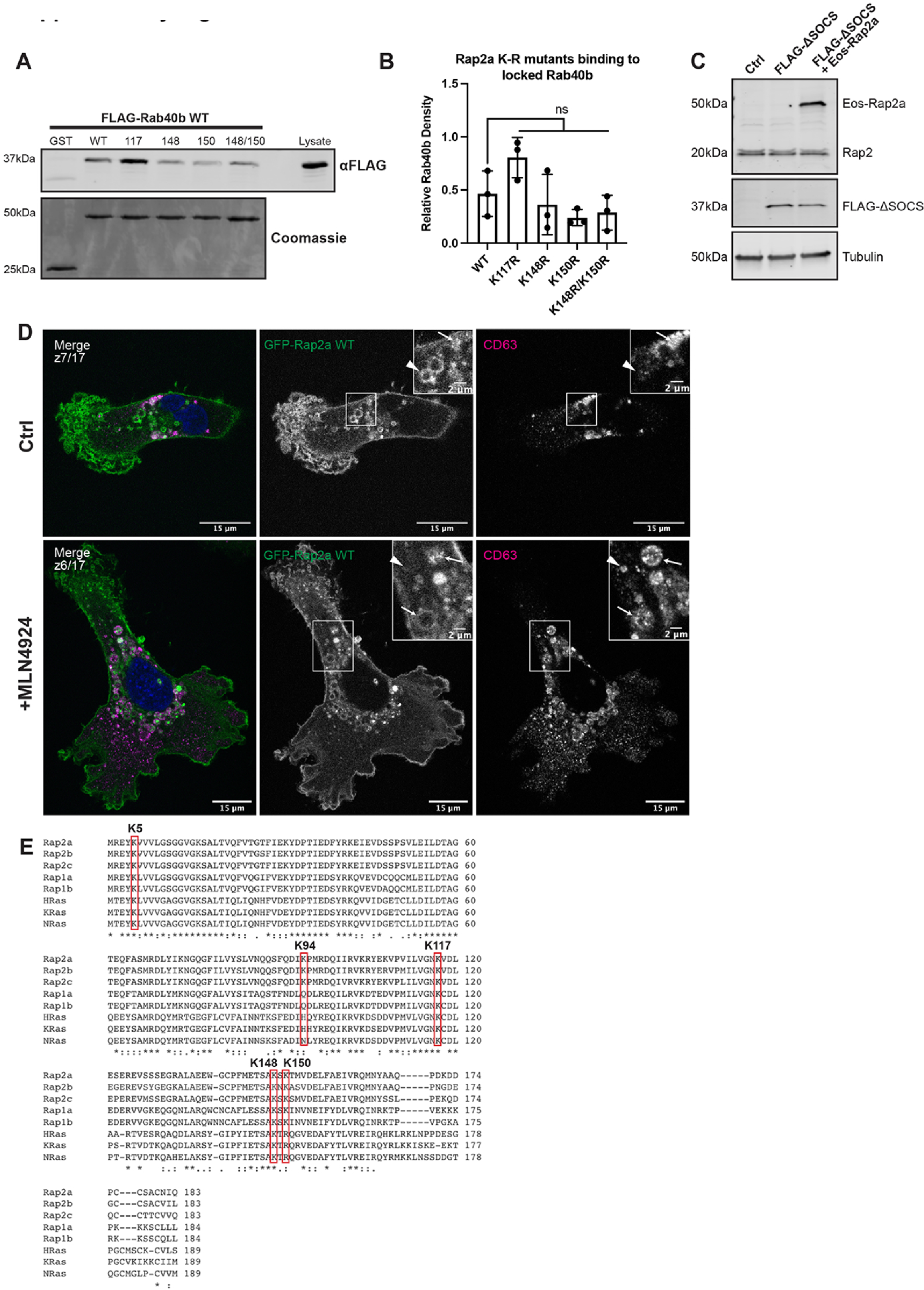
Characterization of Rap2 lysine mutants and the role of Cul5 in mediating Rap2 plasma membrane targeting. (A) Rap2a K-R single/double mutants binding to Rab40b. MDA-MB-231 lysates stably expressing FLAG-Rab40b WT were incubated with either GST, GST-Rap2a-WT or - K117R, K148R, K150R, K148R/K150R, followed by a GST pull-down. Before incubation, FLAG-Rab40b lysates were loaded with GMP-PNP. GST alone was used to control for GST binding to Rab40b. 15µg of lysate input was loaded as a positive control and used to estimate pull-down efficiency. Western blot was probed with αFLAG antibody. Coomassie gel shows equal levels of GST, GST-Rap2a-WT and -K-R mutants. (B) Quantification of GST pull-down in (A). Three biological replicates were performed. Mean ± SD. GST signal was subtracted from experimental lanes, and “Relative Rab40b Density” was calculated by normalizing to lysate. One-way ANOVA with Tukey’s multiple comparisons test. ns=non-significant between WT and all individual mutants. (C) Western blot showing generation of cell line in Figure 8L/M. Rab40b KO cells were first stably transfected with FLAG-Rab40b ΔSOCS (lentivirus, second column). Then, these cells were transfected with Eos-Rap2a and flow sorted (lentivirus, third column, flow sort instead of selection). Ctrl cells are dox-inducible Cas9 MDA-MB-231s that were used to generate Rab40b KO CRISPR line. 50µg of lysate was loaded for each sample. (D) Representative images from MLN4924 experiment in Figure 8N. MDA-MB-231 cells stably expressing GFP-Rap2a were treated with either DMSO (Ctrl) or 300nM MLN4924 for 24 hrs. Cells were then fixed and stained for CD63 (magenta) and DAPI (blue). Arrows indicate examples of GFP-Rap2a and CD63 overlap. Arrowheads point to GFP-Rap2a organelles that are not CD63 positive. Scale bars = 15µm, 2µm. (E) Protein sequence alignment of Rap2a, Rap2b, Rap2c, Rap1a, Rap1b, HRas, KRas, and NRas. Red boxes denote the five lysines within Rap2a-K5R. Stars indicate fully conserved residues. Alignment made using Clustal Omega.

**Supplemental Figure 5.**
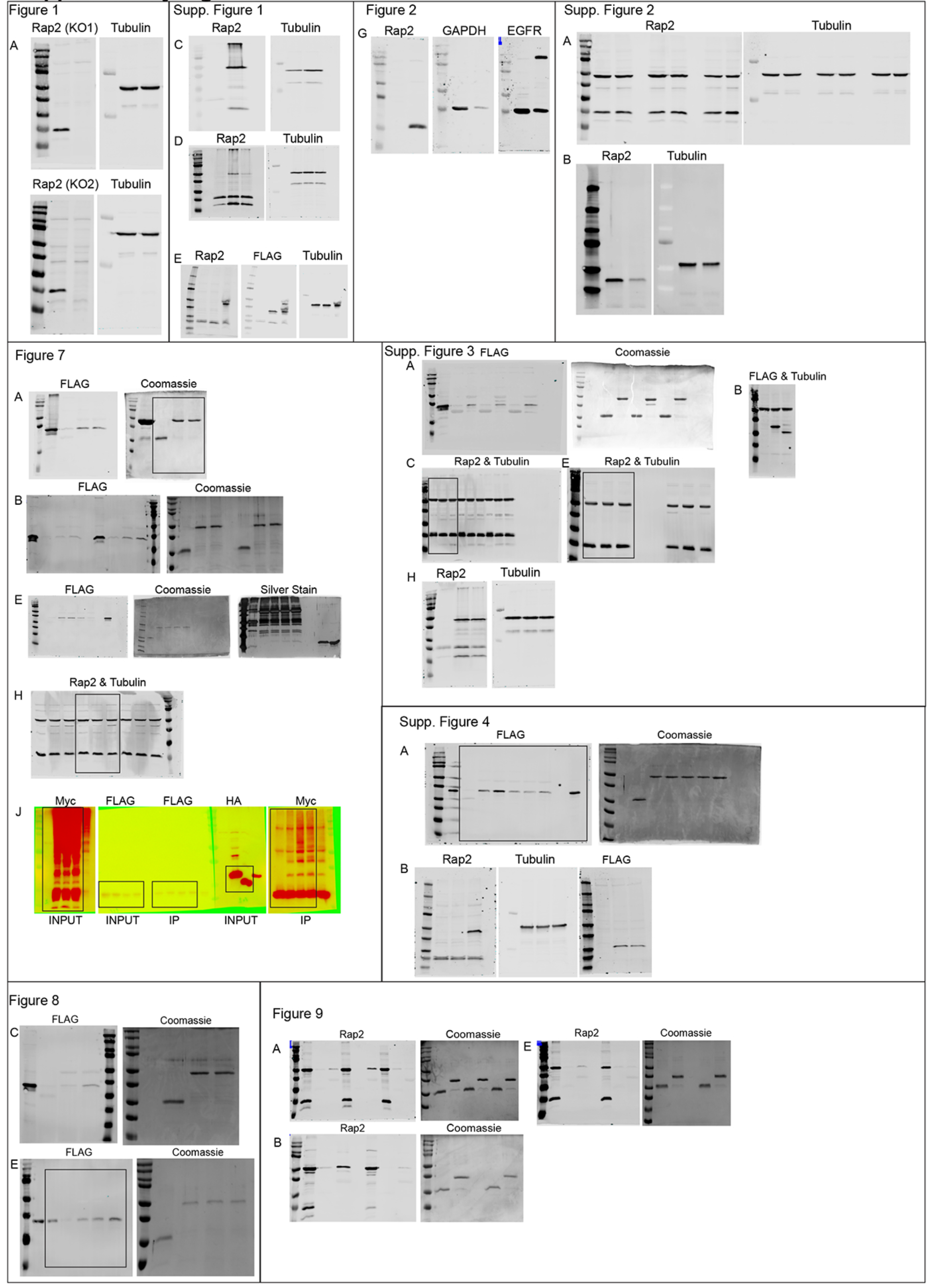
Uncropped western blots and Coomassie gels from all figures.

Video 1. MDA-MB-231 Ctrl 2D cell migration

Live 2D migration on collagen coated glass plate. Ctrl cells are dox-inducible Cas9 MDA-MB-231 cells used to generate the Rap2 CRISPR lines. 10 min intervals, 95 frames. 5fps. Scale bar= 50µm.

Video 2. MDA-MB-231 Rap2 KO1 2D cell migration

Live 2D migration on collagen coated glass plate. MDA-MB-231 Rap2 KO1 cells. 10 min intervals, 95 frames. 5fps. Scale bar= 50µm.

Video 3. MDA-MB-231 Rap2 KO2 2D cell migration

Live 2D migration on collagen coated glass plate. MDA-MB-231 Rap2 KO1 cells. 10 min intervals, 95 frames. 5fps. Scale bar= 50µm.

Video 4. GFP-Rap2a live dynamics

Live 2D time-lapse imaging of MDA-MB-231 cells stably expressing GFP-Rap2a. 1 second intervals (plus exposure time), 100 frames. 5fps. Nearest neighbor deconvolution. Widefield microscope. Scale bar= 10µm.

Video 5. GFP-Rap2a live dynamics continued

Same set up as Video 4, providing additional cell. Live 2D time-lapse imaging of MDA-MB-231 cells stably expressing GFP-Rap2a. 5 second intervals (plus exposure time), 51 frames. 5fps. Nearest neighbor deconvolution. Widefield microscope. Scale bar= 10µm.

Video 6. GFP-Rap2a and mCherry-Rab5 live dynamics

Live 2D time-lapse imaging of MDA-MB-231 cells stably expressing GFP-Rap2a, transiently transfected with mCherry-Rab5. 5 second intervals (plus exposure time), 100 frames. 5fps. Nearest neighbor deconvolution. Widefield microscope. Scale bar= 5µm.

Video 7. mCherry-Rap2a and GFP-RBD_RalGDS_ live dynamics

Live 2D time-lapse imaging of MDA-MB-231 cells transiently expressing mCherry-Rap2a and GFP-RBD_RalGDS_. 3 second intervals (plus exposure time), 50 frames. 5fps. Nearest neighbor deconvolution. Widefield microscope. Scale bar= 10µm.

## References

1. Akutsu, Masato, Ivan Dikic, and Anja Bremm. 2016. “Ubiquitin Chain Diversity at a Glance.” Journal of Cell Science 129 (5): 875–80.

2. Asha, H., N. D. de Ruiter, M. G. Wang, and I. K. Hariharan. 1999. “The Rap1 GTPase Functions as a Regulator of Morphogenesis in Vivo.” The EMBO Journal 18 (3): 605–15.

3. Baker, Rachael, Steven M. Lewis, Atsuo T. Sasaki, Emily M. Wilkerson, Jason W. Locasale, Lewis C. Cantley, Brian Kuhlman, Henrik G. Dohlman, and Sharon L. Campbell. 2012. “Site-Specific Monoubiquitination Activates Ras by Impeding GTPase-Activating Protein Function.” Nature Structural & Molecular Biology 20 (1): 46–52.

4. Baker, Rachael, Emily M. Wilkerson, Kazutaka Sumita, Daniel G. Isom, Atsuo T. Sasaki, Henrik G. Dohlman, and Sharon L. Campbell. 2013. “Differences in the Regulation of K-Ras and H-Ras Isoforms by Monoubiquitination *.” The Journal of Biological Chemistry 288 (52): 36856–62.

5. Bivona, Trever G., Heidi H. Wiener, Ian M. Ahearn, Joseph Silletti, Vi K. Chiu, and Mark R. Philips. 2004. “Rap1 Up-Regulation and Activation on Plasma Membrane Regulates T Cell Adhesion.” The Journal of Cell Biology 164 (3): 461–70.

6. Boettner, Benjamin, and Linda Van Aelst. 2009. “Control of Cell Adhesion Dynamics by Rap1 Signaling.” Current Opinion in Cell Biology 21 (5): 684–93.

7. Bos, J. L., K. de Bruyn, J. Enserink, B. Kuiperij, S. Rangarajan, H. Rehmann, J. Riedl, J. de Rooij, F. van Mansfeld, and F. Zwartkruis. 2003. “The Role of Rap1 in Integrin-Mediated Cell Adhesion.” Biochemical Society Transactions 31 (1): 83–86.

8. Bos, Johannes L. 2005. “Linking Rap to Cell Adhesion.” Current Opinion in Cell Biology 17 (2): 123–28.

9. Bravo-Cordero, Jose Javier, Louis Hodgson, and John Condeelis. 2012. “Directed Cell Invasion and Migration during Metastasis.” Current Opinion in Cell Biology 24 (2): 277–83.

10. Bruurs, Lucas J. M., and Johannes L. Bos. 2014. “Mechanisms of Isoform Specific Rap2 Signaling during Enterocytic Brush Border Formation.” PloS One 9 (9): e106687–9.

11. Chang, Yu-Chiuan, Jhen-Wei Wu, Yi-Chi Hsieh, Tzu-Han Huang, Zih-Min Liao, Yi-Shan Huang, James A. Mondo, Denise Montell, and Anna C. C. Jang. 2018. “Rap1 Negatively Regulates the Hippo Pathway to Polarize Directional Protrusions in Collective Cell Migration.” CellReports 22 (8): 2160–75.

12. Chiu, Vi K., Trever Bivona, Angela Hach, J. Bernard Sajous, Joseph Silletti, Heidi Wiener, Ronald L. Johnson 2nd, Adrienne D. Cox, and Mark R. Philips. 2002. “Ras Signalling on the Endoplasmic Reticulum and the Golgi.” Nature Cell Biology 4 (5): 343–50.

13. Choi, Sun-Cheol, and Jin-Kwan Han. 2005. “Rap2 Is Required for Wnt/Beta-Catenin Signaling Pathway in Xenopus Early Development.” The EMBO Journal 24 (5): 985–96.

14. Coppola, Ugo, Filomena Ristoratore, Ricard Albalat, and Salvatore D’Aniello. 2019. “The Evolutionary Landscape of the Rab Family in Chordates.” Cellular and Molecular Life Sciences: CMLS 76 (20): 4117–30.

15. Dart, Anna E., Gary M. Box, William Court, Madeline E. Gale, John P. Brown, Sarah E. Pinder, Suzanne A. Eccles, and Claire M. Wells. 2015. “PAK4 Promotes Kinase-Independent Stabilization of RhoU to Modulate Cell Adhesion.” The Journal of Cell Biology 211 (4): 863–79.

16. Di, Jiehui, Hui Huang, Debao Qu, Juangjuan Tang, Wenjia Cao, Zheng Lu, Qian Cheng, et al. 2015. “Rap2B Promotes Proliferation, Migration, and Invasion of Human Breast Cancer through Calcium-Related ERK1/2 Signaling Pathway.” Scientific Reports, July, 1–11.

17. Dohlman, Henrik G., and Sharon L. Campbell. 2019. “Regulation of Large and Small G Proteins by Ubiquitination.” The Journal of Biological Chemistry 294 (49): 18613– 23.

18. Duncan, Emily D., Ezra Lencer, Erik Linklater, and Rytis Prekeris. 2021. “Methods to Study the Unique SOCS Box Domain of the Rab40 Small GTPase Subfamily.” Methods in Molecular Biology 2293: 163–79.

19. Feig, Larry A. 1999. “Tools of the Trade: Use of Dominant-Inhibitory Mutants of Ras-Family GTPases.” Nature Cell Biology 1 (2): E25–7.

20. Feig, Larry A., and Geoffrey M. Cooper. 1988. “Inhibition of NIH 3T3 Cell Proliferation by a Mutant Ras Protein with Preferential Affinity for GDP.” Am Soc Microbiol, January. https://mcb.asm.org/content/8/8/3235.short.

21. Franke, B., J. W. Akkerman, and J. L. Bos. 1997. “Rapid Ca2+-Mediated Activation of Rap1 in Human Platelets.” The EMBO Journal 16 (2): 252–59.

22. Franz, Clemens M., Gareth E. Jones, and Anne J. Ridley. 2002. “Cell Migration in Development and Disease.” Developmental Cell 2 (2): 153–58.

23. Gibbs, J. B., M. D. Schaber, T. L. Schofield, E. M. Scolnick, and I. S. Sigal. 1989. “Xenopus Oocyte Germinal-Vesicle Breakdown Induced by [Val12]Ras Is Inhibited by a Cytosol-Localized Ras Mutant.” Proceedings of the National Academy of Sciences 86 (17): 6630–34.

24. Gimple, Ryan C., and Xiuxing Wang. 2019. “RAS: Striking at the Core of the Oncogenic Circuitry.” Frontiers in Oncology 9 (September): 965.

25. Gloerich, Martijn, Jean Paul ten Klooster, Marjolein J. Vliem, Thijs Koorman, Fried J. Zwartkruis, Hans Clevers, and Johannes L. Bos. 2012. “Rap2A Links Intestinal Cell Polarity to Brush Border Formation.” Nature Cell Biology, July, 1–17.

26. Hancock, John F. 2003. “Ras Proteins: Different Signals from Different Locations.” Nature Reviews. Molecular Cell Biology 4 (5): 373–84.

27. Herrmann, C., G. Horn, M. Spaargaren, and A. Wittinghofer. 1996. “Differential Interaction of the Ras Family GTP-Binding Proteins H-Ras, Rap1A, and R-Ras with the Putative Effector Molecules Raf Kinase and Ral-Guanine Nucleotide Exchange Factor.” The Journal of Biological Chemistry 271 (12): 6794–6800.

28. Itoh, M., C. M. Nelson, C. A. Myers, and M. J. Bissell. 2007. “Rap1 Integrates Tissue Polarity, Lumen Formation, and Tumorigenic Potential in Human Breast Epithelial Cells.” Cancer Research 67 (10): 4759–66.

29. Jacob, A., J. Jing, J. Lee, P. Schedin, S. M. Gilbert, A. A. Peden, J. R. Junutula, and R. Prekeris. 2013. “Rab40b Regulates Trafficking of MMP2 and MMP9 during Invadopodia Formation and Invasion of Breast Cancer Cells.” Journal of Cell Science 126 (20): 4647–58.

30. Jacob, Abitha, Erik Linklater, Brian A. Bayless, Traci Lyons, and Rytis Prekeris. 2016. “The Role and Regulation of Rab40b-Tks5 Complex during Invadopodia Formation and Cancer Cell Invasion.” Journal of Cell Science 129 (23): 4341–53.

31. Jacob, Abitha, and Rytis Prekeris. 2015. “The Regulation of MMP Targeting to Invadopodia during Cancer Metastasis.” Frontiers in Cell and Developmental Biology 3 (February): 16844.

32. Jeon, Taeck J., Dai-Jen Lee, Sylvain Merlot, Gerald Weeks, and Richard A. Firtel. 2007. “Rap1 Controls Cell Adhesion and Cell Motility through the Regulation of Myosin II.” The Journal of Cell Biology 176 (7): 1021–33.

33. Jura, Natalia, Elizabeth Scotto-Lavino, Aleksander Sobczyk, and Dafna Bar-Sagi. 2006. “Differential Modification of Ras Proteins by Ubiquitination.” Molecular Cell 21 (5): 679–87.

34. Kahana, A., and D. E. Gottschling. 1999. “DOT4 Links Silencing and Cell Growth in Saccharomyces Cerevisiae.” Molecular and Cellular Biology 19 (10): 6608–20.

35. Kamura, Takumi, Katsumi Maenaka, Shuhei Kotoshiba, Masaki Matsumoto, Daisuke Kohda, Ronald C. Conaway, Joan Weliky Conaway, and Keiichi I. Nakayama. 2004. “VHL-Box and SOCS-Box Domains Determine Binding Specificity for Cul2-Rbx1 and Cul5-Rbx2 Modules of Ubiquitin Ligases.” Genes & Development 18 (24): 3055–65.

37. Kawabe, Hiroshi, Antje Neeb, Kalina Dimova, Samuel M. Young Jr, Michiko Takeda, Shutaro Katsurabayashi, Miso Mitkovski, et al. 2010. “Regulation of Rap2A by the Ubiquitin Ligase Nedd4-1 Controls Neurite Development.” Neuron 65 (3): 358–72.

38. Kile, Benjamin T., Brenda A. Schulman, Warren S. Alexander, Nicos A. Nicola, Helene M. E. Martin, and Douglas J. Hilton. 2002. “The SOCS Box: A Tale of Destruction and Degradation.” Trends in Biochemical Sciences 27 (5): 235–41.

39. Komander, David, and Michael Rape. 2012. “The Ubiquitin Code.” Annual Review of Biochemistry 81 (1): 203–29.

40. Kooistra, Matthijs R. H., Nadia Dubé, and Johannes L. Bos. 2007. “Rap1: A Key Regulator in Cell-Cell Junction Formation.” Journal of Cell Science 120 (Pt 1): 17–22.

41. Lawson, Campbell D., and Anne J. Ridley. 2018. “Rho GTPase Signaling Complexes in Cell Migration and Invasion.” The Journal of Cell Biology 217 (2): 447–57.

42. Lee, Rebecca Hui Kwan, Hidekazu Iioka, Masato Ohashi, Shun-Ichiro Iemura, Tohru Natsume, and Noriyuki Kinoshita. 2007. “XRab40 and XCullin5 Form a Ubiquitin Ligase Complex Essential for the Noncanonical Wnt Pathway.” The EMBO Journal 26 (15): 3592–3606.

43. Linklater, Erik S., Emily D. Duncan, Ke-Jun Han, Algirdas Kaupinis, Mindaugas Valius, Traci R. Lyons, and Rytis Prekeris. 2021. “Rab40-Cullin5 Complex Regulates EPLIN and Actin Cytoskeleton Dynamics during Cell Migration.” The Journal of Cell Biology 220 (7). https://doi.org/10.1083/jcb.202008060.

44. Linossi, Edmond M., and Sandra E. Nicholson. 2012. “The SOCS Box-Adapting Proteins for Ubiquitination and Proteasomal Degradation.” IUBMB Life 64 (4): 316–23.

45. Liu, Chang, Maho Takahashi, Yanping Li, Tara J. Dillon, Stefanie Kaech, and Philip J. S. Stork. 2010. “The Interaction of Epac1 and Ran Promotes Rap1 Activation at the Nuclear Envelope.” Molecular and Cellular Biology 30 (16): 3956–69.

46. Machida, Noriko, Masato Umikawa, Kimiko Takei, Nariko Sakima, Bat-Erdene Myagmar, Kiyohito Taira, Hiroshi Uezato, Yoshihide Ogawa, and Ken-Ichi Kariya. 2004. “Mitogen-Activated Protein Kinase Kinase Kinase Kinase 4 as a Putative Effector of Rap2 to Activate the c-Jun N-Terminal Kinase.” The Journal of Biological Chemistry 279 (16): 15711–14.

47. McLeod, Sarah J., Anson H. Y. Li, Rosaline L. Lee, Anita E. Burgess, and Michael R. Gold. 2002. “The Rap GTPases Regulate B Cell Migration Toward the Chemokine Stromal Cell-Derived Factor-1 (CXCL12): Potential Role for Rap2 in Promoting B Cell Migration.” The Journal of Immunology 169 (3): 1365–71.

48. McLeod, Sarah J., Andrew J. Shum, Rosaline L. Lee, Fumio Takei, and Michael R. Gold. 2004. “The Rap GTPases Regulate Integrin-Mediated Adhesion, Cell Spreading, Actin Polymerization, and Pyk2 Tyrosine Phosphorylation in B Lymphocytes.” The Journal of Biological Chemistry 279 (13): 12009–19.

49. Meng, Zhipeng, Yunjiang Qiu, Kimberly C. Lin, Aditya Kumar, Jesse K. Placone, Cao Fang, Kuei-Chun Wang, et al. 2018. “RAP2 Mediates Mechanoresponses of the Hippo Pathway.” Nature, August, 1–31.

50. Moreau, Hélène D., Carles Blanch-Mercader, Rafaele Attia, Mathieu Maurin, Zahraa Alraies, Doriane Sanséau, Odile Malbec, et al. 2019. “Macropinocytosis Overcomes Directional Bias in Dendritic Cells Due to Hydraulic Resistance and Facilitates Space Exploration.” Developmental Cell 49 (2): 171–188.e5.

51. Mukhopadhyay, Debdyuti, and Howard Riezman. 2007. “Proteasome-Independent Functions of Ubiquitin in Endocytosis and Signaling.” Science 315 (5809): 201–5.

52. Murphy, Danielle A., and Sara A. Courtneidge. 2011. “The ‘ins’ and ‘Outs’ of Podosomes and Invadopodia: Characteristics, Formation and Function.” Nature Publishing Group 12 (7): 413–26.

53. Myagmar, Bat-Erdene, Masato Umikawa, Tsuyoshi Asato, Kiyohito Taira, Minoru Oshiro, Asako Hino, Kimiko Takei, Hiroshi Uezato, and Ken-Ichi Kariya. 2005. “PARG1, a Protein-Tyrosine Phosphatase-Associated RhoGAP, as a Putative Rap2 Effector.” Biochemical and Biophysical Research Communications 329 (3): 1046–52.

54. Nancy, V., R. M. Wolthuis, M. F. de Tand, I. Janoueix-Lerosey, J. L. Bos, and J. de Gunzburg. 1999. “Identification and Characterization of Potential Effector Molecules of the Ras-Related GTPase Rap2.” The Journal of Biological Chemistry 274 (13): 8737–45.

55. Nethe, Micha, and Peter L. Hordijk. 2010. “The Role of Ubiquitylation and Degradation in RhoGTPase Signalling.” Journal of Cell Science 123 (23): 4011–18.

56. Nonaka, Hideo, Kimiko Takei, Masato Umikawa, Minoru Oshiro, Kouichi Kuninaka, Maitsetseg Bayarjargal, Tsuyoshi Asato, et al. 2008. “MINK Is a Rap2 Effector for Phosphorylation of the Postsynaptic Scaffold Protein TANC1.” Biochemical and Biophysical Research Communications 377 (2): 573–78.

57. Ohba, Yusuke, Kazuo Kurokawa, and Michiyuki Matsuda. 2003. “Mechanism of the Spatio-Temporal Regulation of Ras and Rap1.” The EMBO Journal 22 (4): 859– 69.

58. Okumura, Fumihiko, Akiko Joo-Okumura, Kunio Nakatsukasa, and Takumi Kamura. 2016. “The Role of Cullin 5-Containing Ubiquitin Ligases.” Cell Division, March, 1–16.

59. Parsons, J. Thomas, Alan Rick Horwitz, and Martin A. Schwartz. 2010. “Cell Adhesion: Integrating Cytoskeletal Dynamics and Cellular Tension.” Nature Reviews. Molecular Cell Biology 11 (9): 633–43.

60. Peden, Andrew A., Eric Schonteich, John Chun, Jagath R. Junutula, Richard H. Scheller, and Rytis Prekeris. 2004. “The RCP–Rab11 Complex Regulates Endocytic Protein Sorting.” Molecular Biology of the Cell 15 (8): 3530–41.

61. Petroski, Matthew D., and Raymond J. Deshaies. 2005. “Function and Regulation of Cullin–RING Ubiquitin Ligases.” Nature Reviews. Molecular Cell Biology 6 (1): 9– 20.

62. Pickart, Cecile M., and David Fushman. 2004. “Polyubiquitin Chains: Polymeric Protein Signals.” Current Opinion in Chemical Biology 8 (6): 610–16.

63. Piper, Robert C., Ivan Dikic, and Gergely L. Lukacs. 2014. “Ubiquitin-Dependent Sorting in Endocytosis.” Cold Spring Harbor Perspectives in Biology 6 (1). https://doi.org/10.1101/cshperspect.a016808.

64. Pizon, V., M. Desjardins, C. Bucci, R. G. Parton, and M. Zerial. 1994. “Association of Rap1a and Rap1b Proteins with Late Endocytic/Phagocytic Compartments and Rap2a with the Golgi Complex.” Journal of Cell Science 107 ( Pt 6) (June): 1661–70.

65. Pollard, Thomas D., and Gary G. Borisy. 2003. “Cellular Motility Driven by Assembly and Disassembly of Actin Filaments.” Cell 112 (4): 453–65.

66. Prekeris, R., J. Klumperman, Y. A. Chen, and R. H. Scheller. 1998. “Syntaxin 13 Mediates Cycling of Plasma Membrane Proteins via Tubulovesicular Recycling Endosomes.” The Journal of Cell Biology 143 (4): 957–71.

67. Rebstein, P. J., J. Cardelli, G. Weeks, and G. B. Spiegelman. 1997. “Mutational Analysis of the Role of Rap1 in Regulating Cytoskeletal Function in Dictyostelium.” Experimental Cell Research 231 (2): 276–83.

68. Rebstein, P. J., G. Weeks, and G. B. Spiegelman. 1993. “Altered Morphology of Vegetative Amoebae Induced by Increased Expression of the Dictyostelium Discoideum Ras-Related Gene Rap1.” Developmental Genetics 14 (5): 347–55.

69. Reiner, David J., and Erik A. Lundquist. 2018. “Small GTPases.” WormBook: The Online Review of C. Elegans Biology 2018 (August): 1–65.

70. Ridley, Anne J., Martin A. Schwartz, Keith Burridge, Richard A. Firtel, Mark H. Ginsberg, Gary Borisy, J. Thomas Parsons, and Alan Rick Horwitz. 2003. “Cell Migration: Integrating Signals from Front to Back.” Science 302 (5651): 1704–9.

71. Sadok, Amine, and Chris J. Marshall. 2014. “Rho GTPases: Masters of Cell Migration.” Small GTPases 5 (June): e29710.

72. Sasaki, Atsuo T., Arkaitz Carracedo, Jason W. Locasale, Dimitrios Anastasiou, Koh Takeuchi, Emily Rose Kahoud, Sasson Haviv, John M. Asara, Pier Paolo Pandolfi, and Lewis C. Cantley. 2011. “Ubiquitination of K-Ras Enhances Activation and Facilitates Binding to Select Downstream Effectors.” Science Signaling 4 (163): ra13.

73. Schwartz, S. L., C. Cao, O. Pylypenko, A. Rak, and A. Wandinger-Ness. 2008. “Rab GTPases at a Glance.” Journal of Cell Science 121 (2): 246–246.

74. Shinde, Swapnil Rohidas, and Subbareddy Maddika. 2018. “Post Translational Modifications of Rab GTPases.” Small GTPases 9 (1–2): 49–56.

75. Spaargaren, M., and J. R. Bischoff. 1994. “Identification of the Guanine Nucleotide Dissociation Stimulator for Ral as a Putative Effector Molecule of R-Ras, H-Ras, K-Ras, and Rap.” Proceedings of the National Academy of Sciences 91 (26): 12609–13.

76. Stenmark, Harald. 2009. “Rab GTPases as Coordinators of Vesicle Traffic.” Nature Publishing Group 10 (8): 513–25.

77. Stork, Philip J. S. 2003. “Does Rap1 Deserve a Bad Rap?” Trends in Biochemical Sciences 28 (5): 267–75.

78. Swatek, Kirby N., and David Komander. 2016. “Ubiquitin Modifications.” Nature Publishing Group, March, 1–24.

79. Taira, Kiyohito, Masato Umikawa, Kimiko Takei, Bat-Erdene Myagmar, Manabu Shinzato, Noriko Machida, Hiroshi Uezato, Shigeo Nonaka, and Ken-Ichi Kariya. 2004. “The Traf2-and Nck-Interacting Kinase as a Putative Effector of Rap2 to Regulate Actin Cytoskeleton.” The Journal of Biological Chemistry 279 (47): 49488–96.

80. Thurman, Ryan, Edhriz Siraliev-Perez, and Sharon L. Campbell. 2020. “RAS Ubiquitylation Modulates Effector Interactions.” Small GTPases 11 (3): 180–85.

81. Vega, Michelle de la, James F. Burrows, and James A. Johnston. 2014. “Ubiquitination.” Small GTPases 2 (4): 192–201.

82. Vicente-Manzanares, M. 2005. “Cell Migration at a Glance.” Journal of Cell Science 118 (21): 4917–19.

83. Wang, Lei, Bingxin Zhu, Shiquan Wang, Yuxuan Wu, Wenjian Zhan, Shao Xie, Hengliang Shi, and Rutong Yu. 2017. “Regulation of Glioma Migration and Invasion via Modification of Rap2a Activity by the Ubiquitin Ligase Nedd4-1.” Oncology Reports 37 (5): 2565–74.

84. Warner, Harry, Beverley J. Wilson, and Patrick T. Caswell. 2019. “ScienceDirect Control of Adhesion and Protrusion in Cell Migration by Rho GTPases.” Current Opinion in Cell Biology 56 (February): 64–70.

85. Wennerberg, Krister, Kent L. Rossman, and Channing J. Der. 2005. “The Ras Superfamily at a Glance.” Journal of Cell Science 118 (Pt 5): 843–46.

86. Wiedenmann, Jorg, Sergey Ivanchenko, Franz Oswald, Florian Schmitt, Carlheinz Rocker, Anya Salih, Klaus-Dieter Spindler, and G. Ulrich Nienhaus. 2004. “EosFP, a Fluorescent Marker Protein with UV-Inducible Green-to-Red Fluorescence Conversion.” Proceedings of the National Academy of Sciences 101 (45): 15905–10.

87. Willenborg, Carly, Jian Jing, Christine Wu, Hugo Matern, Jerome Schaack, Jemima Burden, and Rytis Prekeris. 2011. “Interaction between FIP5 and SNX18 Regulates Epithelial Lumen Formation.” The Journal of Cell Biology 195 (1): 71– 86.

88. Xu, Peng, Hannah M. Hankins, Chris MacDonald, Samuel J. Erlinger, Meredith N. Frazier, Nicholas S. Diab, Robert C. Piper, Lauren P. Jackson, Jason A. MacGurn, and Todd R. Graham. 2017. “COPI Mediates Recycling of an Exocytic SNARE by Recognition of a Ubiquitin Sorting Signal.” *ELife* 6 (October). https://doi.org/10.7554/eLife.28342.

89. Yatsu, A., H. Shimada, N. Ohbayashi, and M. Fukuda. 2015. “Rab40C Is a Novel Varp-Binding Protein That Promotes Proteasomal Degradation of Varp in Melanocytes.” Biology Open 4 (3): 267–75.

90. Yin, Guowei, Jerry Zhang, Vinay Nair, Vinh Truong, Angelo Chaia, Johnny Petela, Joseph Harrison, Alemayehu A. Gorfe, and Sharon L. Campbell. 2020. “KRAS Ubiquitination at Lysine 104 Retains Exchange Factor Regulation by Dynamically Modulating the Conformation of the Interface.” IScience 23 (9): 101448.

91. Zerial, M., and H. McBride. 2001. “Rab Proteins as Membrane Organizers.” Nature Reviews. Molecular Cell Biology 2 (2): 107–17.

92. Zhang, Mingshu, Hao Chang, Yongdeng Zhang, Junwei Yu, Lijie Wu, Wei Ji, Juanjuan Chen, et al. 2012. “Rational Design of True Monomeric and Bright Photoactivatable Fluorescent Proteins.” Nature Methods 9 (7): 727–29.

